# Transcriptome-wide sites of collided ribosomes reveal principles of translational pausing

**DOI:** 10.1101/710061

**Authors:** Alaaddin Bulak Arpat, Angélica Liechti, Mara De Matos, René Dreos, Peggy Janich, David Gatfield

**Affiliations:** Center for Integrative Genomics, University of Lausanne, 1015 Lausanne, Switzerland; Krebsliga Schweiz, 3001 Bern, Switzerland

## Abstract

Translation initiation is the major regulatory step defining the rate of protein production from an mRNA. Meanwhile, the impact of non-uniform ribosomal elongation rates is largely unknown. Using a modified ribosome profiling protocol based on footprints from two closely packed ribosomes (disomes), we have mapped ribosomal collisions transcriptome-wide in mouse liver. We uncover that the stacking of an elongating onto a paused ribosome occurs frequently and scales with translation rate, trapping ∼10% of translating ribosomes in the disome state. A distinct class of pause sites, independent of translation rate, is indicative of deterministic pausing signals. Pause site association with specific amino acids, peptide motifs and nascent polypeptide structure, is suggestive of programmed pausing as a widespread mechanism associated with protein folding. Evolutionary conservation at disome sites indicates functional relevance of translational pausing. Collectively, our disome profiling approach allows unique insights into gene regulation occurring at the step of translation elongation.

## Introduction

The translation of messenger RNA (mRNA) to protein is a central step in gene expression. Our knowledge of this process has exploded since the emergence of ribosome profiling (Ingolia et al., 2009), a technique based on the high-throughput sequencing of the ∼30 nt mRNA footprints that are buried inside the translating ribosome and thus protected from the nuclease treatment that is used to digest the mRNA regions that are not occupied by ribosomes. A plethora of studies have built on the quantitative, transcriptome-wide and nucleotide-resolved information that ribosome profiling provides to gain insight into a variety of aspects of protein biosynthesis (see Ingolia et al. (2019) for a recent review). This includes, among others, the annotation of translated mRNA regions, the study of differential translation across various biological and experimental paradigms, the characterization of intermediate states of the translating ribosome, the subcellular compartmentalization of protein biosynthesis, or functional differences in translational capacity within a heterogeneous cellular ribosome population.

Most available ribosome profiling data is consistent with the longstanding notion that of the four distinct phases defining translation, i.e. initiation, elongation, termination and ribosome recycling, it is the first step – the commitment of the ribosome to initiate – that is rate-limiting for the overall process in eukaroytes (Hinnebusch, 2014). It is thus assumed that the quantity of elongating ribosome footprints (i.e., the species mainly captured by conventional ribosome profiling methodology) is proportional to initiation rate and to overall protein biosynthesis. Elongating ribosome footprints typically distribute in a distinctly non-uniform fashion across a given protein coding sequence (CDS), which has been attributed to variations in ribosome decoding speed and dwell times (Ingolia et al., 2011). Integrating footprint reads across the entire CDS is thought to correct for local variation in footprint density and to allow for accurate estimates of relative translation efficiencies per gene (TEs, calculated as CDS-mapping footprint reads normalized to RNA abundance) in a transcriptome-wide fashion. Nevertheless, a possible influence of local footprint variation on overall translation speed of an mRNA has been suggested early on (Dana and Tuller, 2012) and, in general, how to interpret apparent local differences in footprint densities is not fully resolved. Of note, it remains an intrinsic limit of the technique that it delivers static snap-shots of ribosome occupancy rather than dynamic data of the translation process. Therefore, and somewhat paradoxically, in the two extreme, hypothetical scenarios of one transcript whose elongating ribosomes are translationally paused (resulting in low or no protein biosynthesis), and of another transcript that shows strong, productive flux of elongating ribosomes (resulting in high protein biosynthesis), the actual footprint snap-shots that would be seen in ribosome profiling may actually be indistinguishable. To discern such cases, a dedicated genome-wide method for the direct detection of ribosomal pausing would be crucial. In yeast, specific footprint size classes associated with stalled ribosomes have been described (Guydosh and Green, 2014; Diament et al., 2018).

Historically, early evidence for paused elongation – leading to the subsequent stacking of upstream elongating ribosomes onto the paused one – has come from *in vitro* translation reactions (Wolin and Walter, 1988). For a limited number of prominent cases, pausing has since then been shown to be functionally important for proper protein localization to membranes (Yanagitani et al., 2011; Mariappan et al., 2010), to serve as a mechanism for start codon selection (Ivanov et al., 2018), and to regulate the extent of productive full-length protein biosynthesis (Yordanova et al., 2018). It is tempting to extrapolate from such individual examples to general roles for elongation pausing that cells could employ to control protein biosynthesis post-initiation. At the other end of the spectrum, hard elongation stalls caused by various obstacles to processive translation (including defective mRNAs or specific amino acid motifs in the nascent peptide) require resolution by the ribosome-associated quality control pathway (RQC), and the mechanisms through which such terminally stalled ribosomes are sensed and handled is a highly active field of current research (reviewed in Joazeiro (2019)).

An early ribosome profiling study in mouse embryonic stem cells (mESCs) already addressed the question of how to extract potential pause sites from footprint data (Ingolia et al., 2011), which resulted in the identification of thousands of alleged pauses within CDS sequences as well as an enrichment at termination codons. In combination with quantitative modelling approaches, subsequent studies have identified parameters that can impinge on local translation speed and pausing (reviewed in Schuller and Green (2018)). Among these are, notably, specific amino acids (Charneski and Hurst, 2013), codon pairs (Gamble et al., 2016), tRNA availability (Darnell et al., 2018; Guydosh and Green, 2014), RNA secondary structures (Zhang et al., 2017; Pop et al., 2014), or the folding (Doring et al., 2017) and exit tunnel interactions (Dao Duc and Song, 2018; Charneski and Hurst, 2013) of the nascent peptide. However, to what extent translational pausing occurs *in vivo* in a mammalian system, which characteristics these pause sites have, and whether they are functionally relevant is still poorly understood.

Here, we have applied a modified ribosome profiling strategy to a mammalian organ, mouse liver, in order to directly reveal in a transcriptome-wide fashion the sites where two ribosomes collide. The characteristics associated with these ∼60 nt “disome footprints” are consistent with the expectations for collision events of an upstream elongating ribosome onto a downstream, paused ribosome. Through the use of synthetic footprint spike-ins, we estimated the quantitative relationship between disome and monosome footprints. Deep analysis of the disome sites allowed identifying features predictive of ribosome pausing, including sequence features of the mRNA and structural features of the nascent polypeptide. Finally, we address the question of the functional relevance of translational pause sites.

## Results

### Disome footprint sequencing allows transcriptome-wide mapping of ribosomal collisions

A critical step in ribosome profiling is the quantitative, nuclease-mediated conversion of polysomes to individual monosomes, from which protected mRNA footprints of ∼30 nt can be purified and converted to sequenceable libraries. During the setup of this technique from mouse liver polysomal extracts for a previous study (Janich et al., 2015), we used Northern blots to monitor the efficiency of RNase I-mediated footprint generation. Radioactively labelled short oligonucleotide probes antisense to the protein coding sequences (CDS) of the highly abundant *Albumin (Alb)* and *Major urinary protein 7 (Mup7)* mRNAs indeed showed the expected ∼30 nt monosome footprints (Fig. 1A, B). Moreover, several of the probes also detected distinct higher-order bands whose estimated sizes were compatible with those expected for multiples of monosome footprints (i.e. ∼60 nt, ∼90 nt etc.). These additional bands were particularly prominent for the probes designed to anneal to the transcripts just downstream of where the signal peptide (SP) was encoded (see monosome footprint and two higher-order bands for probe Alb_71−101_ in Fig. 1A; see monosome footprint and >5 higher-order bands for probe Mup7_58−81_ in Fig. 1B). We initially interpreted the presence of these bands as an indication of suboptimal conditions during nuclease treatment, leading to an incomplete collapse of polysomes to monosomes. We hence tested other nuclease digestion conditions. However, neither changes in temperature or detergent concentrations during extract preparation and nuclease treatment (Supplemental Fig. S1A), nor higher RNase I activity (Supplemental Fig. S1B), nor a different nuclease altogether, micrococcal nuclease (Supplemental Fig. S1C), were able to collapse the higher-order bands quantitatively to the size of the monosome footprint. We thus speculated that the higher-order footprints reflected a distinct, relatively stable state of translating ribosomes, possibly resulting from two (disome), three (trisome) or, in the case of the bands seen for Mup7_58−81_, even higher numbers of ribosomes whose dense stacking rendered the mRNA inaccessible to nucleases. This scenario was reminiscent of the ribosomal pausing and stacking described in the 1980s for *in vitro* translated preprolactin mRNA (Wolin and Walter, 1988). Here, a major translation stall site at codon 75 (a GGC glycine codon), which led to the queuing of subsequent incoming ribosomes, was related to the recruitment of the signal recognition particle (SRP) to the SP. We wished to determine whether our higher-order footprints reflected a similar phenomenon and whether they would allow us to detect ribosomal pause and collision sites transcriptome-wide and *in vivo*. We thus selected a subset of samples from our previously collected mouse liver time series (Janich et al., 2015), corresponding to three timepoints at the beginning of the daily light (*Zeitgeber* Times ZT0 and ZT2) and dark phases (ZT12), and subjected them to ribosome profiling for both the ∼30 nt monosome footprints and the ∼60 nt alleged disome footprints; we also determined RNA abundances from the same samples by RNA-seq (Fig. 1C). Libraries were sequenced sufficiently deeply to obtain >10^8^ cDNA-mapping reads per footprint species (Fig. 1D, Supplemental Fig. S2A and Supplemental Table S1). Monosome footprints showed the expected length and mapping features, i.e. the majority was 29-30 nt in size (Fig. 1E) and they were enriched on CDS and depleted from untranslated regions (UTRs) (Supplemental Fig. S2B). This depletion was considerably stronger for 3′ UTRs than for 5′ UTRs, which was expected given that 5′ UTRs are known to harbour considerable translational activity on upstream open reading frames (uORFs). Disome footprint lengths showed a broader distribution with two distinct populations at 59-60 nt and 62-63 nt (Fig. 1E) that resembled the bimodal pattern that has been reported in yeast as well (Guydosh and Green, 2014). The mapping to transcript regions was similar to that of monosome footprints, albeit with a more pronounced depletion from 5′ UTRs (Supplemental Fig. S2B). As the median uORF length in mice is <40 nt (Johnstone et al., 2016), it is likely that many uORFs are simply too short to accommodate two translating ribosomes simultaneously. Reduced levels in 5′ UTR disome footprints were thus fully compatible with the hypothesis that they reflected ribosomal collisions.

**Figure 1.**
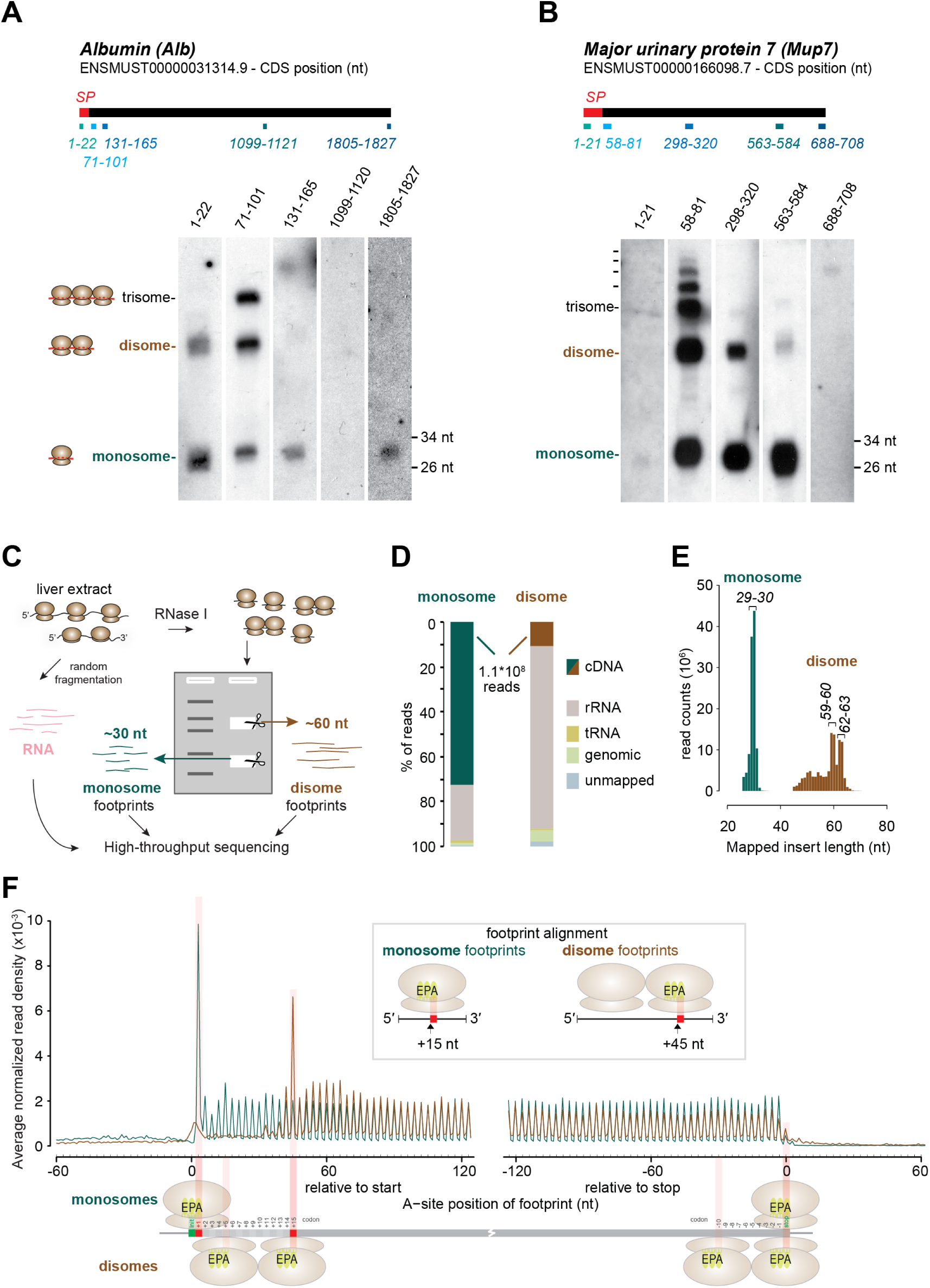
Sequencing of disome footprints identifies transcriptome-wide ribosomal collisions. **(A and B)** Northern blot analysis of RNase I-treated mouse liver extracts probed with radioactively labelled short oligonucleotides antisense to different regions of *Albumin (Alb)* **(A)** and *Major urinary protein 7 (Mup7)* mRNAs **(B)**. Expected footprint sizes for monosomes, disomes and trisomes are shown to the left of blots. Positions of probes (nt) relative to the annotated CDS start sites on the indicated transcripts, ENSMUST00000031314.9 and ENSMUST00000166098.7, are shown above each lane and also depicted as blue boxes below the black bars representing the CDS regions (top). Red boxes at the 5′ end indicate the coding region for signal peptide (SP) on each transcript. **(C)** Graphical representation of experimental setup for sequencing of 60 nt disome footprints. **(D)** Bar-plots of percentages of reads from monosome and disome libraries that were mapped to different sequence types: rRNA (gray), tRNA (golden), genomic (green) and cDNA/mRNA (teal for monosomes and brick red for disomes). Percentages of unmapped reads are shown in blue. **(E)** Histogram of insert size (nt) for reads that mapped to cDNA/mRNA sequences (monosomes: teal, disomes: brick red). A single mode for monosomes (29-30 nt) and two modes for disomes (59-60 and 62-63 nt) are labeled above histograms. **(F)** Density distribution of footprint reads within 120 nt from the start or −120 nt from the stop codons reveals 3-nt periodicity of monosome and disome footprints within coding sequences. The meta-transcript analysis quantified the mean of per-transcript normalized number of reads (monosomes: teal, disomes: brick red) at each nucleotide based on the A-site prediction (15 nt and 45 nt downstream of the 5′ end of monosome and disome footprints, respectively). Only transcripts from single protein isoform genes with totalRNA-RPKM >5, CDS >400 nt, and UTRs of >180 nt (N = 4994) were used. Positions on footprints, corresponding to the predicted E-, P-, and A- sites of ribosomes that presumably protected the corresponding footprints, are shown in graphical depictions. Start/stop codons are highlighted as green boxes on a representative transcript (at the bottom).

We next analyzed frame preference and distribution along the CDS for the two footprint species. To this end, we mapped the predicted position of the ribosomal aminoacyl-tRNA acceptor site (A-site) codon of each monosome footprint (i.e. nucleotides 15-17 for 29-30 nt footprints, see Janich et al. (2015)) onto the meta-transcriptome. We observed the characteristic 3-nt periodicity of ribosome footprints across coding sequences, starting at the +1 codon relative to the initiation site (note that initiating ribosomes carry the first tRNA already in their P-site, and the A-site is thus placed over the +1 codon) and ending at the termination codon (Fig. 1F). Moreover, the profile showed previously reported features, including elevated and reduced ribosome densities at the start and stop codons, respectively, as well as increased occupancy of the +5 codon, which has been interpreted to reflect a pause occurring between initiation and elongation commitment (Han et al., 2014).

For an equivalent analysis of the disome footprints, we aligned them to the CDS according to their +45 nt-position, corresponding to the alleged A-site of the leading ribosome (Fig. 1F; see Supplemental Fig. S3A for a complementary graph with an alignment to the +15 position, i.e. the A-site of the lagging ribosome). Like the monosome footprints, disome footprints also showed transcriptome-wide 3-nt periodicity. At the CDS 3′ end, footprint coverage ended at the position expected when the disome’s leading ribosome would occupy the termination codon. The globally highest disome footprint abundance was found close to the CDS 5′ end, at a position corresponding to a disome formed from lagging and leading ribosomes on the +5 and +15 codons, respectively (Fig. 1F; Supplemental Fig. S3A). Upstream of this position, i.e. on the first few codons post-initiation, there was a distinct depletion of disomes, whereas further downstream the distribution was overall rather uniform with some 5′-to-3′ decrease over the remainder of the CDS. Of note, we also remarked a small, local increase in footprints that may correspond to a leading ribosome on the initiation codon and a lagging ribosome at the −10 codon in the annotated 5′ UTR (marked with blue arrow in Supplemental Fig. S3A), presumably reflecting translated uORF codons.

Taken together, the above findings were consistent with the hypothesis that the ∼60 nt higher-order bands indeed represented footprints originating from translated mRNA that was protected by two adjacent, and possibly collided, ribosomes. Moreover, it would appear that transcriptome-wide these alleged collision events could occur at most CDS positions, although the likelihood of stacking onto a downstream ribosome would seem reduced for the first few codons post-initiation. Sterical constraints to initiation – for example extra space that may be required to allow the formation of the initiation complex with its associated initiation factors – have been proposed to play a role in a similar phenomenon in yeast (Guydosh and Green, 2014).

*Disome occurrence is locally favored by signal peptides, and globally by high translation efficiency* The SRP-dependent ribosome pausing and stacking described for preprolactin mRNA (Wolin and Walter, 1988) suggested that transcripts encoding signal peptide-containing proteins (termed SP transcripts in the following) could serve as a positive control to validate that our approach was capturing similar events. We thus assessed footprint densities along the CDS for SP transcripts (N=713) vs. non-SP transcripts (N=4743). SP transcripts showed a striking build-up of disome footprints towards the 5′ end of the CDS that extended virtually to the position (∼75 amino acids) Wolin & Walter had described for preprolactin mRNA, whereas downstream of this region disome read densities were reduced (Fig. 2A-B). Importantly, these features were specific to disomes, as they were absent from monosome footprint data (Fig. 2C-D). We concluded that the profiling of disome footprints was able to capture the same translational pausing and stacking events that had previously been described (Wolin and Walter, 1988).

**Figure 2.**
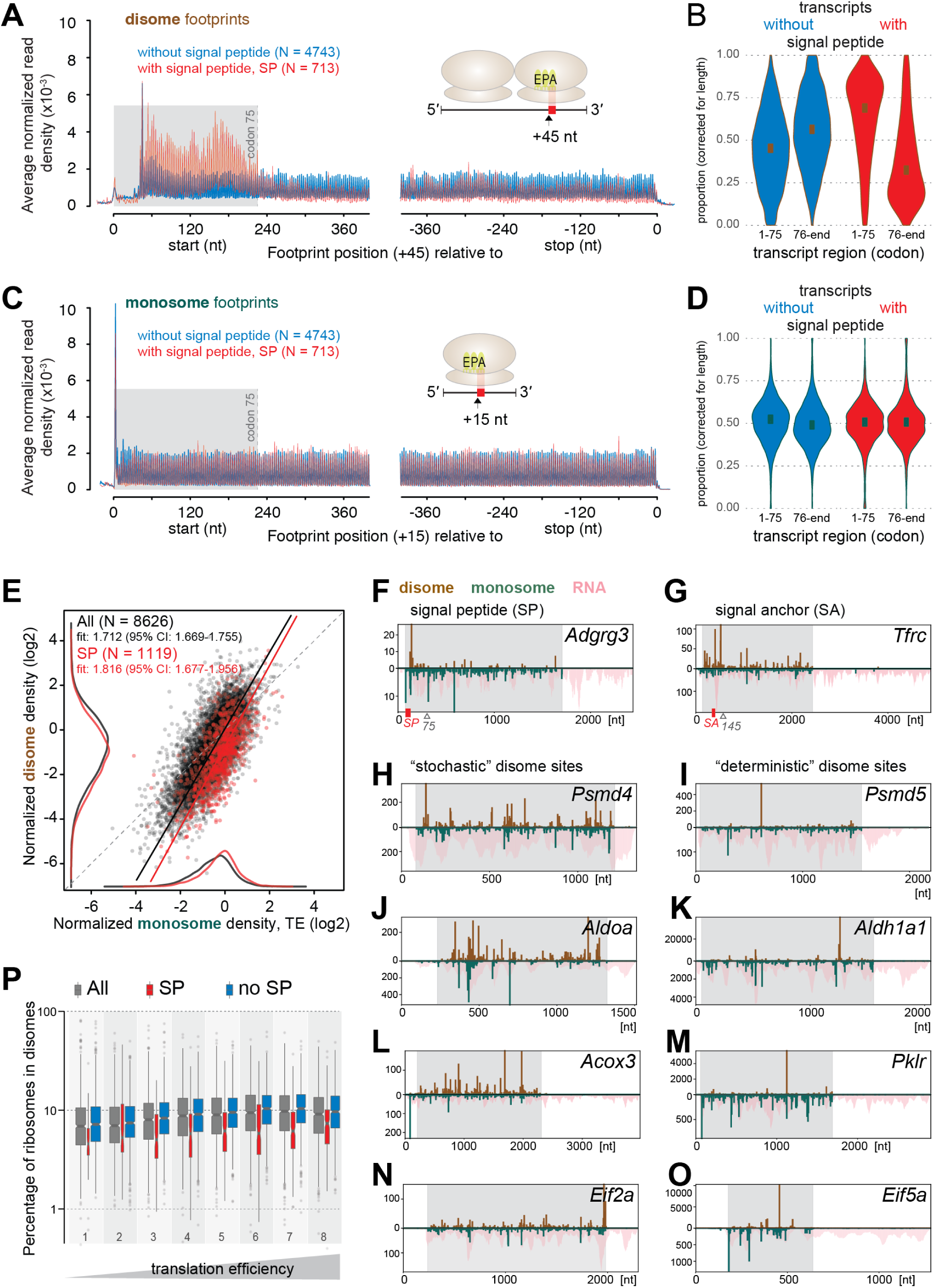
Occurrence of disomes is associated locally with signal peptides, and globally with high volumes of translation. **(A and C)** Density distribution of disome footprints identify signal peptide (SP)-related pausing events. Meta-transcript analysis (see Fig. 1F) quantified the mean normalized footprint densities of disomes **(A)** and monosomes **(C)** within 400 nt from the start or −400 nt from the stop codons of transcripts confirmed or predicted to code for SP (red, N = 713) or not (blue, N = 4743). **(B and D)** Violin-plots showing the probability densities of length normalized proportions of footprints within the first 75 codons and the rest of CDS from transcripts with a signal peptide (red, N = 713) or without one (blue, N = 4743) for disomes **(B)** and monosomes **(D)**. **(E)** Scatter-plot illustrating the relationship between per-gene normalized densities of disome and monosome footprints. The subset of all genes included in the analysis (N = 8626; black and red dots together) that code a signal peptide (SP) (N = 1119) are denoted by red dots. Kernel density estimates are plotted on the margins (monosome on x-, disome on y-axis) for datasets of all genes (black) and SP coding genes (red) (without an axis of ordinates). Deming regression (errors-in-variables model to account for errors in both monosome and disome footprint estimates) lines are shown for all genes (black) and the SP coding subset (red). Regression slopes and their 95% confidence intervals (CI) are given in the top-left legends. Dashed gray line indicates the 1-to-1 slope. **(F - O)** Distribution of normalized counts of monosome and disome footprints (per nt) along transcripts of representative genes reveals presence of stochastic and deterministic sites. Upward y-axis of the bar-plots show the normalized read counts for disomes (brick red), while downward axis was used for monosomes (teal) and totalRNA (pink, pile-up). Transcript coordinates (nt) are shown on x-axis; the regions corresponding to respective CDS are shaded in gray. If present, the signal peptide coding regions are indicated with small red boxes along the x-axis. Plots are shown for *Adhesion G protein-coupled receptor G3 (Adgrg3), Transferrin receptor (Tfrc), Proteasome 26S Subunit, Non-ATPase* 4 *(Psmd4)* and *5 (Psmd5), Aldehyde dehydrogenase 1a1 (Aldh1a1), Pyruvate kinase liver and red blood cell (Pklr)* and *Eukaryotic translation initiation factor 5A (Eif5a)* in **(F-O)**, respectively. **(P)** Box-plots illustrating the estimated amounts of ribosomes that were retained in disomes as a percentage of all translating ribosomes for different groups of genes in mouse liver samples. Box-and-whiskers were drawn for all genes, detectably expressed in spike-in experiment (all, gray, N = 7375), subsets that codes for SP (SP, red, N = 892) and not (no SP, blue, N = 6483) and stratified into 8 groups based on the octiles of the translation efficiency (TE) calculated using all genes which had the following right-closed interval boundaries (−5.41, −1.23, −0.77, −0.47, −0.23, −0.04, 0.17, 0.47, 3.17), depicted as increasing TE below the x-axis. Width of each box is proportional to the number of data points it represents.

We next analyzed monosome and disome footprints per gene. Of note, many transcripts that were detectable at the monosome footprint and RNA-seq levels also showed sufficient coverage in the disome data, allowing for robust quantification of both footprint species across a large portion of the expressed genome (N=8626 genes). We first computed for each gene individually the ratio of CDS-mapping footprint to RNA-seq reads. For monosome footprints, this ratio is frequently referred to as “ribosome density” and considered to reflect a transcript’s relative translational efficiency (TE). In an analogous fashion, for the disome footprints this density would correspond to a measure for the extent of ribosomal pausing and stacking. When comparing disome and monosome footprint densities per gene, we made two main observations. First, disome densities were positively correlated with translation efficiencies (Fig. 2E). SP transcripts showed this correlation as well; however, they were globally shifted to lower disome footprint levels, indicating that the high disome occurrence up to codon ∼75 was outweighed by the reduction seen over the remainder of the CDS (Fig. 2B, Supplemental Fig. S3C). Second, the steepness of the fit in the double-log plot in Fig. 2E was considerably greater than 1, indicating a power relation between disome and monosome densitites. Conceivably, increased ribosomal flux on mRNAs was thus associated with an even higher relative increase in ribosomal collisions.

We next assessed whether this relationship between disome and monosome footprint levels was not only observable across different transcripts, but also for a given transcript at different translation efficiencies. To this end, we analyzed mRNAs encoding for ribosomal proteins (RPs), which show prominent, feeding-dependent daily rhythms in TE (Sinturel et al., 2017). Using the two timepoints from our datasets that corresponded to states of low (ZT2) and high (ZT12) ribosomal protein mRNA translation (Janich et al., 2015), our analyses revealed that the increase in disome density on RP transcripts was significantly greater than the ∼2-fold increase in monosome TE between the two timepoints (Supplemental Fig. S3D), consistent with the power relation between disome and monosome footprints. Taken together, these findings suggested that – at least in part – disome footprints were a consequence of ribosomal crowding that could in principle occur during any translation event, but that was favoured under high ribosomal traffic (”traffic jams”, Diament et al. (2018)). In such a case, one might hypothesize that the positions on the CDS where ribosomes collided could have a sizable stochastic component; in addition, local differences in ribosomal dwell times – which are associated with amino acid/codon usage and the size of the amino acid-loaded tRNA pool (Gobet et al., 2020) – would be expected to bias the collision sites as well. In contrast, however, the observation that signal peptides represented general triggers for ribosome stalling and queuing, as well as differences in disome levels across transcripts that were not simply attributable to TE differences (Supplemental Fig. S3E), suggested that beyond the alleged “stochastic” sites, more specific, “deterministic” signals and stall sites existed, too.

In order to gain a sense of whether these different scenarios and classes of disome sites (stochastic/deterministic) truly existed, we first visually inspected various individual transcript examples (Fig. 2F-O; see also Supplemental Fig. S4 for transcript plots stratified by individual ZT libraries). To begin with, we noted that individual SP transcripts indeed exhibited the expected disome patterns. As shown for the case of *Adhesion G protein-coupled receptor G3 (Adgrg3) –* whose annotated signal peptide spans amino acids (aa) 1-18 – disome footprint coverage was elevated in the region up to codon ∼75 and was lower and appeared more randomly distributed over the remainder of the transcript (Fig. 2F). Similarly, *Transferrin receptor (Tfrc)*, which contains an SRP-dependent signal anchor (SA) sequence at aa 68-88 (Zerial et al., 1986), showed elevated disome levels extending until codon ∼145 (Fig. 2G), indicating a direct relationship between the positions of disome buildup and of the signal sequence/SRP interaction. We next examined individual transcripts lacking signal sequences for the presence of the alleged stochastic and deterministic sites. For example, the transcripts coding for two subunits of the 19S regulatory particle of the proteasome, *Psmd4* (Fig. 2H) and *Psmd5* (Fig. 2I), showed distinct patterns of disome distribution that were consistent with our expectations for stochastic and determinitic sites, respectively. *Psmd4* thus showed disome coverage at numerous positions along the CDS. By contrast, a specific, dominant site was apparent for *Psmd5*. Many other transcripts showed such patterns with distinct dominant sites as well, e.g. *Aldehyde dehydrogenase 1a1 (Aldh1a1)* (Fig. 2K), *Pyruvate kinase liver and red blood cell (Pklr)* (Fig. 2M) and *Eukaryotic translation initiation factor 5A (Eif5a)* (Fig. 2O). Dispersed disome patterns similar to *Psmd4*, as well as mixed cases combining broad coverage with specific dominant sites were frequent, too, e.g *Aldolase A (AldoA)* (Fig. 2J), *Acyl-Coenzyme A oxidase 3 (Acox3)* (Fig. 2L) and *Eukaryotic translation initiation factor 2A (Eif2a)* (Fig. 2N). Furthermore, we made the empirical observation that in some cases there was not (e.g. *Aldh1a1*), and in others there was (e.g. *Pklr*), a correspondence between the sites of strong disome and monosome accumulation. Indeed, both scenarios – correlation and anti-correlation – between strong disome and monosome sites appear plausible: on the one hand, extended ribosomal dwell times should lead to the capture of more monosome footprints from slow codons – and since these positions would also represent sites of likely ribosomal collisions, they would be enriched in the disome data as well. On the other hand, however, for sites where collisions are very frequent – to the extent that stacked ribosomes become the rule – one may expect to see an effective loss of these positions in the monosome footprint data. An important consequence of elongating ribosomes getting trapped in disomes is that conventional (i.e., monosome footprint-based) ribosome profiling datasets will inevitably underestimate the number of translating ribosomes per transcript, in particular for mRNAs with high translation efficiency. We wished to quantify this effect. This was, however, not possible from our existing monosome and disome footprint datasets because they originated from independent libraries (Fig. 1C). Consequently, we had no means of normalizing the disome and monosome data relative to each other. We therefore sequenced new libraries from liver samples to which, early in the protocol, we had added defined quantities of synthetic 30mer and 60mer RNA spike-ins (Supplemental Fig. S5A-B), which subsequently allowed for a quantitative realignment of monosome and disome footprint data. This approach revealed that for transcripts with high TE, typically ∼10% of translating ribosomes were trapped in disomes (Fig. 2P). This proportion decreased with decreasing TE and was generally reduced for SP-transcripts, as expected. Of note, this estimates were consistent with the strong disome bands seen in the Northern blots (Fig. 1A-B).

In summary, we concluded that disome formation was a common phenomenon and observable across most of the transcriptome. The association with signal peptides and with high translational flux indicated that disome footprints were indeed resulted from ribosomal collisions between a downstream, slow decoding event and an upstream ribosome that stacked onto the (temporarily) stalled ribosome.

### Disome sites are associated with specific amino acids and codons

We next investigated whether disome sites were associated with mRNA sequence features, in particular with specific codons or amino acids. To this end, we adapted a method developed for the analysis of monosome-based footprint data, termed Ribo-seq Unit Step Transformation (RUST), which calculates observed-to-expected ratios for a given feature at each codon position within a window that encompasses the footprint and surrounding upstream and downstream regions (O′Connor et al., 2016). RUST-based enrichment analyses in O′Connor et al. showed that ribosome footprints had the highest information content (relative entropy, expressed as Kullback-Leibler divergence) on the codons placed within the ribosome decoding center. Moreover, the sequence composition at the 5′ and 3′ termini of the mRNA fragments showed pronounced non-randomness as well, which was, however, not specific to footprints but found in RNA-seq data as well. It was thus concluded that the main contributing factors to footprint frequency at a given mRNA location were, first, the identity of the codons in the A- and P-sites and, second, the sequence-specificity of the enzymes used for library construction (O′Connor et al., 2016).

To be able to apply the RUST pipeline to the disome footprints, we first needed to investigate the origins of their bimodal length distribution (Fig. 1E) and determine which footprint nucleotides likely corresponded to the ribosomal E-, P- and A-sites. To this end, enrichment analyses for codons, conducted individually for the different size classes of disome footprints (Supplemental Fig. S6A), resulted in profiles that resembled the previously described RUST-based enrichment profiles for monosome data (O′Connor et al., 2016). Increased information content was thus apparent at the footprint 5′ and 3′ boundaries, indicative of the aforementioned biases from library construction. Moreover, codon selectivity was consistently seen in the footprint region that would be occupied by the leading ribosome’s decoding center, ∼15 nt upstream of the footprint 3′ end. Notably, no selectivity was visible for the region occupied by the upstream ribosome or at the boundary between the ribosomes. These findings were consistent with the model that the leading ribosome defined the pause site (with preference for specific codons) and an upstream ribosome colliding in a sequence-independent fashion. Importantly, the comparison of the enrichment plots from the different footprint lengths allowed us to propose a likely interpretation for the observed length heterogeneity. The two major populations of 59-60 nt and 62-63 nt thus appeared to correspond to ribosome collisions in which the upstream ribosome stacked onto the stalled ribosome in two distinct states that differed by one codon (Supplemental Fig. S6B). Conceivably, the 1 nt variation (i.e., 59 nt vs. 60 nt and 62 nt vs. 63 nt) corresponded to differences in trimming at the footprint 3′ end. Using this model, we aligned the main populations from the range of disome footprint lengths (i.e., 58-60 nt and 62-63 nt, representing together about two-thirds of all disome footprints) according to the predicted A-site of the paused, leading ribosome (see Supplemental Methods). Applying these corrected A-site predictions to the meta-transcriptome analyses led to an overall improvement of the 3-nt periodic signal of the disome footprints (Supplemental Fig. S3B; compare with Fig. 1F). We used these A-site-corrected footprints for the RUST pipeline.

Enrichment analysis showed marked selectivity for amino acids in the decoding center (P- and A-site) of the disome’s leading ribosome (Fig. 3A, left panel). The magnitude of amino acid preference was considerably greater than that seen for the monosome footprints (Fig. 3A, middle (G) panel), whereas RNA-seq data lacked selectivity beyond the effects attributable to library generation enzymology, as expected (Fig. 3A, right panel). Specific amino acids stood out as preferred ribosome stall sites, irrespective of codon usage. The most striking associations were, notably, the prominent overrepresentation of aspartic acid (D) in both the P- and A-sites (Fig. 3B), the enrichment of isoleucine (I) in the A-site and its depletion from the P-site (Fig. 3C), and the enrichment of glycine (G) in the P-site (Fig. 3D) of paused ribosomes. We next transformed the full amino acid analysis (Supplemental Fig. S7) to a position weight matrix representing the ensemble of positive and negative amino acid associations with disome sites (Fig. 3E). The enrichment of acidic (D, E) and the depletion of certain basic amino acids (K, H) within the decoding center of the leading ribosome suggested that amino acid charge may be a relevant factor for ribosomal pausing. Moreover, we noticed that for certain amino acids, association with disome sites was dependent on codon usage. For example, P-site Asparagin (N) was strongly associated with pause sites only when encoded by AAT, but not by AAC (Fig. 3F); Lysine (K) was depleted at P-sites irrespective of the codon, but the association of A-site Lys was highly dependent on codon usage, showing either depletion (AAA) or enrichment (AAG) (Fig. 3G). Next, we compared the enrichment patterns with those obtained for the two major footprint populations of 59-60 nt or 62-63 nt individually (Supplemental Fig. S8A-B). At the decoding center of the paused ribosome, the weight matrices showed near identical patterns, indicating that the two footprint size classes did not discriminate pausing events of different specificity. We also compared the reproducibility of enrichment patterns. Across the 6 independent biological samples, RUST profiles were near-identical at the decoding center of the paused ribosome (Supplemental Fig. S9, S10), and the completely independent libraries from the spike-in experiment showed highly similar enrichment patterns as well (Supplemental Fig. S8C). Finally, we realized that the observed amino acid signatures showed resemblance with ribosomal dwell times that were recently estimated through modelling of conventional mouse liver ribosome profiling data (Gobet et al., 2020). Indeed, our monosome footprint data, too, showed similar patterns of amino acid enrichment and depletion, though much reduced in magnitude (Supplemental Fig. S7, S10, S11A). We next investigated the association of pause sites with specific amino acid combinations. Strong selectivity with regard to the 400 possible dipeptide motifs was apparent in the P- and A-sites of the leading ribosome (Fig. 3H, left panel). This effect was much weaker and absent for monosome and RNA data, respectively (Fig. 3H, middle and right panels). In the disome data, the enrichment was highest, and independent of codon usage, for dipeptides consisting of the most enriched single amino acids, i.e. Gly-Ile (GI), Asp-Ile (DI) and Gly-Asp (GD) (Fig. 3I-K). By contrast, the pausing of ribosomes at several other dipeptides was strongly dependent on codon usage. In particular the presence of Lysine (K) or Glycine (G) in the A-site of the leading ribosome was associated with codon selectivity (Fig. 3L). For instance, the Asp-Lys (DK) dipeptide was highly associated with disomes when encoded by GATAAG (Fig. 3M; blue trace); notably, with transcriptome-wide 910 cases of disome peaks observed on the 2030 existing GATAAG positions (i.e. 44.8%), it was the 8th most disome-prone dicodon out of the total 3721 (i.e., 61 x 61) possible dicodon combinations (Supplemental Table S2; Fig. 3O). By contrast, when encoded by GACAAA (Fig. 3M; black trace), disomes were observable on no more than 7.8% of sites (272 out of 3529), ranking this dicodon at position 1419. The Gly-Gly (GG) dipeptide represented a similar case (Fig. 3N); of the 16 dicodon combinations, GGAGGA (blue trace) was most strongly enriched (698/2407, i.e. 29.0% of sites showed disome peaks; rank 64), whereas GGCGGC (black trace) showed depletion from disome sites (92/1738, i.e. 5.3% of sites showed disome peaks; rank 2304).

**Figure 3.**
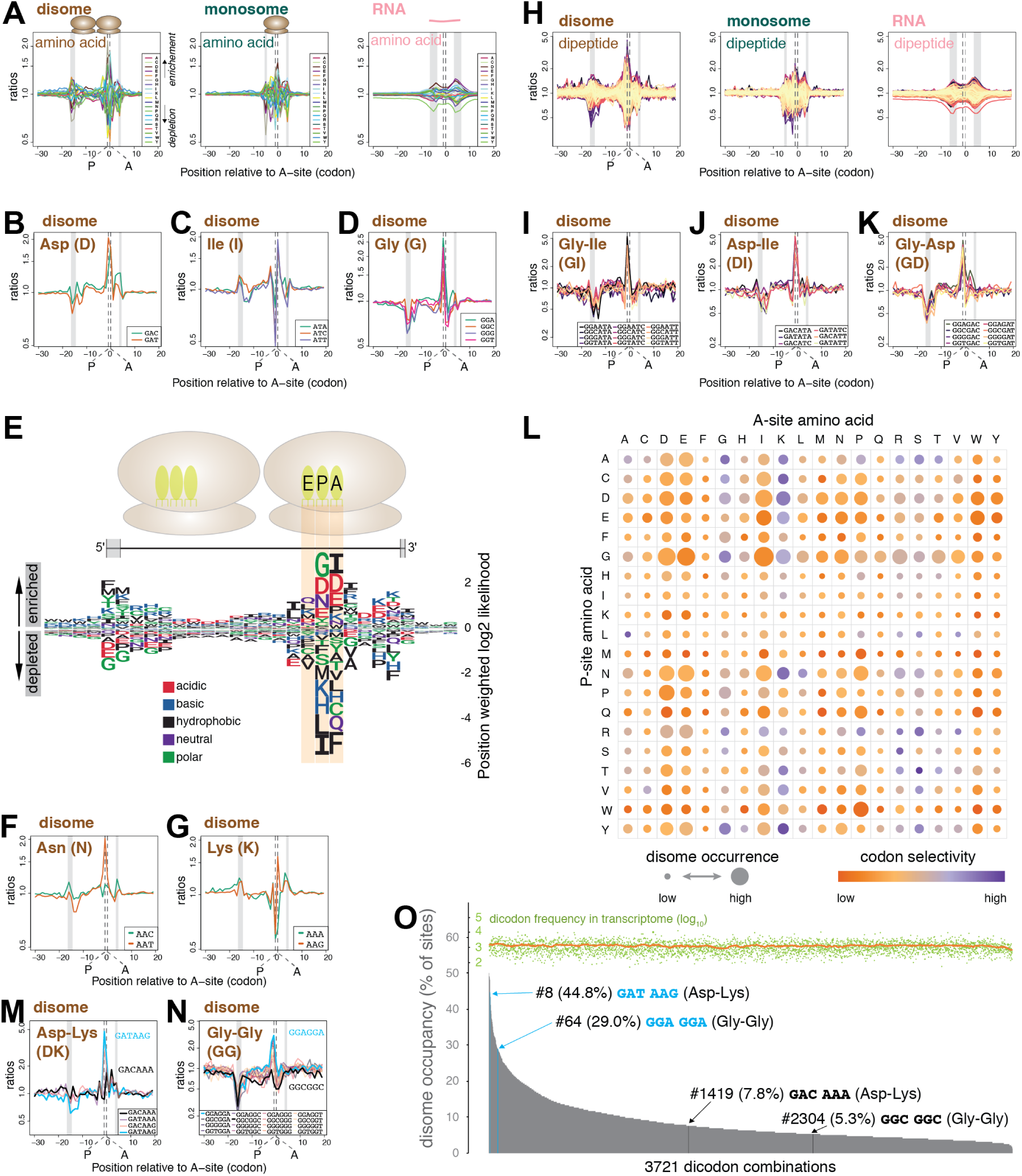
Disome sites show specific enrichment of amino acids and codons. **(A)** Position-specific enrichment analysis of proximal sequences reveals selectivity for amino acids in the decoding center of pausing ribosomes. Normalized ratios of observed-to-expected occurrences (y-axis, log-scaled) of nucleotide triplets, grouped by the amino acid they code (color codes in the right inset), are plotted for each codon position relative to the estimated A-site (0 at x-axis) of the leading ribosome of disomes (left), or of the individual ribosome in the case of monosomes (middle). For total RNA (right), position 0 denotes the midpoint of the reads. Ratios above and below 1 suggest enrichment and depletion, respectively. The vertical gray bars indicate the likely positions of the 5′ and 3′ ends of the read inserts for different library types. In addition to the A-site, positions corresponding to P-sites are also marked by vertical dashed lines. **(B - D)** Position-specific enrichment plots of sequences coding for representative amino acids at and around pause sites identified by disomes. Similar to (A), however, the triplets were not combined by the amino acids, instead shown individually (color codes in the right inset) for aspartic acid (Asp), isoleucine (Ile) and glycine (Gly), respectively, in **(B - D)**. **(E)** Position weight matrix of sequence triplets grouped by amino acids illustrates enrichment and depletion of specific amino acids within the decoding center of the leading ribosome of the disomes. Position-specific weighted *log*_2_-likelihood scores were calculated from the observed-to-expected ratios (A). Enrichments and depletions were represented with positive and negative scores, respectively. Height of each single-letter amino acid character is determined by its absolute score. At each codon position, the order of letters was sorted by the absolute scores of the corresponding amino acids, decreasing towards the 0-line. Amino acid letters are colored by their hydrophobicity and charges (color codes at bottom). The disome pair and their footprint are depicted graphically at the top. The gray zones at the extremities of the footprint denote the likely positions/regions of the 5′ and 3′ ends of the read inserts. **(F and G)** Similar to (B - D), for asparagin (Asn) **(F)** and lysine (Lys) **(G)**. **(H)** Position-specific enrichment plots of sequences coding for dipeptides. Similar to (A), however, instead of triplets and single amino acids, 6mers coding for a pair of amino acids (dipeptides) were used to calculate the observed-to-expected ratios for all possible dipeptides. Color code is not given due to vast number of dipeptides. **(I - K)** Similar to (B - D), however, enrichment of individual 6mers are shown for representative dipeptides: Gly-Ile **(I)**, Asp-Ile **(J)** and Gly-Asp **(K)**. **(L)** Enrichment and codon selectivity of all amino acid combinations corresponding to the predicted P- and A-sites of the leading ribosome. Identities of the amino acids at the P- and A-site are resolved vertically and horizontally, respectively. The area of disks represents enrichment of disome sites at the dipeptide signature. The color of disks represents the codon selectivity for a dipeptide signature, calculated as the difference between the max. and min. enrichment ratios (log) of all 6mers coding for that dipeptide. **(M and N)** Similar to (I - K), for Asp-Lys **(M)** and Gly-Gly **(N)**. **(O)** Relative disome occupancy by dicodon. Disome occupancy for the 3721 dicodon combinations was plotted in descending order. Occupancies were calculated for a given 6mer (dicodon) as the raw percentage of sites with disome to all present sites (with + without disome) across the studied transcriptome. The number of all present sites is shown at the top of the graph coloured in lime (moving average trendline in orange). Annotated are two pairs of 6mers from panels (N) and (O), coding for Asp-Lys or Gly-Gly, which show large differences in disome occupancies depending on codon usage (blue vs black fonts for high vs low occupancy, respectively).

In summary, the preference for codons, amino acids and amino acid combinations at the predicted P- and A-sites of the leading ribosome suggested that specific sequence signatures are an important contributor to the locations of collision events. Moreover, ribosome pausing that depends on codon usage opens the possibility to modulate the kinetics of translation elongation independently of amino acid coding potential.

### Disome sites are related to structural features of the nascent polypeptide

It was unlikely that the two factors associated with ribosomal collisions revealed above – i.e. high ribosomal flux (Fig. 2) and specific amino acids/codons (Fig. 3) – would suffice to provide the specificity required to discriminate between the disome sites that were actually observable (e.g. the alleged “deterministic sites” in Fig. 2I, K, M, O), as compared to other positions on the mRNA that were devoid of disome footprints despite similar codon composition. We therefore expected that additional features of the transcript and of the polypeptide would be critical in specifying ribosomal collision sites as well. Our above findings indicated that the signal peptide represented one such element promoting stalling and stacking. In this context, it was interesting that even for SP-related stalling, the actual sites on which disomes were observable were in accordance with the generic disome site characteristics described above. Thus, disome density on SP-transcripts was also dependent on TE (Fig. 2P) and the amino acid preference of disome sites at SP sequences (Supplemental Fig. S11B) closely resembled that identified transcriptome-wide (Fig. 3E).

In order to identify other protein traits associated with ribosomal stalling, we first assessed the relationship between disome sites and the electrostatic charge of the nascent polypeptide. These analyses revealed, first, a strong association of negatively charged amino acids with the decoding center of the stalled ribosome (Fig. 4A, left panel). This was an expected outcome given the enrichment of Asp and Glu in the P- and A-sites (Fig. 3E). Second, there was a broad stretch of positive charge on the nascent polypeptide that extended >20 codons upstream of the sequence actually occupied by the stalled and stacked ribosomes (Fig. 4A, left panel; red shaded area). As this region was far upstream of the footprint, the enrichment of codons specifying positively charged amino acids was not an effect of sequence bias in library generation. Importantly, the marked charge association was specific to disome footprints; in monosome footprint and RNA data it was only weakly detectable and absent, respectively (Fig. 4A, middle and right panels). These observations suggested that there was an interplay between the nascent polypeptide and the speed at which codons that were located substantially further downstream were translated (Fig. 4B). This idea is consistent with previous work that has suggested that electrostatic interactions between a positively charged nascent peptide and the negatively charged lining of the exit tunnel are a major factor promoting local slowdown of elongating ribosomes (Charneski and Hurst, 2013).

**Figure 4.**
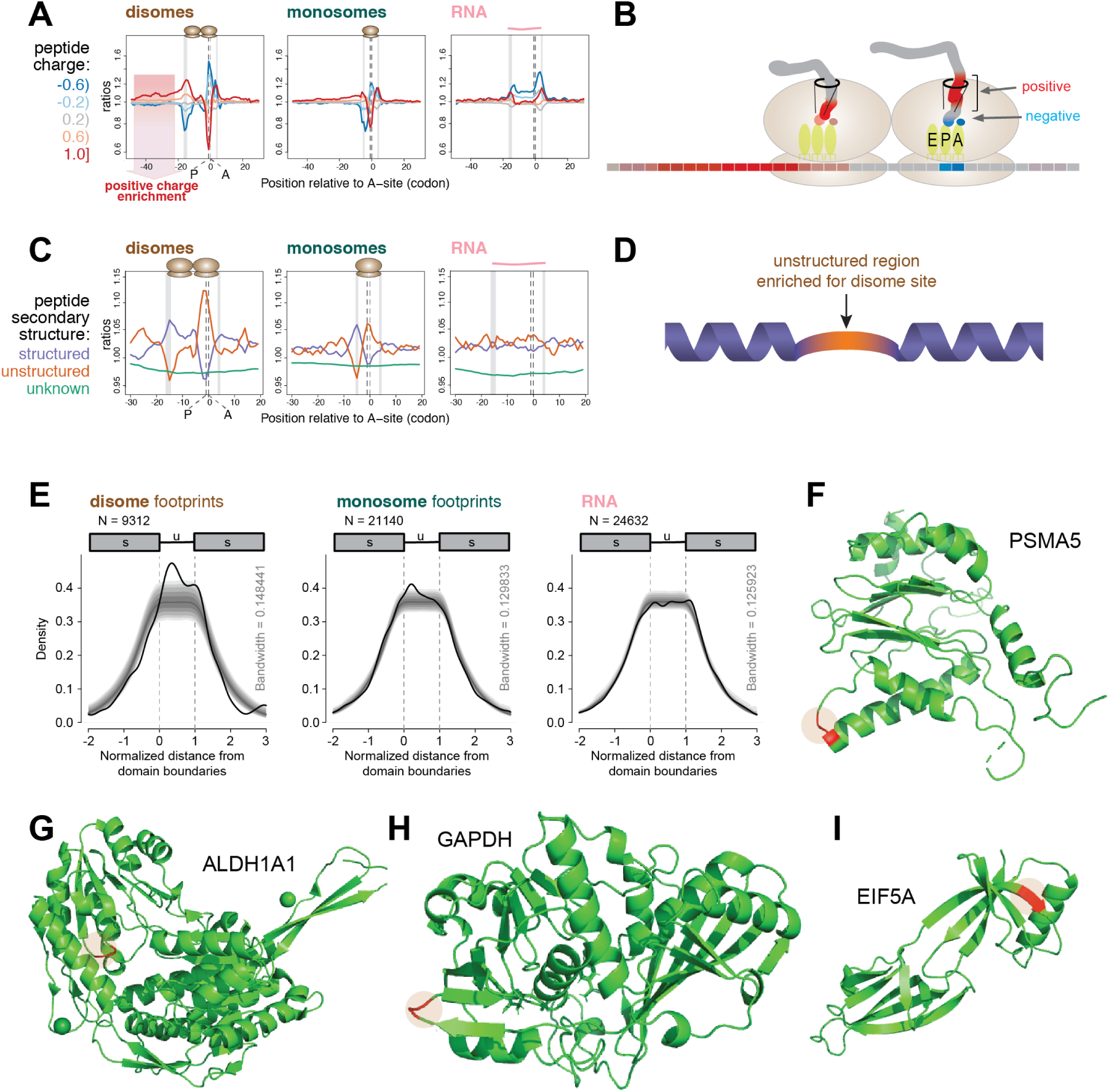
Disome site positions are related to the charge and secondary structure of the nascent polypeptide. **(A)** Position-specific enrichment analysis of proximal sequences reveals association of positive charges in the nascent polypeptide with pausing ribosomes. Average charge of 3 consecutive amino acids were stratified into 5 charge groups (interval boundaries and color codes on the left). Normalized ratios of observed-to-expected occurrences (y-axis, log-scaled) of charge groups were plotted at the center position of the tripeptide relative to the estimated A-site (0 at x-axis) of the leading ribosome of disomes (left panel), or of the individual ribosome in the case of monosomes (middle). For total RNA (right), position 0 denotes the midpoint of the reads. The vertical gray bars indicate the likely positions of the 5′ and 3′ ends of the read inserts for different library types. Positions corresponding to P- and A-sites are also marked by vertical dashed lines. **(B)** Graphical representation of the electrostatic interactions between the leading ribosome and the nascent peptide chain. Associations of negatively charged residues (blue) with the P-A sites and a stretch of positively charged residues (red) with the exit tunnel are depicted. Corresponding codons on the mRNA (series of filled rectangles at the bottom) are colored similarly based on the charge of the amino acids they code for. **(C)** Association between disome sites and the structure of the translated polypeptide. Based on the UniProt database, each position of translated peptides was labeled according to their structural information: ’structured’ for α-helix or β-sheet, ’unstructured’ and ’unknown’; β-turns were excluded. Normalized ratios of observed-to-expected occurrences (y-axis, log-scaled) of structural categories were plotted relative to the estimated A-site (0 at x-axis) of the leading ribosome of disomes (left), or of the individual ribosome in the case of monosomes (middle). See (A) for other elements. **(D)** Graphical model depicting a preference for pausing during the translation of unstructured polypeptide stretches (orange) that were preceded and followed by structured regions (purple). **(E)** Enrichment of disome sites within the unstructured stretches of polypeptides that are preceded and followed by structured regions. Structured (min. 3 aa, up to 30th position) - unstructured (min. 6, max. 30 aa) - structured (min. 3 aa, up to 30th position) regions were identified transcriptome-wide. Positions across regions were scaled to the length of the unstructured region and centered to its start, such that the start and the end of the unstructured region would correspond to 0 and 1, respectively (x-axis). Kernel density estimates (thick black lines) were calculated for peaks across normalized positions weighted with their normalized counts, estimated at the A-site of the leading ribosome for disomes (left, N = 9312), A-site of the monosomes (center, N = 21140) or center of total RNA reads (right, N = 24632). Confidence intervals for the kernel densities, which were calculated by randomly shuffling (N = 10000) peaks within each transcript, are shown by gray shaded regions (and allow estimating statistical significance of the signal): darkest at the center, 50% (median) to outward, 25%, 12.5%, 5%, 2.5% and 1%. **(F - I)** Three-dimensional structure of four individual proteins with disome site amino acids highlighted. PSMA5, structure for the *H. sapiens* homologue (PDB ID: 5VFT); ALDH1A1, structure for the *H. sapiens* homologue (4WJ9); GAPDH, structure for the *H. sapiens* homologue (4WNC), corresponding residues at position 65-66; murine EIF5A (5DLQ). The strongest disome site of each transcript is highlighted in red.

We next explored the relationship between disome sites and the structure of the translated polypeptide. Using genome-wide peptide secondary structure predictions with the three categories, structured (α-helix, β-sheet), unstructured and unknown, we calculated position-specific observed-to-expected ratios. These analyses revealed that the decoding center of the downstream ribosome was enriched for codons predicted to lie in unstructured parts of the polypeptide, whereas structured amino acids were depleted (Fig. 4C, left panel). Upstream and downstream of the stalled ribosome this pattern was inverted, with an increase in structured and a decrease in unstructured amino acids. The identical analysis on monosome footprints yielded associations that were only small, although qualitatively similar to those seen for disomes, (Fig. 4C, middle panel). Moreover, these associations were absent from the RNA data (Fig. 4C, right panel). These findings indicated a high degree of specificity for disome sites and, taken together, they were consistent with a model according to which there was a preference for pausing during the translation of unstructured polypeptide stretches that were preceded and followed by structured regions (Fig. 4D). We investigated this hypothesis more explicitly by retrieving the transcript regions encoding “structured-unstructured-structured” (s-u-s) polypeptide configurations transcriptome-wide (N=9312). After re-scaling to allow for the global alignment of structured and unstructured areas, we assessed the relative disome distributions across the s-u-s-encoding regions. These analyses revealed that disomes were enriched within the 5′ portion of the unstructured region, just downstream of the s-u boundary (Fig. 4E, left panel). By comparing with distributions obtained from randomizations of the disome peak positions within the same dataset, we could conclude that the observed disome enrichment was significantly higher than expected by chance. As before, a weak and no effect, respectively, were detectable in monosome footprint and RNA data (Fig. 4E, middle and right panels). Finally, the position-specific analysis (i.e. without re-scaling) at the s-u boundary indicated that disome sites were particularly enriched at the second codon downstream of the s-u transition (Supplemental Fig. S12A, right panel). As an additional control for the specificity of these associations, we also analyzed the inverse configuration, u-s-u (Supplemental Fig. S12D) and conducted all analyses on monosome footprint (Supplemental Fig. S12B, E) and RNA data (Supplemental Fig. S12C, F) as well. Taken together, the analyses established that the most prominent enrichment was that of disome sites within the unstructured area of the s-u-s configuration, frequently directly after the s-u boundary. Visual inspection of individual examples of where disome-associated amino acids mapped within known protein structures confirmed this notion, as shown for PSMA5, ALDH1A1, GAPDH, and EIF5A (Fig. 4F-I).

In summary, we concluded that there was a direct link between ribosomal pause sites and structural features of the nascent polypeptide. Translational pausing was thus more likely to occur while decoding negatively charged amino acids that were downstream of extended positively charged regions of the polypeptide, and within unstructured areas downstream of structured regions. These associations are suggestive of a link between elongation pausing and protein folding.

### Disome sites are enriched within distinct transcript groups and are associated with previously documented translational pauses

Thus far, only a limited number of translational pauses have been documented in the literature. One of them is the pausing associated with SRP recruitment (Wolin and Walter, 1988) that we can also observe in the disome data (Fig. 2A-G, Supplemental Fig. S3C). We wished to examine whether our analysis could provide insights into other previously described pausing events, and whether specific groups of transcripts, processes, pathways or co-translatinoal events were especially prone to pausing. To do so, we first selected the most prominent deterministic sites – i.e. pausing events that were not merely attributable to high ribosomal traffic – in order to enrich for potential functionally relevant cases (Supplemental Table S3; top 5650 disome peaks from 1185 genes). Of note, these strong disome sites showed high correspondence across all six independent biological samples (Supplemental Fig. S13), indicating that they represented reproducible strong pausing events. First, we searched whether structural data was available for the proteins with prominent disome sites. Out of the first ∼50 genes in the list, around 20-25 structures (murine proteins or orthologous proteins from other mammals) were available from published data. Mapping the disome site amino acids onto the structures revealed that in most cases, these were located in unstructured regions, and very often directly at the structured-unstructured boundary (Supplemental Fig. S14; Fig. 4F-I). These findings were consistent with the idea that efficient pausing in coding sequences may be important for the structural integrity and folding of the nascent polypeptide.

In order to determine whether specific pathways were affected by the pausing phenomenon, we next assessed if functional categories were over-represented among the genes with prominent disome peaks (top-200 genes from Supplemental Table S3). The analysis revealed a strong bias towards transcripts encoding proteins that were annotated as “co-factor and co-enzyme binding”, that were involved in “oxidation-reduction processes”, and that localized to mitochondria (Fig. 5A; Supplemental Table S4). It is tempting to speculate that the integration and/or covalent attachment of co-factors (a common feature of oxidoreductases) onto polypeptides is coordinated co-translationally and dependent on translational pausing. Moreover, there is evidence for co-translational protein localization/import to mitochondria (reviewed in Lesnik et al. (2015)). The enrichment of prominent disome peaks among transcripts encoding mitochondrially located proteins may reflect a specific feature of their translational kinetics.

**Figure 5.**
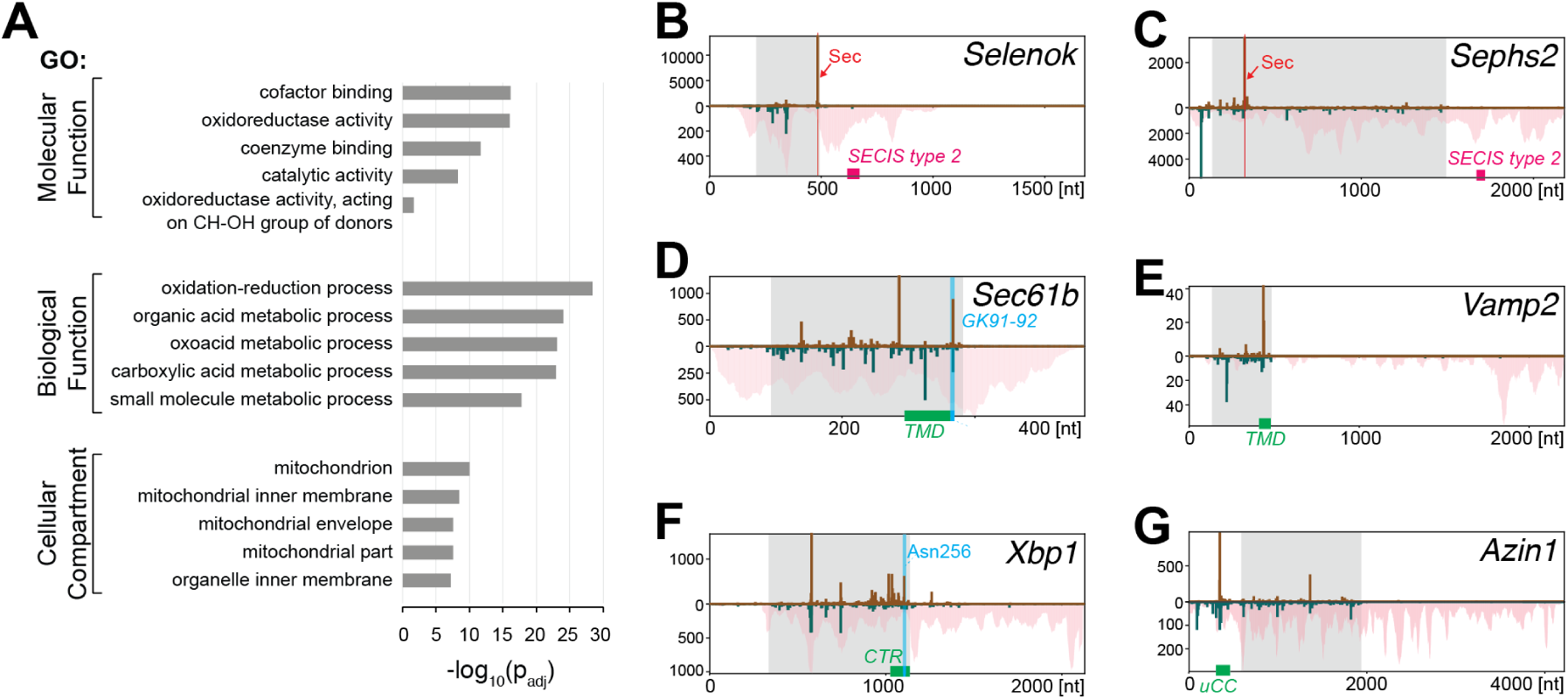
Disome site association with specific pathways and with known pausing events. **(A)** Functional enrichment analysis of top 200 genes containing most prominent deterministic sites. Five terms with the highest -log_10_(*p*_*adj*_) values (horizontal bars) are shown from each Gene Ontology (GO) group: GO molecular function, GO cellular component, GO biological process. Full analysis is given in Supplemental Table S4. **(B - G)** Distribution of normalized counts of monosome and disome footprints (per nt), similar to Fig. 2F-O, along transcripts of selected genes that carry strong disome sites. *Selenok* **(B)**, ENSMUST00000112268, and *Sephs2* **(C)**, ENSMUST0000082428, show a strong disome peak on the selenocysteine codon (Sec, marked in pink). Position of the SECIS elements is indicated in pink. *Sec61b* **(D)**, ENSMUST00000065678, and *Vamp2* **(E)**, ENSMUST00000021273, are tail-anchored proteins with a transmembrane domain (TMD, green). For *Sec61b*, a strong disome site is located on GK91-92 (marked in blue). *Xbp1* **(F)**, ENSMUST00000063084, contains a C-terminal region (CTR, green) that shows high density of disome sites over several positions. The strong site on Asn256 is marked in blue, *Azin1* **(G)**, ENSMUSG00000037458, contains an upstream conserved coding region (uCC, green) that undergoes translational elongation in a polyamine-dependent fashion, which leads to ribosome queuing and main CDS start site selection. The strong disome signal in the uCC coding region is located on a GP dipeptide at uCC amino acids 14-15.

Next, we visually inspected the prominent disome peak list (Supplemental Table S3) for notable specific cases of translational pausing. We found it striking to spot 9 transcripts specifying proteins that contained the rare amino acid selenocysteine (Sec/U) among the translatome-wide most prominent disome peaks (Supplemental Table S3; Supplemental Fig. S15A). An additional six Seccodon-containing transcripts (Supplemental Fig. S15B) were expressed in liver as well, as judged from RNA-seq and monosome footprint data, yet they showed varying degrees of disome occurrence and did not feature among the most prominent peaks translatome-wide. Sec is encoded by the UGA stop codon, whose reinterpretation involves the 3′ UTR-located Selenocysteine Incorporation Sequence element (SECIS) (Vindry et al., 2018). Decoding of selenocysteine is known to be particularly slow (Stoytcheva et al., 2006; Howard et al., 2013). It would hence appear quite plausible that ribosomal collisions occur during prolonged dwelling of elongating ribosomes on Sec codons. In several cases, for example on *Selenok (Selenoprotein K)* (Fig. 5B) and *Sephs2 (Selenophosphate synthetase 2)* (Fig. 5C), the disome peak indeed coincided with the Sec codon. In other cases, however, disome peak and Sec codon did not correspond to each other (Supplemental Fig. S15A, B). We inspected whether the presence or absence of specific RNA elements could be responsible for the differences in disome location. For example, SECIS elements come in two variants, type 1 and type 2 (Vindry et al., 2018). A diversity of other structural RNA features have been identified in conjunction with Sec decoding as well (Mariotti et al., 2017), such as the selenocysteine codon redefinition element (SRE) that consists of a stem-loop directly downstream of the UGA codon and was first described on *Selenoprotein N (Selenon)* (Howard et al., 2005, 2007). However, we did not observe any striking association between disome sites and specific RNA elements (Supplemental Fig. S15A, B and data not shown).

Further inspection of the prominent disome transcripts revealed that *Sec61b* featured within the list. *Sec61b* encodes a tail-anchored protein whose translation has been reported to slow down downstream of the single, C-terminally located transmembrane domain (TMD) (Mariappan et al., 2010). This pausing is understood to provide time to recruit the machinery for membrane insertion before the TMD is released from the ribosomal exit tunnel. Our liver data revealed several disome peaks on *Sec61b* (Fig. 5D), of which one of the strongest occurred on a GK dipeptide that was downstream of (and immediately adjacent to) the TMD and had the disome-prone codon usage GGC AAG. Mariappan et al. had observed pausing for tail-anchored proteins also on the *Vamp2* transcript. Of note, *Vamp2* carried a strong disome peak towards the 3′ end of the CDS that lay, however, within rather than after the TMD (Fig. 5E). Finally, we examined the disome profiles of other documented translational pauses. Pausing on the *Xbp1* transcript, which involves translation of a hydrophobic C-terminal region (CTR), facilitates the cotranslational mRNA localization to the endoplasmic reticulum membrane (Yanagitani et al., 2011). Our data confirm multiple disome sites in this area (Fig. 5F). A strong site was specifically on Asn256, which is the last codon required for translational arrest (Yanagitani et al., 2011) and on which a pausing event was identified by Ingolia et al. (2011). Two unusual cases of regulatory translational stalling were recently described for transcripts encoding components of the polyamine biosynthesis pathway, i.e. *Amd1* (Yordanova et al., 2018) and *Azin1* (Ivanov et al., 2018). For *Amd1*, the mechanism involves stochastic stop codon read-through and the formation of a ribosome queue at the next in-frame stop codon that eventually extends up into the main coding sequence and halts its translation. Unfortunately, due to low sequencing coverage of *Amd1*, we were unable to validate this model from the disome data. For *Antizyme inhibitor 1 (Azin1)*, Ivanov et al. recently demonstrated how a specialised uORF, termed the upstream conserved coding region (uCC) undergoes translational elongation in a polyamine-dependent fashion, which leads to ribosome queuing and main CDS start site selection (Ivanov et al., 2018). Notably, we observed strong disome signal precisely on the uCC coding region (Fig. 5G). Ivanov et al. mapped a pausing event to the PPW tricodon (uCC amino acids 47-49), whereas our data revealed strongest disome accumulation on a site corresponding to ribosome pausing further upstream, on a GP dipeptide at amino acids 14-15.

### Evolutionary conservation at disome sites suggests an active, functional role of pausing

The above examples showed that disome sites were associated with known cases of ribosomal pausing that have been functionally characterised in previous studies. Globally, however, our analyses did not allow distinguishing whether the observed ribosomal pauses were functionally important – for example to ensure independent folding of individual protein domains, undisturbed from downstream nascent polypeptide stretches – or whether they rather represented an epiphenomenon of such processes. For example, the biosynthesis of proteins and their folding could slow down translation and thus, as a downstream effect, lead to ribosome pausing and collisions, without being of functional relevance for the preceding folding event itself. We therefore sought a way to evaluate the two scenarios and reasoned that in the case of an active, functional role, the codons and dipeptide motifs on which pausing occurred would show higher evolutionary conservation than expected.

RUST analysis using PhyloP conservation scores revealed an enrichment for highly conserved codons at the P- and A-sites of the stalled ribosome (Fig. 6A). Moreover, we observed that highly conserved transcripts generally showed high disome levels while poorly conserved transcripts were rather disome-poor, especially in the group of mRNAs with high translation efficiencies (Fig. 6B). Although these analyses indicated connections between translational pausing and evolutionary conservation, they did not allow us to determine how direct this link was. Most of all, the further interpretation was rendered difficult due to the intrinsic selectivity of disome sites for specific amino acids, codons and dipeptides (Fig. 3), which can be expected to represent a strong confounding factor for simple evolutionary analyses as the one shown in Fig. 6A.

**Figure 6.**
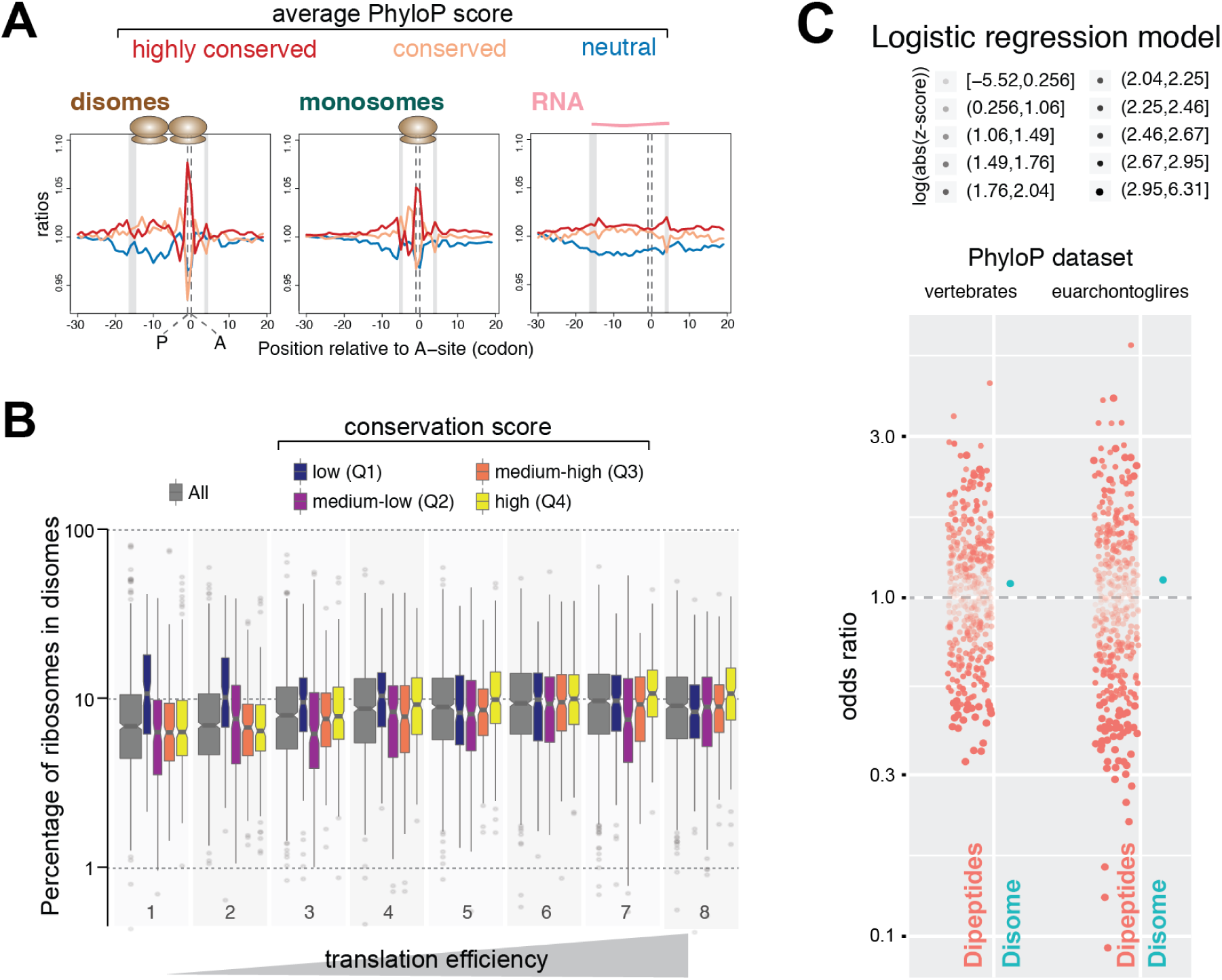
Evolutionary conservation at disome sites. **(A)** Association of evolutionary highly conserved codons with the P- and A-sites of disome sites revealed by position-specific enrichment analysis of proximal sequences. Along coding regions, PhyloP conservation scores were grouped into three categories: neutral - blue, [-3, 3), conserved - orange, [3, 5) and highly conserved [5,). Normalized ratios of observed-to-expected occurrences (y-axis, log-scaled) of conservation categories were plotted relative to the estimated A-site (0 at x-axis) of the leading ribosome of disomes (left), or of the individual ribosome in the case of monosomes (middle). For total RNA (right), position 0 denotes the midpoint of the reads. See Fig. 4A for other elements. **(B)** Box-and-whiskers illustrating the estimated percentages of ribosomes that were pertained in disomes for groups of transcripts with different overall evolutionary conservation. Groups included all detectably expressed genes (All, gray, N = 7375), which were stratified into 4 groups (N = 2270 or 2271 for each) (color code at the top) based on the quartiles of average PhyloP scores with following right-closed boundaries: −0.585, 2.327, 3.356, 4.239, 6.437. x-axis is same as in Fig. 2P. Width of each box is proportional to the number of data points it represents. **(C)** Odds ratio estimates of dipeptides and disome sites for increased PhyloP scores. Odds ratios (OR) for having a high PhyloP score at P-A dicodons were estimated for dipeptides encoded by the dicodon (orange dots, 399 levels relative to dipeptide VH, which had moderate PhyloP scores in both models) and presence of a disome peak (green dots, A-position disome density > mean transcript density) using a logistic regression model. Confidence levels of estimates were represented by transparency levels that corresponded deciles of the logarithm of absolute values of their z-scores as given by the legend. Two separate regression models were fitted using PhyloP scores from the 60-way vertebrate dataset (left) and the Euarchontoglire subset (right). For disomes, OR is larger than 1 (dashed line) indicating that it is more likely to observe a high PhyloP score when disome peaks are present than when they are absent.

As we wished to uncouple the various effects on conservation, we used logistic regression analysis to determine what contribution specifically the disome sites made to overall conservation, against the background of general dipeptide and transcript conservation scores. This analysis showed that different dipeptides, just on there own, had very distinct conservation scores (Fig. 6C). Therefore, the strong association of disome sites with high PhyloP scores seen above (Fig. 6A) was mostly attributable to the specific dipeptide bias observed at these positions. Importantly, the regression analysis further revealed that the presence of disomes increased the odds of having a highly conserved PhyloP score (cutoff: PhyloP >5 for 60-way vertebrate and >1.3 for euarchontoglires datasets, respectively) by approximately 10% (60-way vertebrate: b=0.1258, p-value < 1e-16, OR=1.134 95% CI=1.125, 1.143; euarchontoglires: b=0.1012, p-value < 1e-16, OR=1.107, 95% CI=1.098, 1.115). This outcome established that translational pauses identified by disome sites were significantly more conserved than expected by chance. While the thousands of pause sites that can be detected translatome-wide will span all categories (i.e. from deleterious to beneficial), the globally detectable signal of positive selection on the P-A codons at which ribosomes are more likely to stall, strongly argues for an active, functional role of ribosomal pausing.

### Codon usage at disome sites affects protein output from a reporter gene

The evolutionary conservation at disome sites that we detect at a global scale is in line with the idea that many translational pausing events may play active roles in polypeptide biosynthesis. Future experimental studies – ideally by creating pausing loss- and gain-of-function mutants through genetic knock-in of alternative codon usages at endogenous loci – will allow validating the biological functions of individual pausing events. In the framework of this study, we wished to gain first, preliminary insights into how pause site modification would affect protein output from a reporter gene. In order to select a suitable candidate, we speculated that in cases where polypeptides assembled into large multiprotein complexes, poorly coordinated translation kinetics would affect protein abundances due to changes in the efficiency of their incorporation into the complex. Excess unincorporated protein may be subject to degradation, mislocalization, aggregation or similar, which all may impact steady-state protein levels. In the list of strong disome sites (Supplemental Table S3) we noted that several transcripts encoding ribosomal proteins showed strong, discreet disome peaks. We selected *Rps5* and *Rpl35a* (Supplemental Fig. S16A-B) and cloned their cDNAs in-frame with firefly luciferase in a lentiviral vector that allowed internal normalization to renilla luciferase (Fig. S16C). For *Rps5*, change of codon usage at the Asp-Ile disome site from its natural GATATT (Rps5-wt), a codon usage that is particularly highly disome-prone (46.6% of sites show disomes transcriptome-wide), to the mutant version GACATC (Rps5-mut2), which is transcriptome-wide disome-poorer (26.4% of sites have disomes), led to a significant change in steady-state protein output, although both constructs encoded for precisely the same protein at the amino acid level. Other variants of *Rps5* did not show an effect, including such with disome-poorer codon usages; moreover, codon usage did not affect reporter levels for the analogous *Rpl35a* reporter (data not shown). These observations indicate that the identification of functionally important disomes sites, and testing for such function, will be one of the future challenges.

## Discussion

It has long been known that elongating ribosomes can slow down in response to obstacles that they encounter; most such pauses are assumed to be transitory and resolved in a productive manner (Schuller and Green, 2018). For certain cases, there is evidence that the translational slowdown is even an integral part of the mechanism of nascent polypeptide synthesis, as exemplified by the pausing seen on signal peptide-encoding transcripts that require targeting to the secretory pathway (Wolin and Walter, 1988). Finally, pauses can be unresolvable, thus triggering a dedicated ribosome rescue program (Joazeiro, 2019). While a number of previous studies have attempted to infer pausing from monosome footprint intensities, there is clear benefit in tracking stalled ribosomes using more direct evidence, such as specific footprint size variants (Guydosh and Green, 2014). Ribosome collisions are intrinsically linked to pausing, and the characteristics of the disome footprints that we have analyzed in this study indicate that they truly represent a steady-state snapshot of the translational pausing and collision status in mouse liver *in vivo*. A remarkable finding from our spike-based quantification of disome vs. monosome signals is the sheer quantity of translating ribosomes that are trapped in the disome state. For a typical, highly translated mRNA, we estimate that ∼10% of elongating ribosomes are affected by this phenomenon (Fig. 2P). Because certain stacking events will involve more than two collided ribosomes (see Northern blot analyses, Fig. 1A-B), our disome footprinting analyses likely even underestimate the overall ribosomal queuing phenomenon. Another potential cause of underestimation could be a loss of disomes during the purification protocol through cleavage into two individual monosomes. Two observations, however, argue against systematic loss by nuclease activity during the experiment. First, disomes show high resilience to different conditions of nuclease treatment (Supplemental Fig. S1) and, second, there is no noticable sequence bias within the disome footprint region that is located between the individual ribosomes (Supplemental Fig. S6A; Fig. 3E). Finally, when disomes are cleaved, we would expect the corresponding, 30 nt-spaced monosome footprints to appear. Previous studies have observed such 30 nt phasing of monosome footprints (presumably originating from queued ribosomes) upstream of stop codons (Andreev et al., 2015). However, the analysis of our liver datasets indicates that such monosome footprint phasing only affects a low number of very highly populated disome sites and is therefore overall rather limited in magnitude (Supplemental Fig. S17). Taken together, we view the 10% queuing rate as a realistic estimate that is, in addition, similar in magnitude to the rate of ribosome queuing of ∼20% that was recently calculated in budding yeast (Diament et al., 2018). While we consider many collisions to be an epiphenomenon of high ribosomal flux (i.e.”stochastic”, though more likely to occur on certain codons/amino acids than on others) rather than a sign of specific biological function, the loss of elongating ribosomes into queuing buildups poses challenges to the interpretation of existing ribosome profiling data. Simple monosome footprint-based quantifications very likely underestimate translation rates, especially for highly translated transcripts.

Which stalling events is our disome profiling method capturing, and which ones are missed? In yeast, stalls at truncated 3′ ends of mRNAs have recently been shown to lead to a class of small monosome footprints of ∼21 nt and, when an additional incoming ribosome stacks onto this stalled ribosome, of ∼48 nt (Guydosh and Green, 2014). Similar short footprints also occur in human cells (Wu et al., 2019), and a study in Hela cells has suggested that both transient and hard stalls trigger an endonucleolytic cleavage that leads to short footprint species as well (Ibrahim et al., 2018). It is thus likely that such short footprints reflect the more harmful pauses that provoke specific clearance pathways. The abundant ∼60 nt footprints we describe here are distinct from the above not only in size, but likely also in the translational state that they represent. They match the size reported to occur on SRP-related pausing *in vitro* (Wolin and Walter, 1988). Moreover, footprints of this size were also noted in the above yeast study (Guydosh and Green, 2014) and, although the authors did not pursue their analysis in greater detail, it is intriguing that they reported a similar bimodal size distribution and a depletion from the first codons post-initiation, just as seen in our liver data (Fig. 1E-F). We do not yet understand the significance of the two size populations; could they be associated with specific functional properties or collision states? The pattern of amino acid enrichment at the stalled ribosome is identical for the 59-60 nt and the 62-63 nt footprints (Supplemental Fig. S8A-B). Moreover, the transcriptome-wide most prominent disome-sites always consisted of a mixture of both footprint sizes, yet the relative ratio of the two size classes differed widely across pause sites (Supplemental Table S3). Despite these open questions, we can conclude from the association with signal peptides, the high steady-state abundance, and the absence of signs of mRNA cleavage at the stall site, that our disome method captures in particular the class of “benign” collisions from resolvable stalling events, including possible programmed cases.

Liver disome footprints show distinct sequence characteristics which are largely governed by the P- and A-site amino acids of the downstream ribosome (Fig. 3E). There is little specificity at the E-site, which is notable because previous work that estimated pause sites from monosome footprints identified pausing signatures with a strong E-site bias for proline (e.g. Ingolia et al. (2011); Zhang et al. (2017); Pop et al. (2014)). Due to their particular chemistry, prolines (especially in a polyproline context) are well-known for their difficult peptide bond formation and they are slow decoding, leading to translational stalls that can be resolved through the activity of eIF5A (Gutierrez et al., 2013). Structural studies have suggested direct interactions of eIF5A with a vacant E-site that has lost its empty tRNA before peptide bond formation has occurred between the growing polypepide at the P-site and the incoming charged tRNA at the A-site (Melnikov et al., 2016; Schmidt et al., 2015). Apart from a minor signal in the A-site codon (Fig. 3E; Supplemental Fig. S7), we do not see prolines associated with liver disome sites. Reassuringly, recently published disome data from mESCs (Tuck et al., 2020) does, however, show the expected polyproline motif (Supplemental Fig. S8D). This difference in disome patterns between cell types may be attributable to high activity of eIF5A in the liver. We have noted that based on monosome footprint RPKMs, eIF5A is indeed synthesized at very high levels that even exceed, for example, those of the essential elongation factor eEF2 (data not shown). Moreover, it is curious that eIF5A is itself among the 200 genes with the strongest disome peaks, occurring on a conserved Gly-Ile position (Fig. 2P, 4I; Supplemental Table S3). It would be a fascinating finding if translational pausing on *Eif5A* mRNA turned out to be part of a mechanism designed to autoregulate its own biosynthesis. Beyond the lack of proline signal in the liver data, we noted that termination codons were also absent from the disome data (see Fig. 1F). Of note, we deliberately did not pretreat our tissue samples with cycloheximide in order to avoid artefacts that this translation inhibitor is known to cause (Ingolia et al., 2011; Santos et al., 2019). However, polysome extract preparation and RNaseI digestion occurred in the presence of cycloheximide in order to avoid a loss of elongating ribosomes. We cannot exclude that terminating ribosomes may have been selectively lost at this stage, reducing the disome signal at stop codons.

The specific amino acid and dipeptide motifs that we find enriched at the paused ribosomes (Fig. 3; Supplemental Table S2) show some resemblance as well as distinct differences to previous reports. For example, Asp and Glu have been associated with presumed pause sites (Ingolia et al., 2011; Ibrahim et al., 2018), and Asp codons also figure among those whose footprint signal increases strongest (apart from Pro) in cells deficient of eIF5A (Pelechano and Alepuz, 2017; Schuller et al., 2017). The striking association of pause sites with isoleucine is an unexpected outcome of our study, as to our knowledge this amino acid is not typically reported among the top-listed associations with paused ribosomes. For GI, DI, and a subset of NI dicodons (and a number of non-isoleucine dicodons as well) transcriptome-wide 35-50% of such sites carry a strong disome footprint (Supplemental Table S2). These findings allow a bold speculation, which is that on top of the simple three-letter codon table, a six-letter code punctuates translation to organize the biosynthesis of nascent polypeptides into segments separated by intermittent pause sites. Of note, the phenomenon that translation speed can be governed at the level of the dicodon is actually well known from work on bacterial translation (Irwin et al., 1995). Our analysis from mammalian translation identified thousands such intermittent pause (Supplemental Table S3) which, as an ensemble, likely reflect an array of different protein biosynthetic phenomena whose common denominator is local ribosome slowdown. Importantly, even on the global set, an association of disome sites with structural features of the nascent polypeptide is evident (Fig. 4), as is the signature of evolutionary conservation at disome sites (Fig. 6). Conceivably, this indicates that one of the major reasons for pausing could lie in the coordination of translation with the folding, assembly or structural modification of the nascent polypeptide. Interestingly, there is compelling recent evidence from yeast that many multiprotein complexes assemble in a co-translational fashion (Shiber et al., 2018) and that the association of the individual subunits involves translational pausing (identified from high monosome footprint peaks) during the biosynthesis of the nascent polypeptides (Panasenko et al., 2019). Notably, the showcase example identified in the latter study are two proteins of the yeast proteasome regulatory particle, Rpt1 and Rpt2, whose elongation pausing leads to the association of the translating ribosomes into heavy particles (”assemblysomes”) in which the nascent peptides assemble into the multiprotein complex. Intriguingly, we find that also in mouse liver, a set of proteasomal protein mRNAs carries high disome peaks, e.g. *Psmd5* (Fig. 2I) and more than 10 other proteasome subunits (Supplemental Table S3), indicating potential conservation of the mechanism.

In conclusion, we are confident that the disome profiling methodology that we present here is an important complementary technique to the already available ribosome profiling repertoire. Although it delivers a ’snapshot’ of the translation status (not unlike conventional ribosome profiling), the cellular disome state provides specific, new information on the translation kinetics. Undoubtedly, the kinetics will show regulation in different organisms, tissues, cell types, and under different physiological conditions, which will manifest in distinct disome profiles. It will be exciting to collect and analyze such data across experimental models, allowing to evaluate differences in kinetics and the extent to which translational pausing represents an obligatory, potentially regulated event that contributes to physiological gene expression output. Through such new datasets, and already through the extensive data we have collected and analyzed in the framework of this study and in recent work in mESCs (Tuck et al., 2020), important questions are likely to become experimentally accessible, such as on the assembly of multiprotein complexes, on the co-translational attachment of protein co-factors, on the mechanics of biosynthesis of transmembrane proteins, or on the coupling of translation and RNA decay.

## Methods

### Experimental methods details

#### Experimental model and subject details

Liver extracts from 12-week-old male C57BL/6 mice were the same as reported previously (Janich et al., 2015), with experiments approved by the Veterinary Office of the Canon Vaud (authorization VD2376). Liver extracts for spike-in experiments were prepared independently for this study; all details on extract preparation can be found in the Supplemental Material. NIH3T3 and HEK293FT cell lines were the same as described Janich et al. (2015) and cultured under standard conditions (see Supplemental Material).

#### Northern blot

Northern blots were in principle conducted as described in Gatfield et al. (2009). Briefly, RNA purified from tissue extracts that were treated with nuclease were separated by polyacrylamide gel electrophorese, electroblotted on membrane, immobilised, and hybridised with radioactively labelled oligonucleotides antisense to the *Alb* and *Mup7* transcripts. For details on the protocol and oligonucleotide sequences see Supplemental Material. Please note that the lower part of the Northern blot panels shown in Supplemental Fig. S1B was the same as in our previous publication (Janich et al., 2016).

#### Footprint and library generation (monosome, disome, RNA)

The original mouse liver datasets for monosome footprints and RNA-seq that we used in the current study were the same as those reported in Janich et al. (2015), of which we used the three timepoints, ZT0, 2, 12 (two biological replicates per timepoint; each sample is a pool of liver lysates from two mice). The matching disome footprint datasets from the same samples were produced within the framework of the current study. Disome footprints had already been cut simultaneously, and from the same gels, together with the monosome footprints in Janich et al. (2015). Library generation occurred with a modified ribosome profiling protocol, which is detailed in Supplemental Material. Briefly, libraries were generated according to Illumina’s protocol for TruSeq Ribo Profile (RPHMR12126, Illumina), using the Ribo-Zero Gold rRNA Removal Kit (MRZG12324 Illumina). Details, including for the spike-in experiment, can be found in Supplemental Material. All libraries were sequenced in-house on Illumina HiSeq 2500. Mouse ES-cell data were from Tuck et al. (2020).

#### Cloning, lentiviral production, luciferase assays

The generation of the lentiviral Rps5 dual luciferase (Firefly/Renilla) reporter plasmid, *Rps5* CDS is described in Supplemental Material. Plasmids were used to produce lentiviral particles in HEK293FT cells as described (Salmon and Trono, 2007). 1-2 weeks after lentiviral transduction, NIH3t3 cells were lysed in passive lysis buffer and luciferase activities were quantified using DualGlo luciferase assay system and a GloMax 96 Microplateluminometer (all from Promega). Firefly/Renilla luciferase (FL/RL) of the Rps5 wt plasmid were internally set to 100% in each experiment, and mutant FL/RL ratios expressed relative to wt.

### Computational methods details

#### Basic Analysis of Sequencing Reads

Preprocessing of sequencing reads, mapping and quantification of mRNA and footprint abundances largely followed protocols as described in Janich et al. (2015); a detailed description is given in Supplemental Material.

#### Spike-in Normalization and Global Quantification of Ribosomes Retained in Disomes

Design and sequences of the spike in oligos (30mers and 60mers that were concatamers of the 30mer sequences), mapping procedure and the counting algorithm to avoid counting degradation products of the 60mers as 30mers, are all described in Supplemental Material. In the sequencing data, spike reads were mapped and processed similarly to all other reads. Spike counts were first normalized for library size with upper-quantile method and spike-in normalization factors were calculated as 60mer / 30mer ratios per sample to correct the experimental biases between the disome and monosome counts. The spike-normalized counts of disomes and monosomes were then used to estimate the percentage of ribosomes that were identified within disomes to the whole, taking into account that each disome represented two ribosomes.

#### Observed-to-Expected Ratios For Proximal Sequence Features

The calculation of observed-to-expected ratios for sequence features proximal to footprint sites was performed following the principles of Ribo-seq Unit Step Transformation method (O′Connor et al., 2016). The described method was extended by including additional features such as 6mer, dipeptide, charge, secondary structure, and PhyloP conservation in addition to codon and amino acids. A margin of 30 nt was excluded from each end of the CDS. The analysis window (typically 50-codon wide) was moved along the CDS regions at single nt or 3-nt steps. Enrichment was calculated as the observed-to-present ratio normalized to the expected ratio. All analyses were performed with in-house Python (creation of data matrices) and R software (visualization and statistical analysis). Unless stated otherwise, samples were treated as replicates and in most analyses were combined. Additional details are found in Supplemental Material.

#### Estimation of A-site Positions

The A-sites of the monosomes (RPF) were calculated identically as described in Janich et al. (2015). For disomes, for initial analyses we used a similar approach to estimate the A-site of the upstream ribosome in the disome pair as 15 nt from the 5’ end of the footprints. This approach was suitable for exploratory analyses (e.g meta-transcript analysis) for facilitating the comparability to monosome results. In other analyses, we used an empirical method to estimate the A-site of the leading (downstream) ribosome within the disome pair. In order to infer the optimum offsets for different lengths of footprints, we first split the disome footprints by their size, from 55 to 64 nt. Wihtin each size group, footprints were further split into 3 classes based on their open reading frame relative to that of the main CDS. For each group, position-specific (relative to their 5’ ends at nucleotide resolution) information content matrices were calculated using the Kullback-Leibler divergence scores (O′Connor et al., 2016) of observed-to-expected ratios of codon analysis (see Calculation of Expected-to-observed Ratios For Proximal Sequence Features). For combinations of footprint size and open reading frame, where the position of PA sites could be identified as highest information positions (with 2 peaks 3 nt apart from each other) around 40 - 50 nt downstream of the 5’ ends of the footprints, exact offsets were calculated as the distance of the deduced A-site from the 5’ end. Following offsets for 58, 59, 60, 62 and 63 nt long disome footprints on different open reading frames were used, respectively: [45, 44, 43], [45, 44, 46], [45, 44, 46], [48, 47, 46], [48, 47, 49]. Total RNA reads were offset with different methods to be consistent with the dataset they were being compared to: by their center (general), +15 (when compared to monosomes, also selecting a similar size range of 26-35 nt) or disome offsetting (selecting a size range of 58-63).

#### Other Computational Methods

The Supplemental Material contains a detailed description of other computations methods used in the study, including meta-transcript analyses, the analysis of footprint densities in relation to peptide secondary structures, the analysis of evolutionary conservation at disome sites, the mapping of disome amino acids onto protein three-dimensional structures, and the functional enrichment analysis of genes with prominent disome peaks

## Data Access

The raw sequencing data and processed quantification data from this study have been submitted to the NCBI Gene Expression Omnibus (GEO; http://www.ncbi.nlm.nih.gov/geo/)

## Acknowledgments

We thank the Lausanne Genomics Technologies Facility for high-throughput sequencing infrastructure, and Alex Tuck and members of the Gatfield lab for comments on the manuscript. DG acknowledges funding by the Swiss National Science Foundation through the National Centre of Competence in Research (NCCR) RNA & Disease (grant no. 141735) and individual grant 179190.

*Author Contributions* ABA: conceptualization, investigation, data curation, formal analysis, validation, visualization, methodology, writing draft; AL: investigation, formal analysis, methodology; MDM: Investigation, formal analysis; RD: formal analysis, visualization; PJ: investigation, methodology; DG: conceptualization, resources, formal analysis, supervision, funding acquisition, visualization, project administration, writing draft.

## Supplemental Files

**Supplemental Figures:** Supplemental Figures S1-S16.

**Supplemental Table S1:** Sequencing and mapping information.

**Supplemental Table S2:** Amino acid enrichment at disome site (by dicodon).

**Supplemental Table S3:** Transcripts with prominent (’deterministic’) disome peaks.

**Supplemental Table S4** Enrichment analyses for top-200 genes from Supplemental Table S3.

## References

Andreev, D.E., O’Connor, P.B., Zhdanov, A.V., Dmitriev, R.I., Shatsky, I.N., Papkovsky, D.B., Baranov, P.V., 2015. Oxygen and glucose deprivation induces widespread alterations in mrna translation within 20 minutes. Genome Biology 16, 90.

Charneski, C.A., Hurst, L.D., 2013. Positively charged residues are the major determinants of ribosomal velocity. PLOS Biology 11, 1–20.

Dana, A., Tuller, T., 2012. Determinants of translation elongation speed and ribosomal profiling biases in mouse embryonic stem cells. PLOS Computational Biology 8, 1–11.

Dao Duc, K., Song, Y.S., 2018. The impact of ribosomal interference, codon usage, and exit tunnel interactions on translation elongation rate variation. PLOS Genetics 14, 1–32.

Darnell, A.M., Subramaniam, A.R., O’Shea, E.K., 2018. Translational control through differential ribosome pausing during amino acid limitation in mammalian cells. Molecular Cell 71, 229 – 243.e11.

Diament, A., Feldman, A., Schochet, E., Kupiec, M., Arava, Y., Tuller, T., 2018. The extent of ribosome queuing in budding yeast. PLOS Computational Biology 14, 1–21.

Doring, K., Ahmed, N., Riemer, T., Suresh, H.G., Vainshtein, Y., Habich, M., Riemer, J., Mayer, M.P., OBrien, E.P., Kramer, G., Bukau, B., 2017. Profiling ssb-nascent chain interactions reveals principles of hsp70-assisted folding. Cell 170, 298 – 311.e20.

Gamble, C.E., Brule, C.E., Dean, K.M., Fields, S., Grayhack, E.J., 2016. Adjacent codons act in concert to modulate translation efficiency in yeast. Cell 166, 679 – 690.

Gatfield, D., Le Martelot, G., Vejnar, C.E., Gerlach, D., Schaad, O., Fleury-Olela, F., Ruskeepää, A.L., Oresic, M., Esau, C.C., Zdobnov, E.M., Schibler, U., 2009. Integration of microrna mir-122 in hepatic circadian gene expression. Genes & Development 23, 1313–1326.

Gobet, C., Weger, B.D., Marquis, J., Martin, E., Neelagandan, N., Gachon, F., Naef, F., 2020. Robust landscapes of ribosome dwell times and aminoacyl-trnas in response to nutrient stress in liver. Proceedings of the National Academy of Sciences doi:10.1073/pnas.1918145117.

Gutierrez, E., Shin, B.S., Woolstenhulme, C., Kim, J.R., Saini, P., Buskirk, A., Dever, T., 2013. eif5a promotes translation of polyproline motifs. Molecular Cell 51, 35 – 45.

Guydosh, N.R., Green, R., 2014. Dom34 rescues ribosomes in 3’ untranslated regions. Cell 156, 950 – 962.

Han, Y., Gao, X., Liu, B., Wan, J., Zhang, X., Qian, S., 2014. Ribosome profiling reveals sequence-independent post-initiation pausing as a signature of translation. Cell Research 24, 842–851.

Hinnebusch, A.G., 2014. The scanning mechanism of eukaryotic translation initiation. Annual Review of Biochemistry 83, 779–812.

Howard, M.T., Aggarwal, G., Anderson, C.B., Khatri, S., Flanigan, K.M., Atkins, J.F., 2005. Recoding elements located adjacent to a subset of eukaryal selenocysteine-specifying uga codons. The EMBO Journal 24, 1596–1607.

Howard, M.T., Carlson, B.A., Anderson, C.B., Hatfield, D.L., 2013. Translational redefinition of uga codons is regulated by selenium availability. Journal of Biological Chemistry 288, 19401–19413.

Howard, M.T., Moyle, M.W., Aggarwal, G., Carlson, B.A., Anderson, C.B., 2007. A recoding element that stimulates decoding of uga codons by sec trna[ser]sec. RNA 13, 912–920.

Ibrahim, F., Maragkakis, M., Alexiou, P., Mourelatos, Z., 2018. Ribothrypsis, a novel process of canonical mrna decay, mediates ribosome-phased mrna endonucleolysis. Nature Structural & Molecular Biology 25, 302–310.

Ingolia, N., Lareau, L., Weissman, J., 2011. Ribosome profiling of mouse embryonic stem cells reveals the complexity and dynamics of mammalian proteomes. Cell 147, 789–802.

Ingolia, N.T., Ghaemmaghami, S., Newman, J.R.S., Weissman, J.S., 2009. Genome-wide analysis in vivo of translation with nucleotide resolution using ribosome profiling. Science 324, 218–223.

Ingolia, N.T., Hussmann, J.A., Weissman, J.S., 2019. Ribosome profiling: Global views of translation. Cold Spring Harbor Perspectives in Biology 11, a032698.

Irwin, B., Heck, J.D., Hatfield, G.W., 1995. Codon pair utilization biases influence translational elongation step times. Journal of Biological Chemistry 270, 22801–22806.

Ivanov, I.P., Shin, B.S., Loughran, G., Tzani, I., Young-Baird, S.K.,, Cao, C., Atkins, J.F., Dever, T.E., 2018. Polyamine control of translation elongation regulates start site selection on antizyme inhibitor mrna via ribosome queuing. Molecular Cell 70, 254 – 264.e6.

Janich, P., Arpat, A., Castelo-Szekely, V., Lopes, M., Gatfield, D., 2015. Ribosome profiling reveals the rhythmic liver translatome and circadian clock regulation by upstream open reading frames. Genome Research 25, 1848–1859.

Janich, P., Arpat, A.B., Castelo-Szekely, V., Gatfield, D., 2016. Analyzing the temporal regulation of translation efficiency in mouse liver. Genomics Data 8, 41 – 44.

Joazeiro, C.A.P., 2019. Mechanisms and functions of ribosome-associated protein quality control. Nature Reviews Molecular Cell Biology 20, 368–383.

Johnstone, T., Bazzini, A., Giraldez, A., 2016. Upstream orfs are prevalent translational repressors in vertebrates. EMBO J 35, 706–723.

Lesnik, C., Golani-Armon, A., Arava, Y., 2015. Localized translation near the mitochondrial outer membrane: An update. RNA Biology 12, 801–809.

Mariappan, M., Li, X., Stefanovic, S., Sharma, A., Mateja, A., Keenan, R.J., Hegde, R.S., 2010. A ribosome-associating factor chaperones tail-anchored membrane proteins. Nature 466, 1120–1124.

Mariotti, M., Shetty, S., Baird, L., Wu, S., Loughran, G., Copeland, P.R., Atkins, J.F., Howard, M.T., 2017. Multiple RNA structures affect translation initiation and UGA redefinition efficiency during synthesis of selenoprotein P. Nucleic Acids Research 45, 13004–13015.

Melnikov, S., Mailliot, J., Shin, B.S., Rigger, L., Yusupova, G., Micura, R., Dever, T.E., Yusupov, M., 2016. Crystal structure of hypusine-containing translation factor eif5a bound to a rotated eukaryotic ribosome. Journal of Molecular Biology 428, 3570 – 3576.

O′Connor, P.B.F., Andreev, D.E., Baranov, P.V., 2016. Comparative survey of the relative impact of mrna features on local ribosome profiling read density. Nature Communications 7, 12915.

Panasenko, O.O., Somasekharan, S.P., Villanyi, Z., Zagatti, M., Bezrukov, F., Rashpa, R., Cornut, J., Iqbal, J., Longis, M., Carl, S.H., Penña, C., Panse, V.G., Collart, M.A., 2019. Co-translational assembly of proteasome subunits in not1-containing assemblysomes. Nature Structural & Molecular Biology 26, 110–120.

Pelechano, V., Alepuz, P., 2017. eIF5A facilitates translation termination globally and promotes the elongation of many non polyproline-specific tripeptide sequences. Nucleic Acids Research 45, 7326–7338.

Pop, C., Rouskin, S., Ingolia, N.T., Han, L., Phizicky, E.M., Weissman, J.S., Koller, D., 2014. Causal signals between codon bias, mrna structure, and the efficiency of translation and elongation. Molecular Systems Biology 10, 770.

Salmon, P., Trono, D., 2007. Production and titration of lentiviral vectors. Curr Protoc Hum Genet Chapter 12, Unit 12.10.

Santos, D.A., Shi, L., Tu, B.P., Weissman, J.S., 2019. Cycloheximide can distort measurements of mRNA levels and translation efficiency. Nucleic Acids Research 47, 4974–4985.

Schmidt, C., Becker, T., Heuer, A., Braunger, K., Shanmuganathan, V., Pech, M., Berninghausen, O., Wilson, D.N., Beckmann, R., 2015. Structure of the hypusinylated eukaryotic translation factor eIF-5A bound to the ribosome. Nucleic Acids Research 44, 1944–1951.

Schuller, A.P., Green, R., 2018. Roadblocks and resolutions in eukaryotic translation. Nature 19, 526–541.

Schuller, A.P., Wu, C.C.C., Dever, T.E., Buskirk, A.R., Green, R., 2017. eif5a functions globally in translation elongation and termination. Molecular Cell 66, 194 – 205.e5.

Shiber, A., Doring, K., Friedrich, U., Klann, K., Merker, D., Zedan, M., Tippmann, F., Kramer, G., Bukau, B., 2018. Cotranslational assembly of protein complexes in eukaryotes revealed by ribosome profiling. Nature 561, 268–272.

Sinturel, F., Gerber, A., Mauvoisin, D., Wang, J., Gatfield, D., Stubblefield, J.J., Green, C.B., Gachon, F., Schibler, U., 2017. Diurnal oscillations in liver mass and cell size accompany ribosome assembly cycles. Cell 169, 651–663.

Stoytcheva, Z., Tujebajeva, R.M., Harney, J.W., Berry, M.J., 2006. Efficient incorporation of multiple selenocysteines involves an inefficient decoding step serving as a potential translational check-point and ribosome bottleneck. Molecular and Cellular Biology 26, 9177–9184.

Tuck, A.C., Rankova, A., Arpat, A.B., Liechti, L.A., Hess, D., Iesmantavicius, V., Castelo-Szekely, V., Gatfield, D., Bühler, M., 2020. Mammalian rna decay pathways are highly specialized and widely linked to translation. Molecular Cell.

Vindry, C., Ohlmann, T., Chavatte, L., 2018. Translation regulation of mammalian selenoproteins. Biochimica et Biophysica Acta 1862, 2480 – 2492.

Wolin, S., Walter, P., 1988. Ribosome pausing and stacking during translation of a eukaryotic mrna. EMBO J 7, 3559–3569.

Wu, C.C.C., Zinshteyn, B., Wehner, K.A., Green, R., 2019. High-resolution ribosome profiling defines discrete ribosome elongation states and translational regulation during cellular stress. Molecular Cell 73, 959 – 970.e5.

Yanagitani, K., Kimata, Y., Kadokura, H., Kohno, K., 2011. Translational pausing ensures membrane targeting and cytoplasmic splicing of xbp1u mrna. Science 331, 586–589.

Yordanova, M.M., Loughran, G., Zhdanov, Alexander V. and Mariotti, M., Kiniry, S.J., OConnor, P.B.F., Andreev, D.E., Tzani, I., Saffert, P., Michel, A.M., Gladyshev, V.N., Papkovsky, D.B., Atkins, J.F., Baranov, P.V., 2018. Amd1 mrna employs ribosome stalling as a mechanism for molecular memory formation. Nature 553, 356–360.

Zerial, M., Melancon, P., Schneider, C., Garoff, H., 1986. The transmembrane segment of the human transferrin receptor functions as a signal peptide. EMBO J 5, 1543–1550.

Zhang, S., Hu, H., Zhou, J., He, X., Jiang, T., Zeng, J., 2017. Analysis of ribosome stalling and translation elongation dynamics by deep learning. Cell Systems 5, 212 – 220.e6.

