## Supplemental Figures and Methods for "Transcriptome-wide sites of collided ribosomes reveal principles of translational pausing"

#### SUPPLEMENTAL MATERIAL

### Supplemental Figures and Legends

**Supplemental Figure S1. Higher-order ribosome protected fragments are highly reproducible under various assay conditions.**

**(A)** Northern blot similar to that shown in Figure 1A (probe $\text{Mup7}_{58-81}$), but from experiments in which extract preparation and RNase I digestion was preformed at different temperatures and with harsher detergent conditions, as indicated.

**(B)** Similar to (A), but under conditions in which the concentration of RNase I was varied, as indicated. Probes $\text{Mup7}_{58-81}$ (left panel) and $\text{Alb}_{71-101}$ (right panel) were used to detect possible changes in disome footprints across conditions.

**(C)** Similar to (A) and (B), but using a different nuclease, micrococcal nuclease (MNase). Probes $\text{Alb}_{71-101}$ (left panel), $\text{Mup7}_{58-81}$ (middle panel) and $\text{Mup7}_{298-320}$ (right panel) were used to detect variations in disome footprints across conditions. Note that although MNase produced somewhat different patterns than RNase I, disome footprints were still readily detectable.

### Supplemental Figure S1

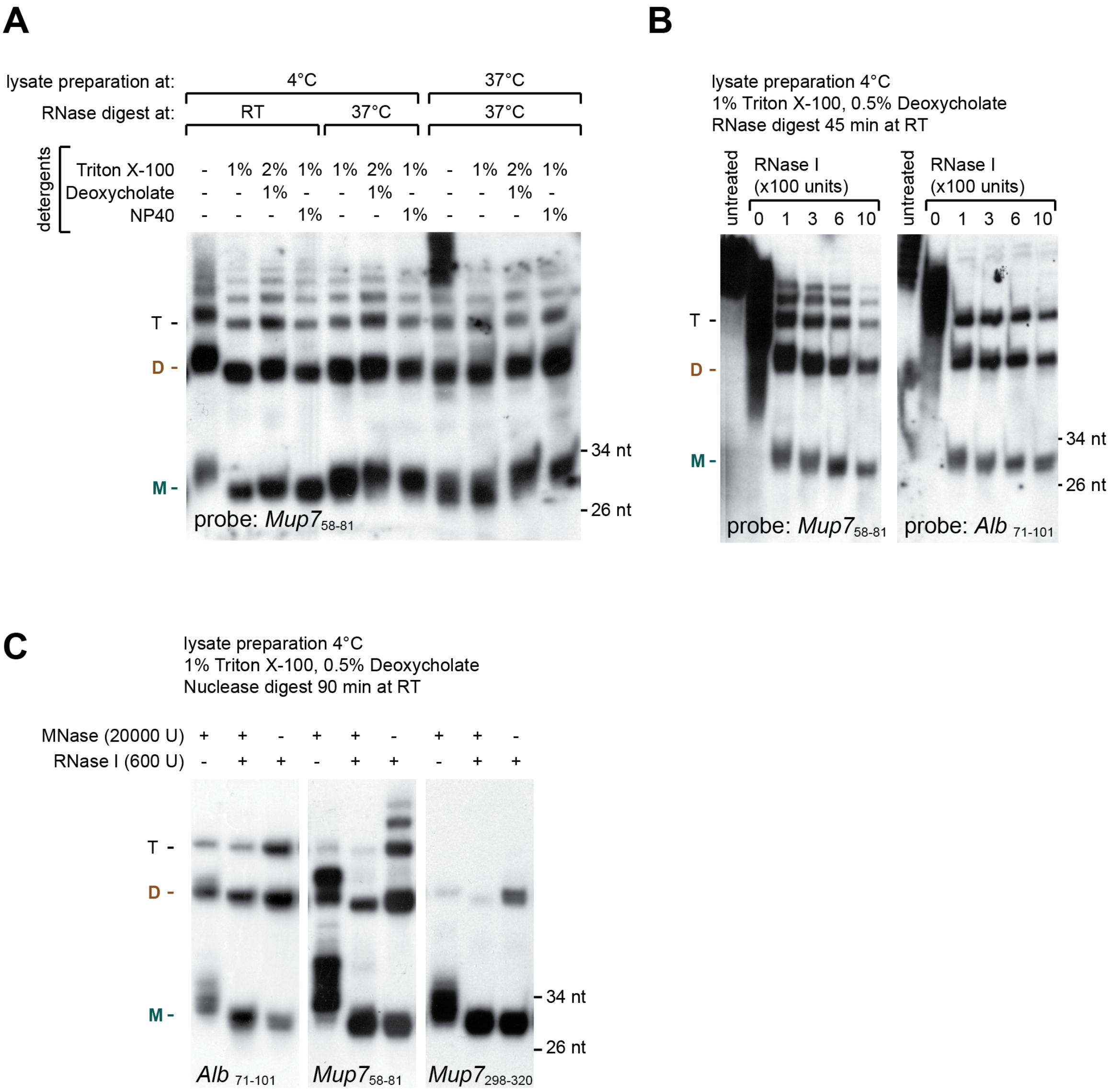

**Supplemental Figure S2. Mapping characteristics of disome reads.**

**(A)** Pie-charts of percentages of reads from monosome, disome and total RNA libraries that were mapped to different sequence types (rRNA, human rRNA, mt-tRNA, tRNA, mouse cDNA and mouse genome) or were unmmaped. Color codes are given in top right legend.

**(B)** Read distribution within 5′ UTRs, CDS, and 3′ UTRs for monosome (teal), disome (brick red), and total RNA (pink) data compared with the distribution expected by chance, which is determined by the feature sizes (gray; $N = 7413$). Note the enrichment of disomes reads within CDS and the depletion from UTRs, similar to that of monosome reads.

#### Supplemental Figure S2

A

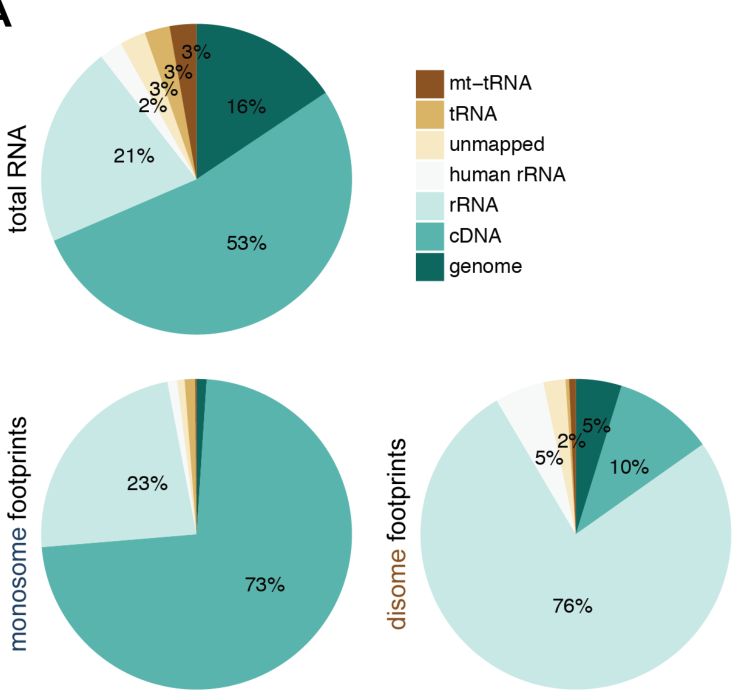


B

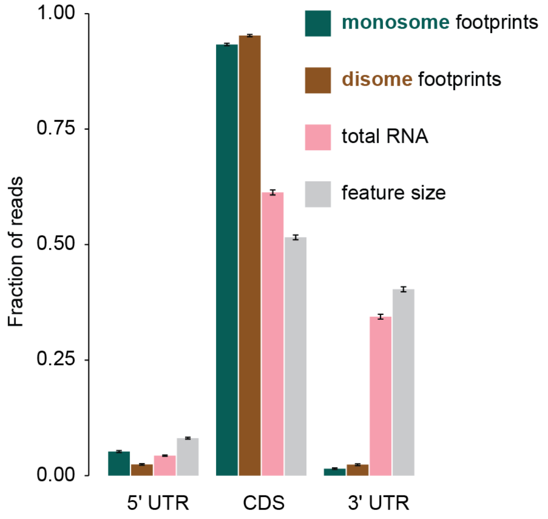

**Supplemental Figure S3. Signal peptide and translational efficiency explain some portion of the observed disome sites.**

**(A)** Density distribution of disome reads within 120 nt from the start or -120 nt from the stop codons reveals a 3-nt periodicity of disome footprints within coding sequences. Similar to Figure 1F, except for disomes, the meta-transcript analysis is aligned relative to the predicted A-site of the lagging, rather than the leading ribosome.

**(B)** Density distribution of disome reads within 120 nt from the start or -120 nt from the stop codons reveals a 3-nt periodicity of disome footprints within coding sequences. Similar to Figure 1F, yet instead of a standard alignment of the A-site of the downstream ribosome using +45 nt offset, the A-site prediction used the footprint length-dependent A-site corrections described in Supplemental Figure S6.

**(C)** Heatmap of disome footprint density across transcripts. Estimated A-sites of footprints (blue dots) are plotted relative to the start (left) or stop (right) codon of each transcript (orange horizontal lines). Transcripts were ranked (y-axis) by their CDS length. Single transcript genes that contained at least 10 disome footprints (N=9454) were included in the analysis (top). A subset of signal peptide encoding transcripts (N=1116) were analysed separately (bottom). To account for differences in expression levels, footprint densities were normalized to the footprint sum for each transcript. Disome densities (low to high) were visualized by six shades of blue (light to dark). A general trend of high-to-low disome density is observed from small to large CDS containing transcripts (top). A high-to-low disome density drop from the start to end of CDS is only observed in signal peptide transcripts (bottom).

**(D)** Time dependent changes in translation efficiencies of ribosomal proteins result in changes in occurrence of disomes. Kernel density estimates of the difference in relative disome densities (log(disome density/monosome density)) between ZT12 and ZT2 were found to be significantly different for ribosomal proteins (RP, N = 57, dashed green line) and others (N = 8558, black line) by the two-sample Kolmogorov-Smirnov test (K-S test, D = 0.1911, p = 0.032). RP genes were identified to have increased translation efficiencies at ZT12 compared to ZT2 (Janich et al., 2015).

**(E)** Difference in translational efficiencies explains only a small portion of the variation in relative disome densities among transcripts. Relative disome density (log(average normalized disome counts / average normalized monosome counts)) for all transcripts in the dataset (N = 8626) was regressed (black line) on the translational efficiency (TE, log(average normalized monosome counts / average normalized total RNA counts)). Statistics on the coefficient and the regression are given in upper part of the scatter-plot. TE explained significantly a small (2.7%) portion of the variance. Dashed line markes the b = 0 line.

#### Supplemental Figure S3

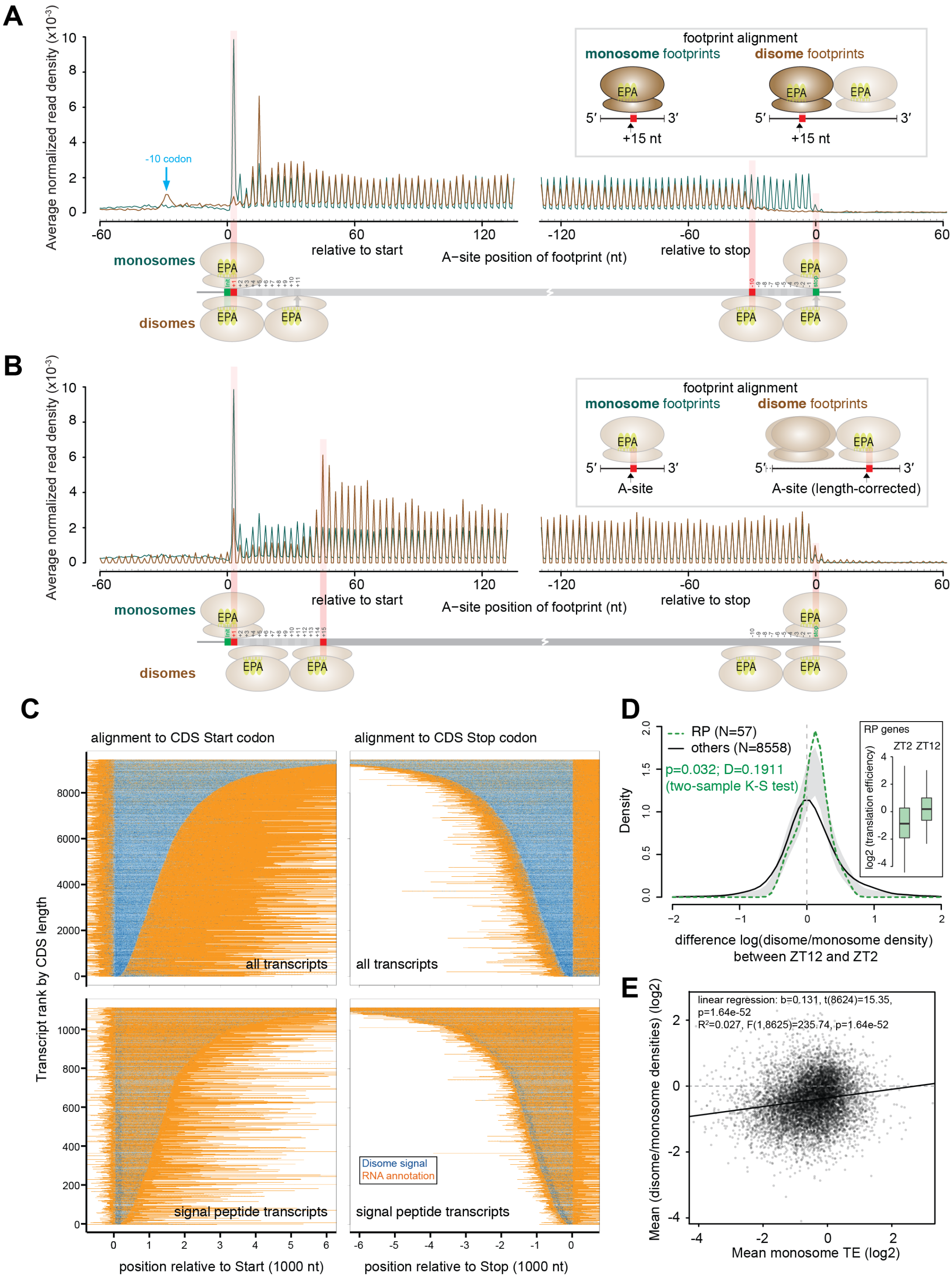

**Supplemental Figure S4. Reproducibility of disome profiles determined by individual timepoint analysis of gene graphs with stochastic and deterministic sites.**

**(A - J)** As in Figure 2F-O, these graphs show the distribution of normalized counts of monosome and disome footprints (per nt) along transcripts of representative genes. In contrast to Figure 2F-O, the data for the three timepoints for which libraries were generated, *Zeitgeber* Time (ZT) 0, 2 and 12, are plotted individually as three individual graphs. Disomes (brick red) on upward axis; monosomes (teal) and totalRNA (pink, pile-up) on downward axis. Normalised read counds on y-axis. Transcript coordinates (nt) are shown on x-axis. For simplicity, only CDS regions with short parts of flanking UTRs are plotted. SP and SA refer to signal peptide and signal anchor, respectively, and are marked by small red boxes along the x-axis. Plots are shown for *Adhesion G protein-coupled receptor G3 (Adgrg3), Transferrin receptor (Tfrc), Proteasome 26S Subunit, Non-ATPase* 4 *(Psmd4)* and *5 (Psmd5), Aldehyde dehydrogenase 1a1 (Aldh1a1), Pyruvate kinase liver and red blood cell (Pklr)* and *Eukaryotic translation initiation factor 5A (Eif5a)* in (A - J), respectively.

### Supplemental Figure S4

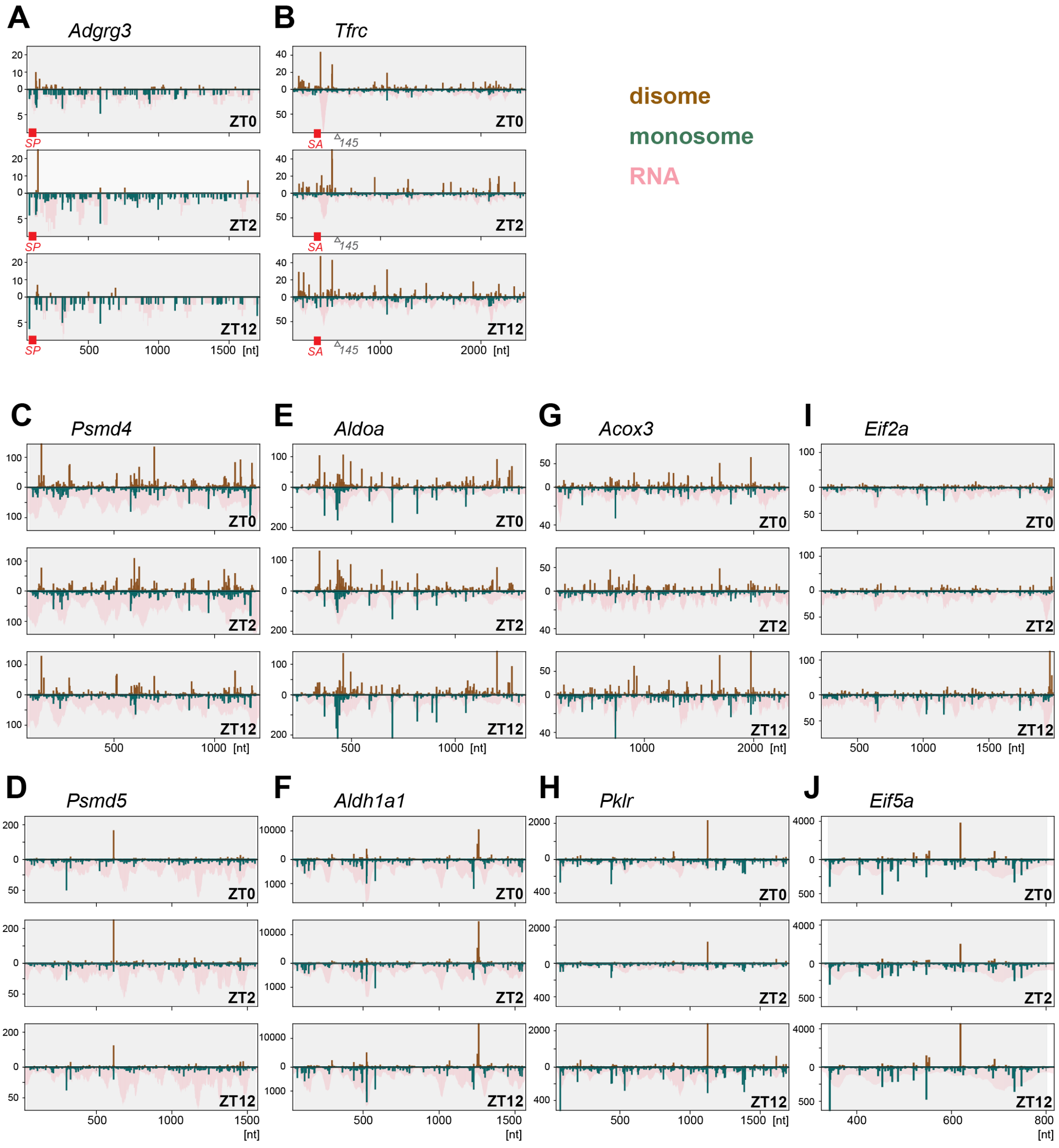

**Supplemental Figure S5. Experimental setup for spike-ins.**

**(A)** Graphical representation of experimental setup for sequencing of monosome and disome footprints spiked with pre-synthesized 30 and 60 nt RNA oligonucleotides.

**(B)** Northern blots of spike-in oligonucleotide mixes to assess the apparent ratio of the 60 nt oligos to 30 nt oligos. Same radioactively 5′ labelled DNA oligo could hybridize to both 30 nt and 60 nt RNA oligos, however, two probe molecules could hybridize to a single 60 nt oligo at the same time, therefore expected signal ratios were around 2.

### Supplemental Figure S5

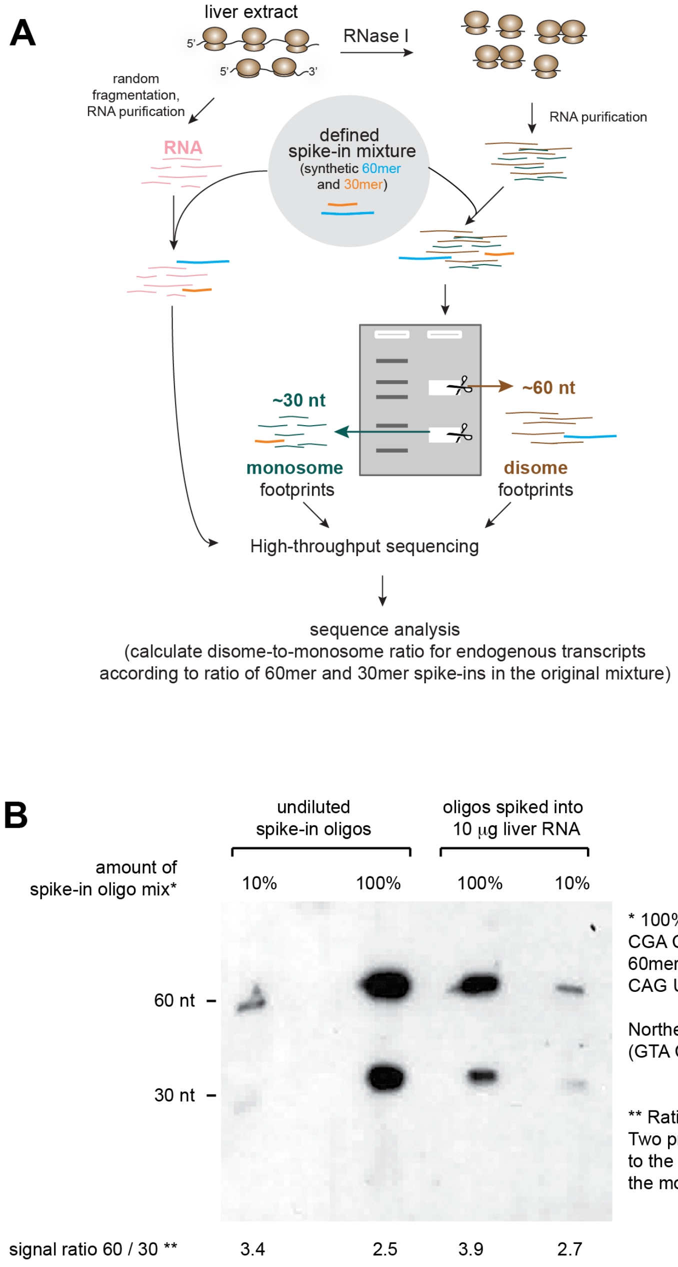


* 100% corresponds to 0.16 ng of 30mer RNA oligo (AAC UAC CGA CUC AUC CCA UCU UGC CAG UAC) and 0.32 ng of 60mer RNA oligo (AAC UAC CGA CUC AUC CCA UCU UGC CAG UAC AAC UAC CGA CUC AUC CCA UCU UGC CAG UAC)

Northern blot detection with 5' labelled DNA antisense oligo (GTA CTG GCA AGA TGG GAT GAG TCG GTA GTT)

** Ratio of signals from 60mer vs. 30mer spike-in RNA oligo. Two probe oligos can hybridise to the 60mer, whereas one oligo to the 30mer, which needs to be accounted for when calculating the molar ratio.

**Supplemental Figure S6. Empirical identification of the offsets for estimation of the A-sites of the leading ribosome of the disome pair.**

**(A)** Position-specific information content for different size classes (55 - 64 nt) of disome footprints combined with their frame (5′ position relative to the main CDS's open reading frame - 0, 1 or 2) revealed the optimal offsets for estimating the A-site of the leading ribosome. Position-specific information content was calculated using the Kullback-Leibler divergence scores of observed-to-expected ratios of codon analysis (similar to Figure 3A but without any A-site estimation - only using the 5' ends of the footprints) as described elsewhere (O′Connor et al., 2016). For each size group, the KL plots were drawn separately for three frame offsets (color code at the top of the figure). For combinations of footprint size and frame, where information content could be resolved at nucleotide level within the expected region of the decoding center (2 - 3 peaks in KL), offsets from the footprint's 5′ end were calculated to the most probable position of the A-sites (colored rectangles). Frequencies of each size group are given at the right side in million reads. For each plot, 5′ and 3′ ends of the footprints were marked with vertical dashed lines and the region occupied by the footprint was shaded in a gray tone.

**(B)** Graphical model illustrating the different configurations of disomes evidenced by the two major populations of 59-60 nt and 62-63 nt footprints. Based on (A), the difference between these two size groups is whether the ribosomes were collided completely (top), or a small gap of a single codon was left between the two (bottom).

### Supplemental Figure S6

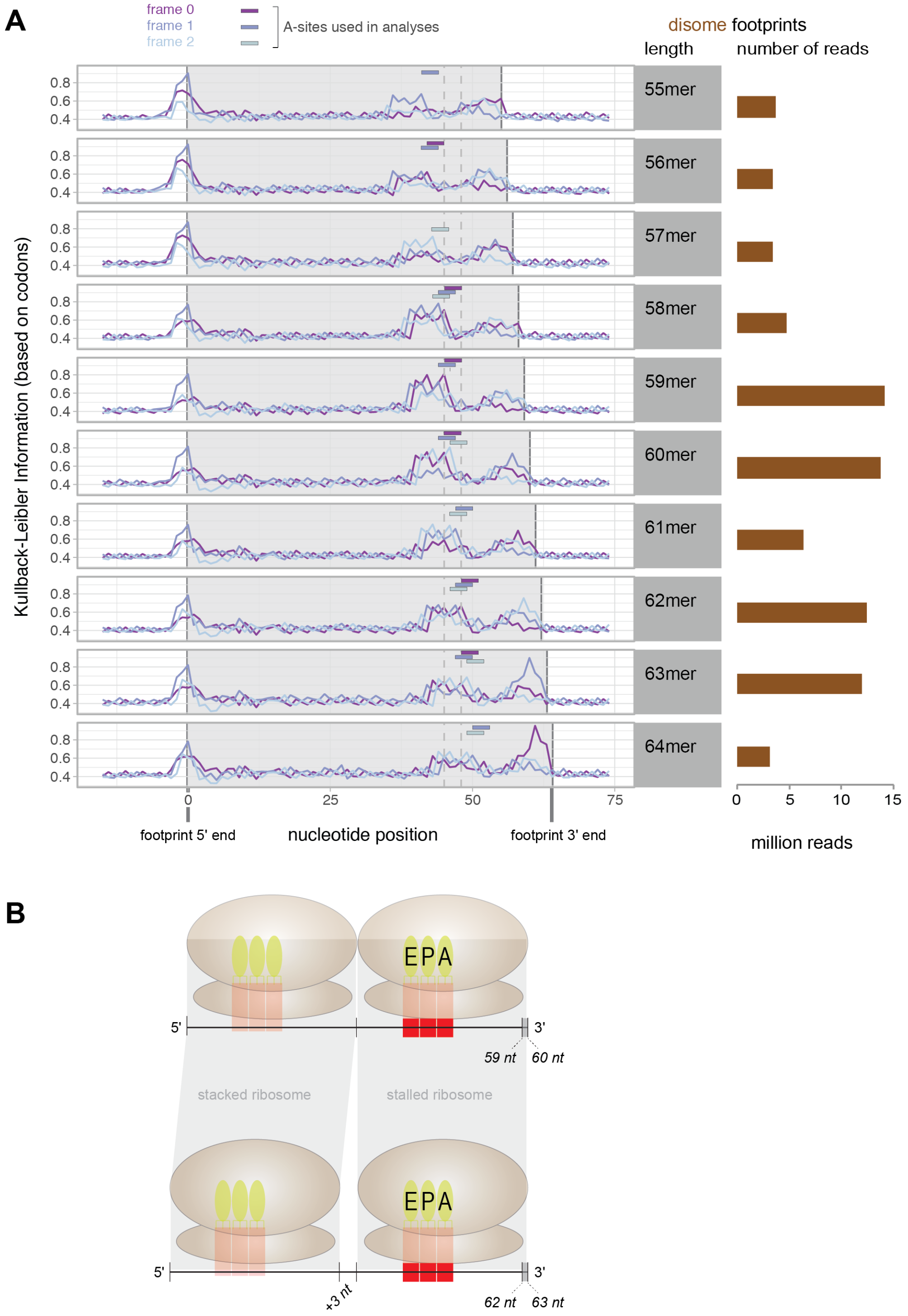

**Supplemental Figure S7. Enrichment of amino acids and codons at disome sites.**

Identical to the analysis in main Figure 3A-D, but for the complete set of amino acids, and for disome, monosome and RNA data. See figure legend for Figure 3A for details.

### Supplemental Figure S7

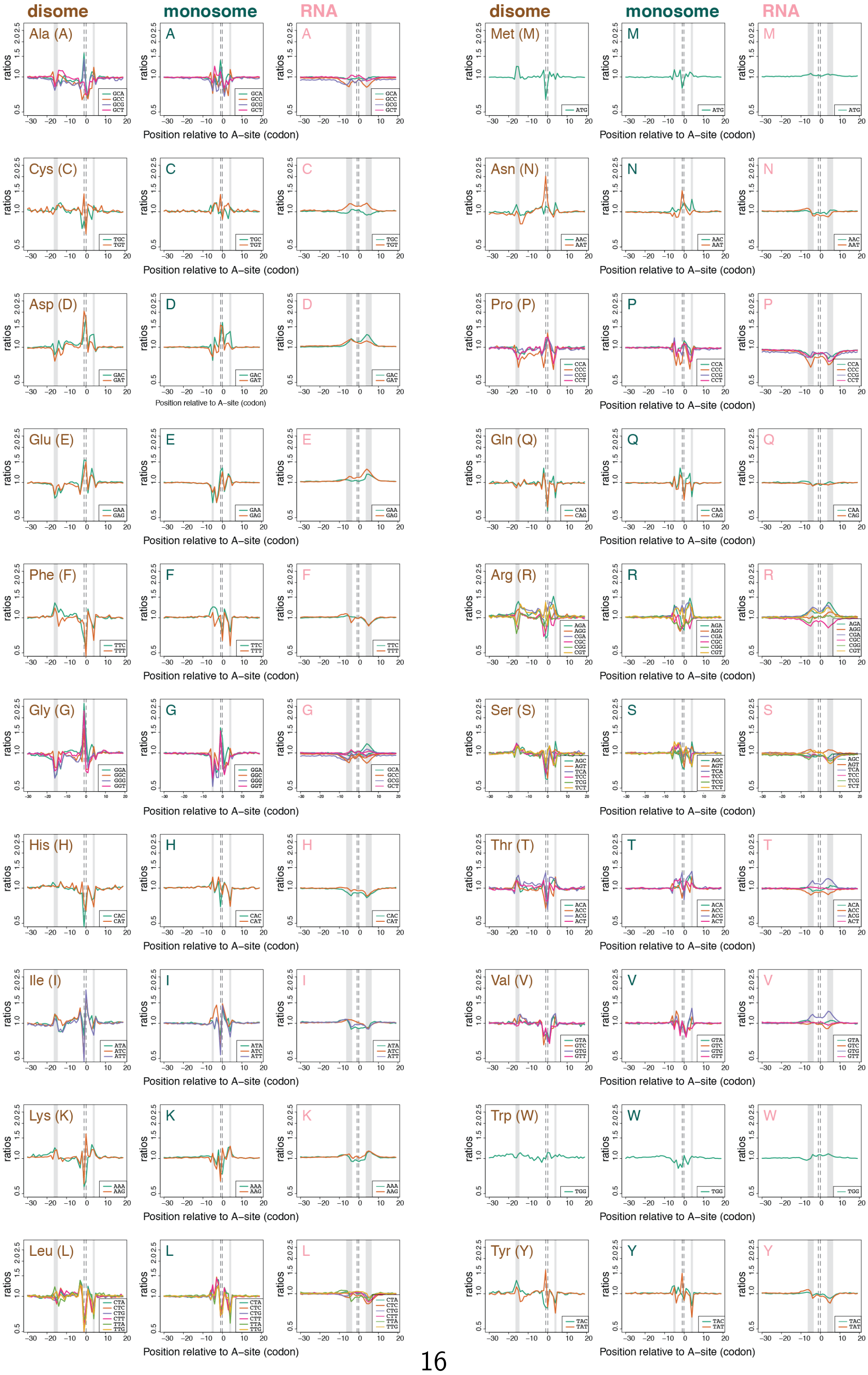

**Supplemental Figure S8. Amino acid logo at disome sites for different footprint sizes and in mESCs.**

These graphs show that similar amino acid enrichment profiles were obtained independently of footprint sizes and across different samples. Moreover, there were some notable differences between mouse liver and mouse ESCs.

**(A)** Amino acid logo plot from enrichment analyses as in Figure 3E, yet only on the 59-60 nt footprints from the liver data.

**(B)** Amino acid logo plot from enrichment analyses as in Figure 3E, yet only on the 62-63 nt footprints from the liver data.

**(C)** Amino acid logo plot from enrichment analyses as in Figure 3E, yet using the independent libraries from other mouse livers, i.e. the samples that were also used for the spike-in experiments. The footprints of sizes 59, 60, 62, 63 nt were used.

**(D)** Amino acid logo plot from enrichment analyses as in Figure 3E, yet using the disome data from mouse ES cells produced for the recent study of (Tuck et al., 2020). Please note that the logo shows an enrichment of Pro at the E- and P-sites that has also been reported for mESC monosome data by Ingolia and colleagues (Ingolia et al., 2011)

#### Supplemental Figure S8

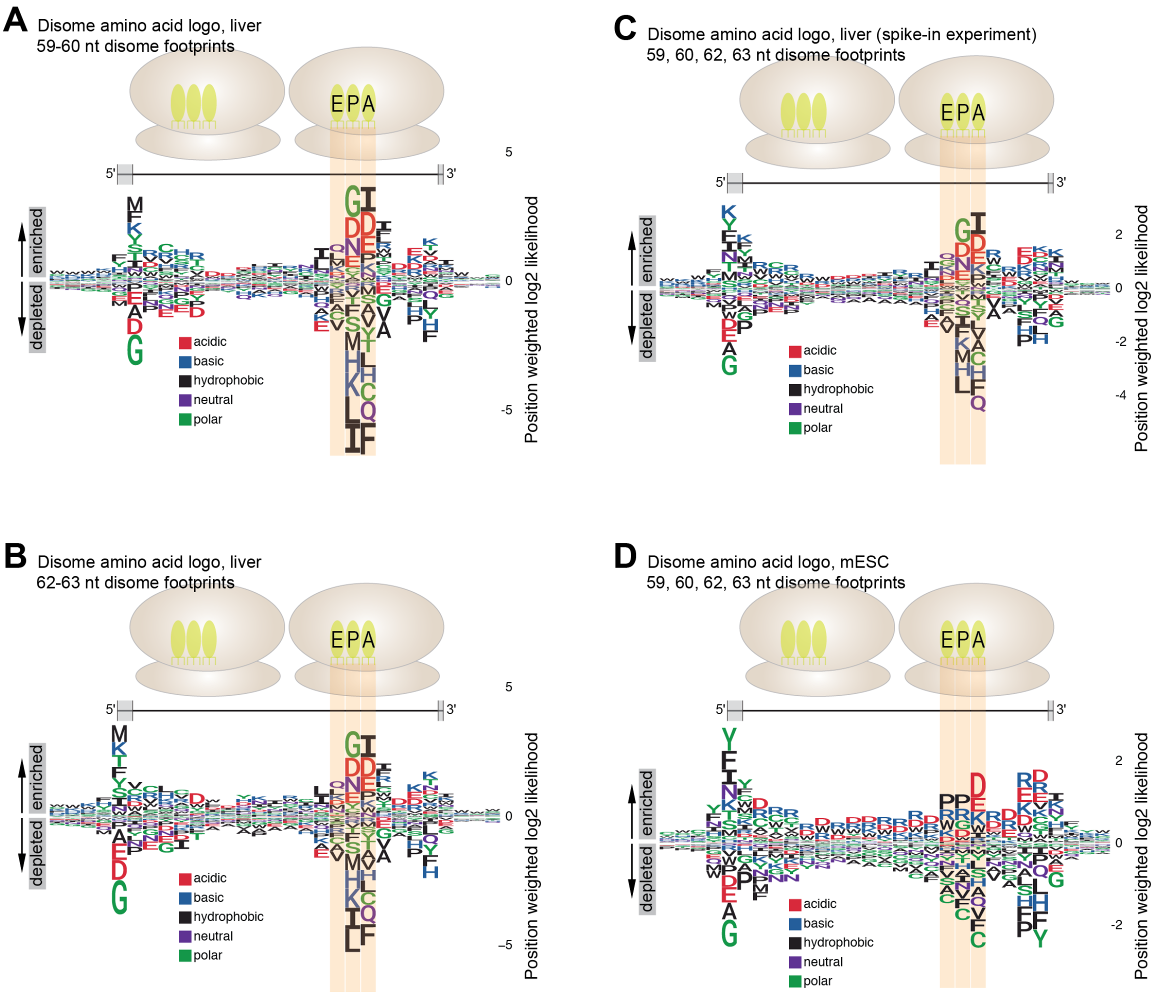

**Supplemental Figure S9. Amino acid enrichment at disome sites, calculated for individual timepoints/libraries.**

Graphs show the amino acid enrichment plots at disome sites, stratified for the 6 independent libraries from ZT0, ZT2 and ZT12. Note that there is near-identical amino acid preferences within the E-, P- and A-sites of the paused ribosome, attesting to the high biological and technical reproducibility of the disome profiling approach.

### Supplemental Figure S9

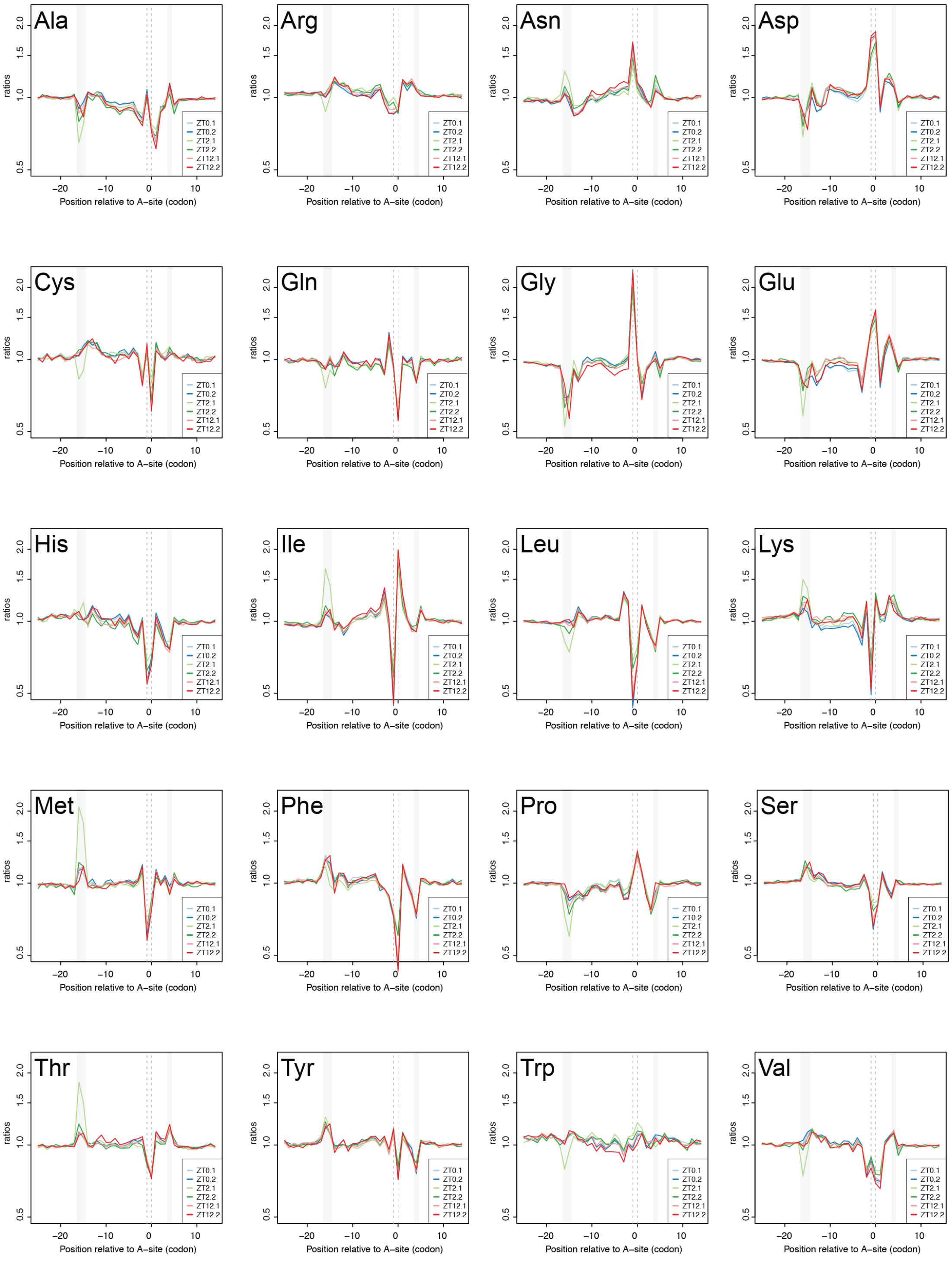

**Supplemental Figure S10. Comparison of enrichment/depletion of amino acids and codons between disome and monosome data.**

**(A)** The plots show the enrichment values (as fold-change) for amino acids at monosome sites (x-axis) vs. disome sites (y-axis) for the footprint P-site (upper panel) and A-site positions (lower panel). The 1:1 diagonal is plotted as a grey dotted line. Enrichments were calculated and plotted individually for the six libraries (i.e., every amino acid symbol is present 6 times per plot).

**(B)** As in (A), but individually for codons.

Both analyses show that, first, the observed preferences at disome sites can also be found at monosome sites, where they are however weaker (note that the relationship is steeper than the 1:1 diagonal). Second, the individual amino acids and codons behave similarly across the independent libraries (i.e. clustering of the 6 corresponding symbols, one each presenting a library), which indicates robustness the of the observed specificity across independent samples.

#### Supplemental Figure S10

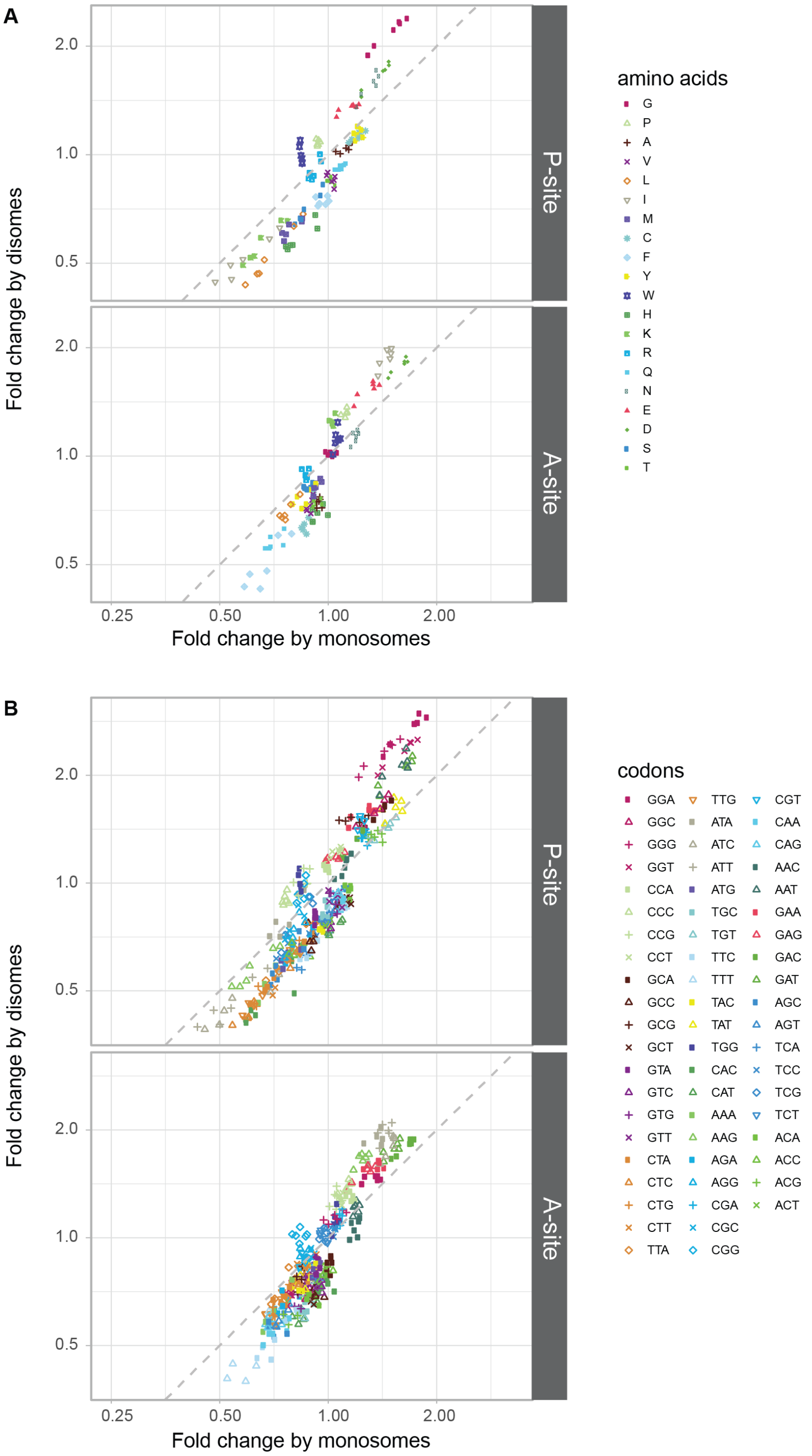

**Supplemental Figure S11. Position weight matrix for disomes, monosomes and RNA data, and specifically on the signal peptide.**

**(A)** Position weight matrix of sequence triplets grouped by amino acids illustrates enrichment and depletion specific amino acids within the decoding center of the leading ribosome of the disomes. At the top of the panel, the same data as in main Figure 3E (disomes) is shown. Middle and lower parts of the panel depict the identical analysis for monosome and RNA-seq data, respectively. See figure legends to Figure 3E for details. Of note, the figure shows that monosome footprints had a similar, though in magnitude massively reduced preference for amino acids compared to the disomes. No specificity was found in total RNA, as expected.

**(B)** As in panel (A) and in Figure 3E, but the position weight matrix was calculated only from the disomes that were found over codons 8-75 of signal peptide-containing transcripts, i.e. over the positions where the disomes related to SRP recruitment are located. Interestingly, even in this area the global pattern of amino acids at which disomes were preferentially found, corresponded to the pattern seen transcriptome-wide.

#### Supplemental Figure S11

**A**

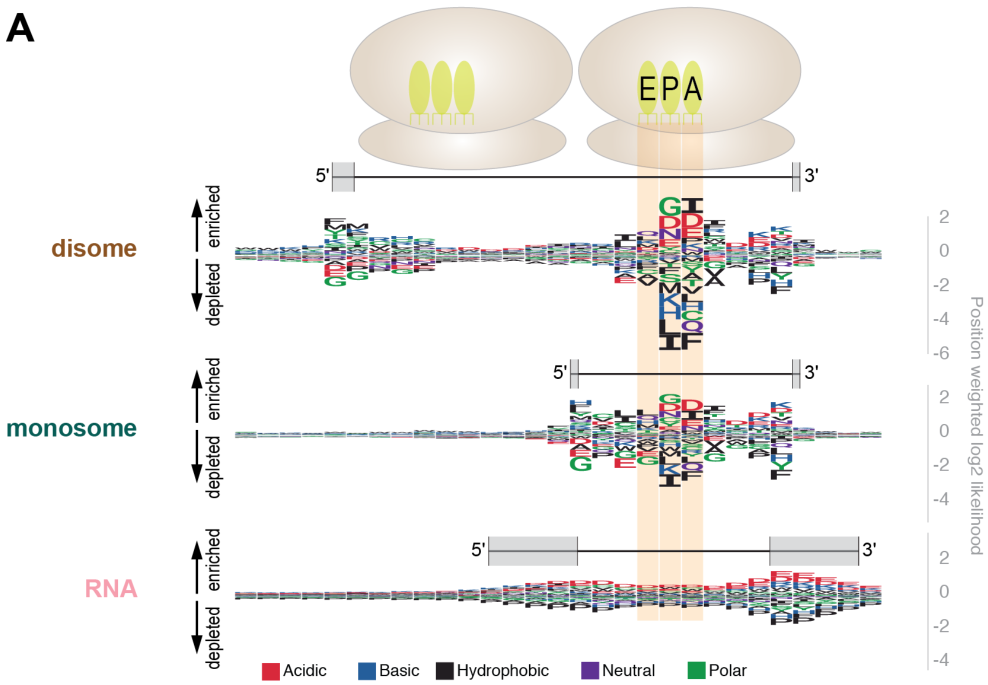


**B** SP-transcripts (analysis over codons 8-75)

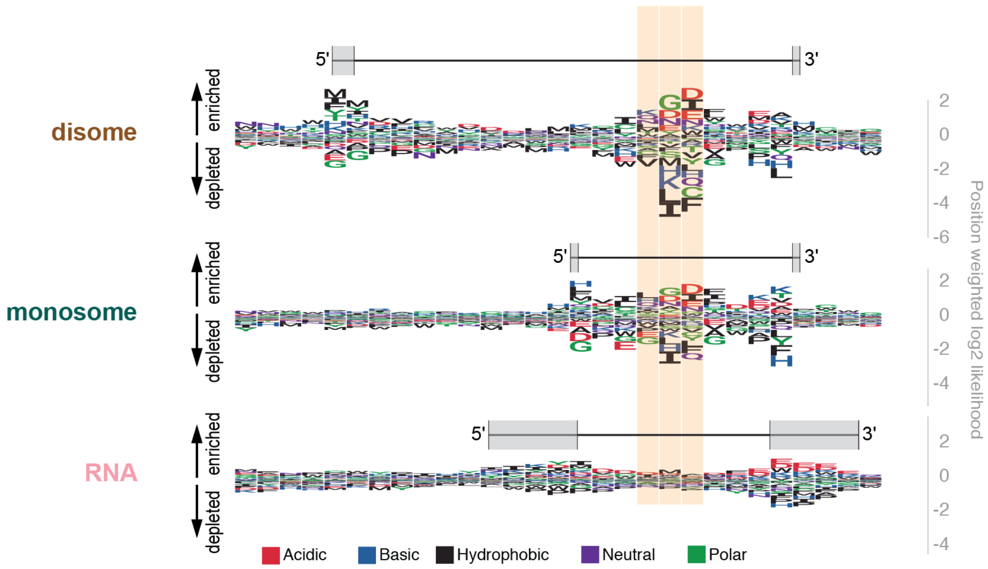

**Supplemental Figure S12. Analysis of disome sites at and around junctions between structured and unstructured regions.**

**(A - C)** On the left, same plots in Figure 4E were redrawn to facilitate comparison. Data from Figure 4E were analyzed within a +/- 30 codon window around positions that corresponded to junctions between structural regions, such as structured-to-unstructured junction (left plots) or unstructured-to-structured (right plots). All junction regions were aligned, such that first codon at the junction was labeled as 0 on the x-axis (codon positions). Average densities (y-axis) of disome (A), monosome (B) footprints or total RNA reads (C) at each position within the window were plotted (red dots). Data from randomized peaks ($N = 10000$) were shown with box-and-whiskers at each codon position.

**(D - F)** Same as (A - C), but the structural configuration was reversed as a control: unstructured - structured - unstructured. In these kind of regions, a decrease in the middle structured section was observed, consistent with Figure 4E and (A - C). Analsis was performed for disome (D), monosome footprints (E) and total RNA reads (F).

### Supplemental Figure S12

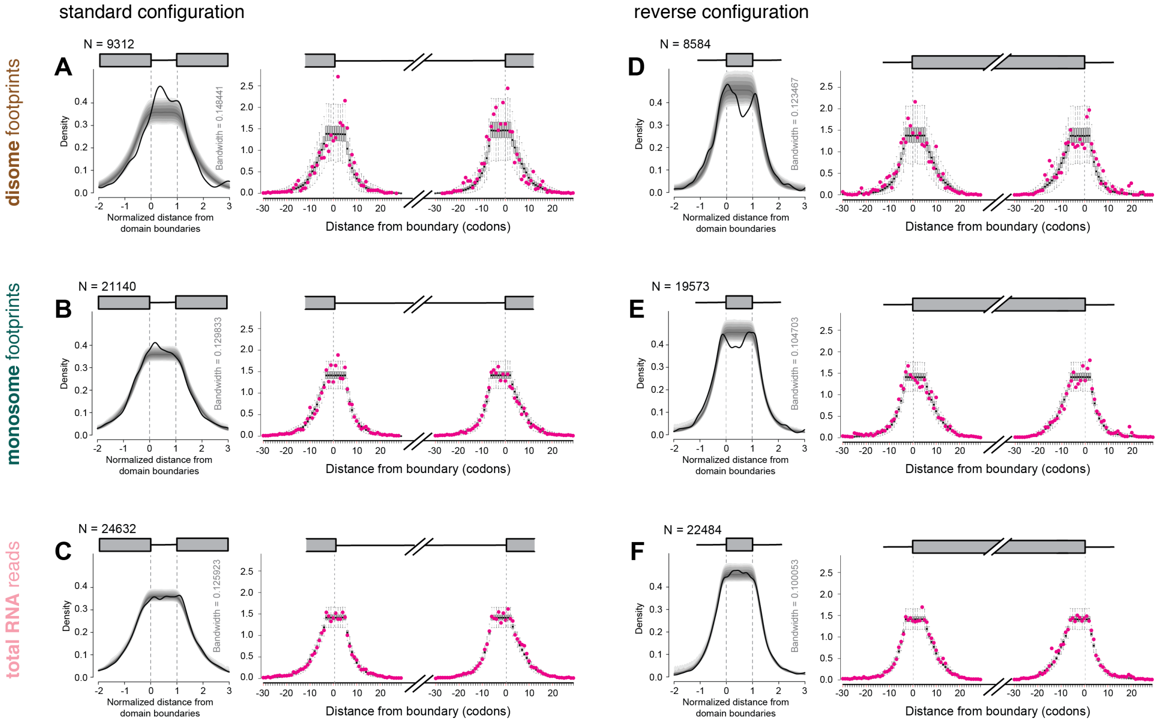

**Supplemental Figure S13. Prominent disome peaks between replicates.**

**(A)** Comparison of ranks of prominent disome peaks identified in individual samples. Prominent disome peaks were identified and ranked as described in Supplemental Table S3, separately for each sample (ZT0, replicate 1 to ZT12, replicate 2 as labeled on top of the heatmap) and for all samples combined. Top 150 peaks from the combined dataset were selected and their ranks were given a color after scaling (bottom). Top 100 peaks from all samples were visualized as a heatmap using these colors or left blank if missing. In general, ranks of disome peaks were similar in individual samples compared to each other and the combined sample indicating reproducibility of prominent peaks.

**(B)** Correlogram of prominent disome peak ranks. Spearman's rank correlation coefficients for all comparisons are given on top-right half. The ellipses represent confidence of correlations: long, narrow ellipses represent high correlations (low correlations would be represented by circular ellipses). Straight Loess lines (red lines inside the ellipses) indicate continuous and uniform correlation of ranks from high to low. Together with **(A)**, these analyses demonstrate robust and reproducible identification of prominent disome peaks.

### Supplemental Figure S13

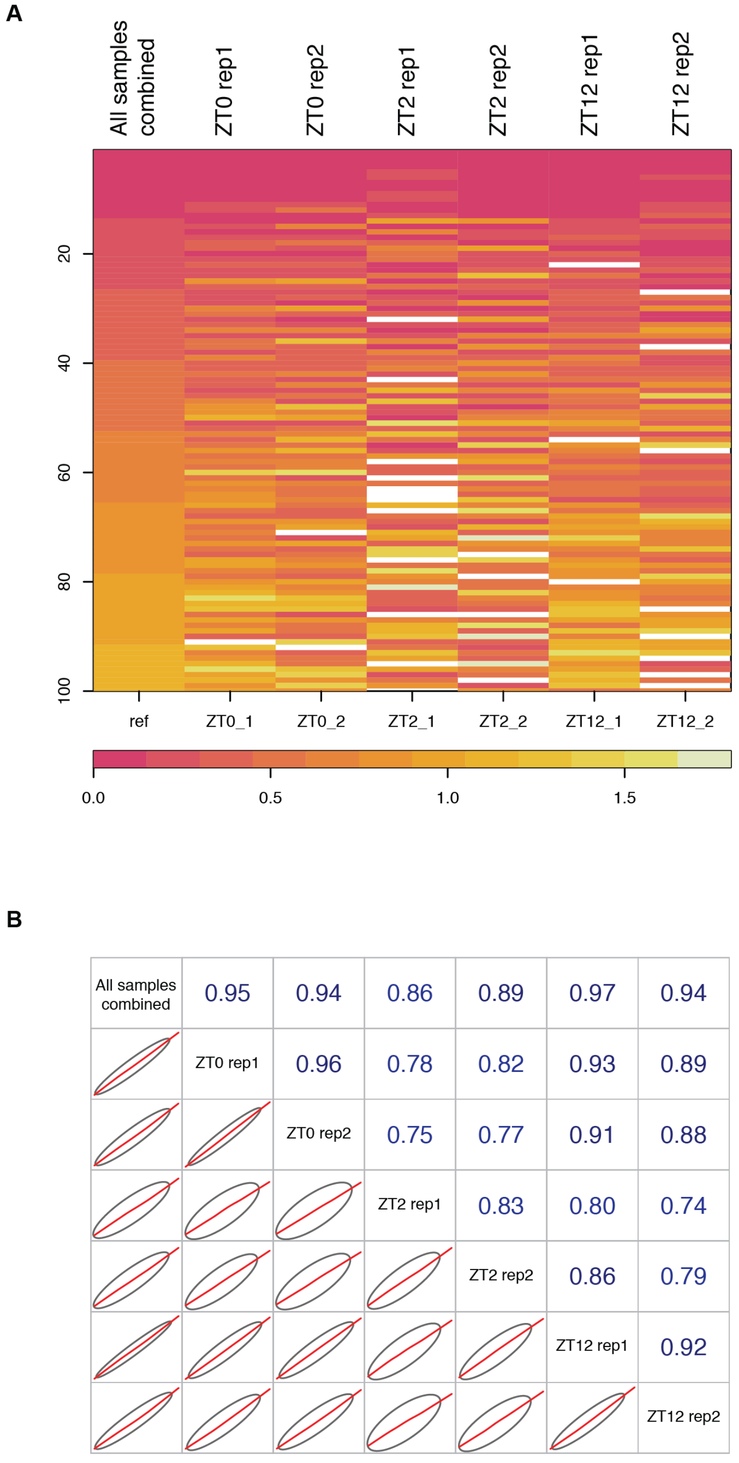

**Supplemental Figure S14. Examples of disome sites in in transcripts/proteins from the list of deterministic sites.**

Each panel is comprised of two parts: the three-dimensional structure of the protein (from mouse or, if not available, a related mammalian species, with the disome site amino acids highlighted in red and with a shaded circle) and transcript plots showing the distribution of disome (brown), monosome (green) and total RNA (pink) signals along the transcript. Shaded areas correspond to the CDS (for UTRs, only the boundaries were plotted). The protein structures have been generated using the following data (PDB ID in parenthesis):

**(A)** Homologue of *Ctsd* from *H. sapiens* (1LYW).
**(B)** Homologue of *Dynlrb1* from *H. sapiens* (6F1Z), corresponding residues at positions 93-94.
**(C)** Homologues of *Fh1* from *H. sapiens* (5UPP), corresponding residues at positions 58-59.
**(D)** *Fth1* structure from *M. musculus* (6S61).
**(E)** Homologue of *Gpd1* from *H. sapiens* (6E8Y).
**(F)** Homologue of *Mrps17* from *S. scrofa* (5AJ3).
**(G)** *Mrsb1* structure from *M. musculus* (2KV1).
**(H)** Homologue of *Nars* from *H. sapiens* (5XIX), corresponding residues at positions 476-477.
**(I)** Homologue of *Ndufb6* from *B. taurus* (5LDW).
**(J)** Homologue of *Nqo2* from *H. sapiens* (1ZX1).
**(K)** Homologue of *Pah* from *R. norvegicus* (1PHZ).
**(L)** *Rdx* structure from *M. musculus* (3X23).
**(M)** Homologue of *Sub1* from *H. sapiens* (4USG).
**(N)** *Sult1d1* structure from *M. musculus* (2ZPT).
**(O)** Homologue of *Tdo2* from *H. sapiens* (4PW8).
**(P)** Homologue of *Tkt* from *H. sapiens* (4KXU).

### Supplemental Figure S14

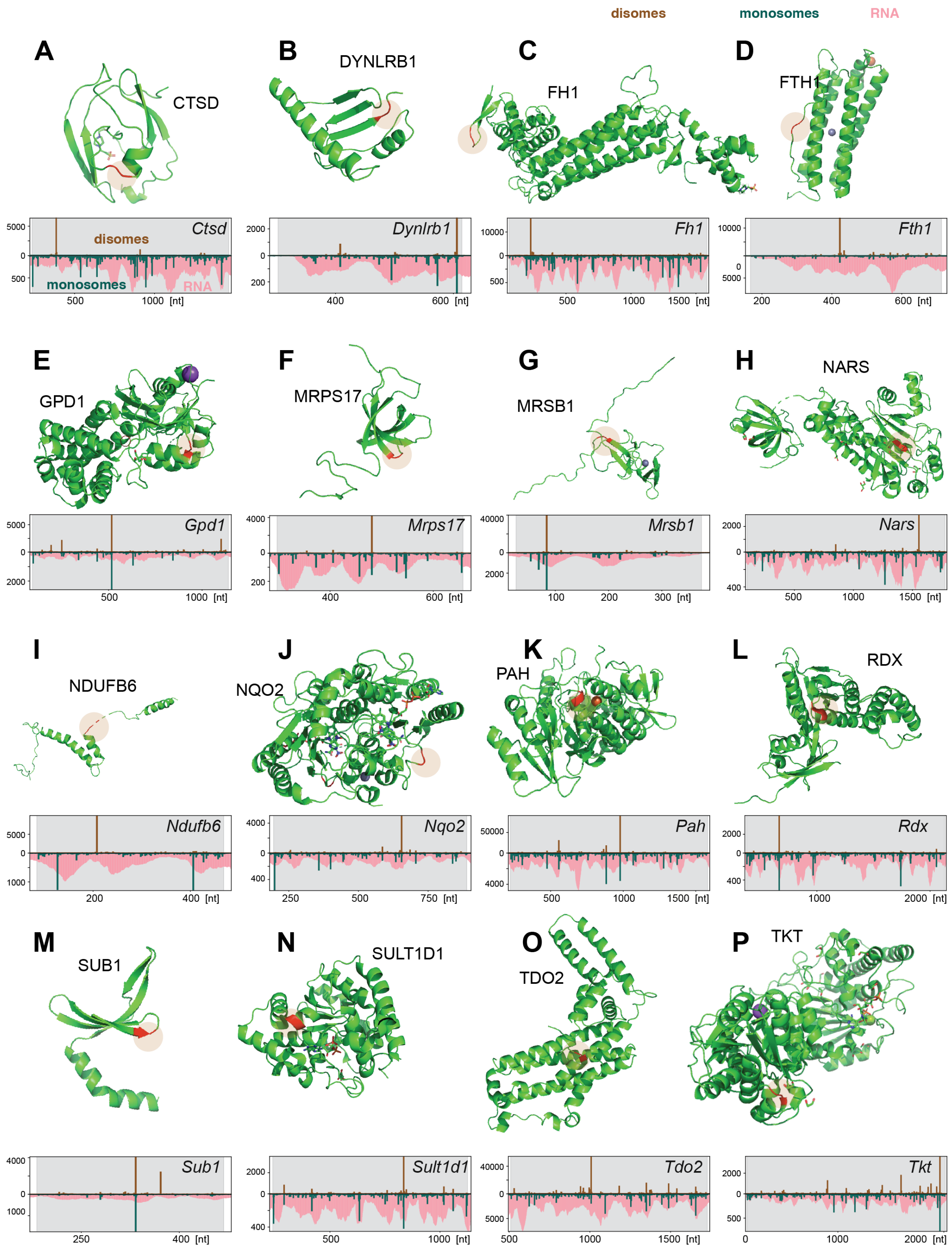

**Supplemental Figure S15. Footprint profiles of transcripts encoding selenocysteine-containing proteins.**

**(A)** Transcript plots of selenocysteine-containing proteins that contained prominent disome peaks (Supplemental Table S3). Plots show the distribution of disome (brown), monosome (green) and total RNA (pink) signals along the transcripts encoding for selenocysteine-containing proteins. Shaded areas correspond to the CDS. Positions of selenocysteine coding UGA codons are marked with red vertical lines and 'U's at the top. In transcript plot of *Selenop*, a cluster of 9 Us is shown as 9xU. Locations of type 1 or type 2 selenocysteine insertion sequences (SECIS) are indicated by red boxes along the x-axis.

**(B)** Same as **(A)** for other selenocysteine-containing proteins in which disome peaks were not among the most prominent ones (were not present in Supplemental Table S3).

### Supplemental Figure S15

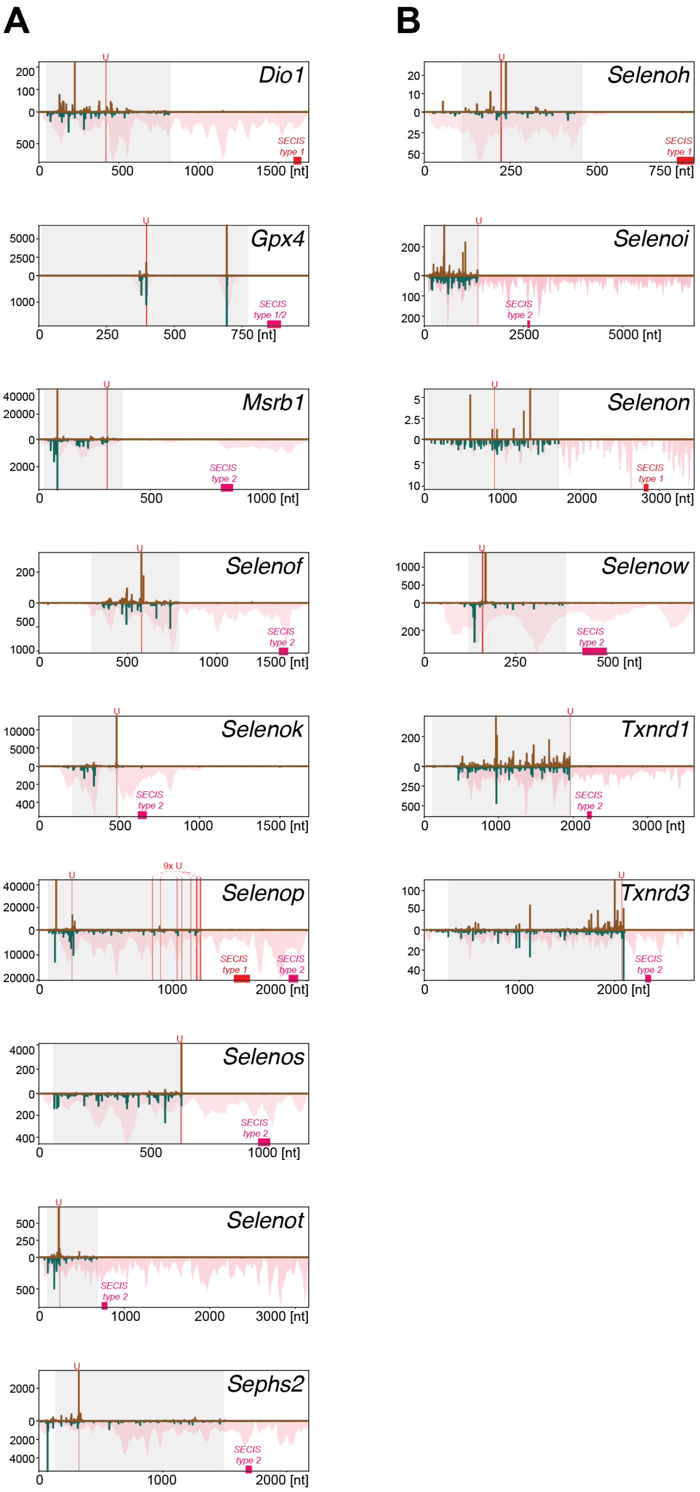

**Supplemental Figure S16. Testing functional importance of disome sites on ribosomal proteins.**

**(A)** Upper: Transcript plot for *Rps5* showing the distribution of normalized disome peaks along the transcript coordinates. Shaded area highlights the CDS and the prominent disome peak at positions 32-33, corresponding to Asp-Ile (DI), is shown with a red arrow. Lower: Same for *Rpl35a* with disome peak at positions 44-45 (Gly-Lys) highlighted. See Figure 2F for further details.

**(B)** Three-dimensional protein structures of human RPS5 (PDB ID: 5VYC) and RPL35A (PDBe ID: 5t2c). The position of the conserved disome site amino acids are highlighted in red.

**(C)** Upper: Schematic of the lentiviral reporter used to probe for the effect of synonymous disome site mutations on steady-state protein abundance. *Rps5* cDNA is fused in-frame to firefly luciferase and transcribed by a bidirectional promoter that also drives the control gene, Renilla luciferase. Lower: Effect of synonymous disome site mutations (mut1-mut5) on Firefly/Renilla ratios, expressed relative to the wild-type reporter which was internally set to 100% in each experiment (N=2-10). Mut2 vs. wild-type: $p=0.008$, Student's t-test. "Dicondon disome occupancy" refers to the percentage of dicodons transcriptome-wide that carry a disome site (see Supplemental Table S2).

#### Supplemental Figure S16

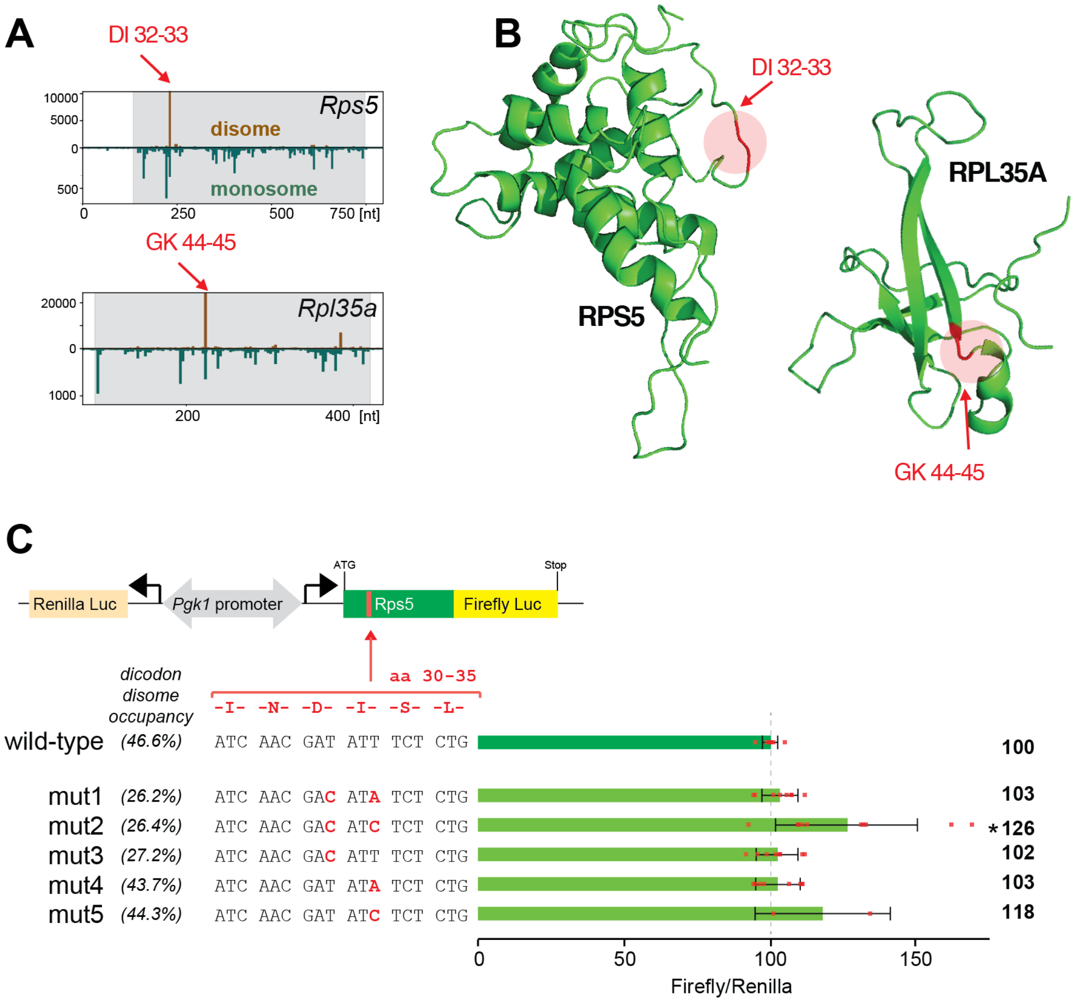

**Supplemental Figure S17. Monosome-Disome peak relationship.**

**(A - C)** On the left, distribution of monosome footprints centered around either monosome peaks **(A)** or disome peaks **(C)** depicted as footprint density heatmaps. Either monosome **(A)** or disome **(C)** peaks were used as anchors to collect all other monosome and disome data (normalized to transcript total) within a 40 codon-wide window spanning from 25 codons upstream to 15 codons downstream of the estimated A-site of the anchoring footprint. Two data matrices were populated by moving the window through all transcripts used in the analysis either using monosome **(A)** peaks as anchors or disomes **(C)**, respectively. Rows in both matrices (measurements within a single window) were sorted by monosome densities at the 0-position corresponding to the A-site of the anchoring peaks and were grouped into approximately 100 groups based on the unique density percentiles of the 0-position monosome densities. Number of observations in each group is given on the histograms on the left side. Within each group, observations were aggregated per position using their trimeans which correspond to the unit rectangles of the heatmaps. Aggregated densities were represented by a 40-level graduated color palette, from blue (low) to red (high) based on their 40-quantiles, as depicted at the bottom of the figure. Genes used in the analyses were filtered to have a single representative transcript model and to have at least 15 disome and 20 monosome footprints in total.

**(B - D)** Distribution of disome footprints centered around either monosome peaks **(B)** or disome peaks **(D)**. Same as **(A - C)**, but all observations were sorted by the disome densities measured at the A-site. One data matrix (anchored/selected by monosome peaks) was used in **(A and B)**, and second one (anchored/selected by disome peaks) was used in **(C and D)**, before sorting. These analyses enabled us to investigate the distribution of monosome and disome densities around other monosome or disome peaks. The presence of phasing (green arrows) monosome footprints that are 10 codon (size of a single ribosome footprint) upstream of either disome or monosome peaks suggests that some disome footprints could be degraded by nucleases. Phasing could only be detected either by investigating regions around disome sites **(C and D)** or by sorting monosome densities by disome densities **(B)**.

### Supplemental Figure S17

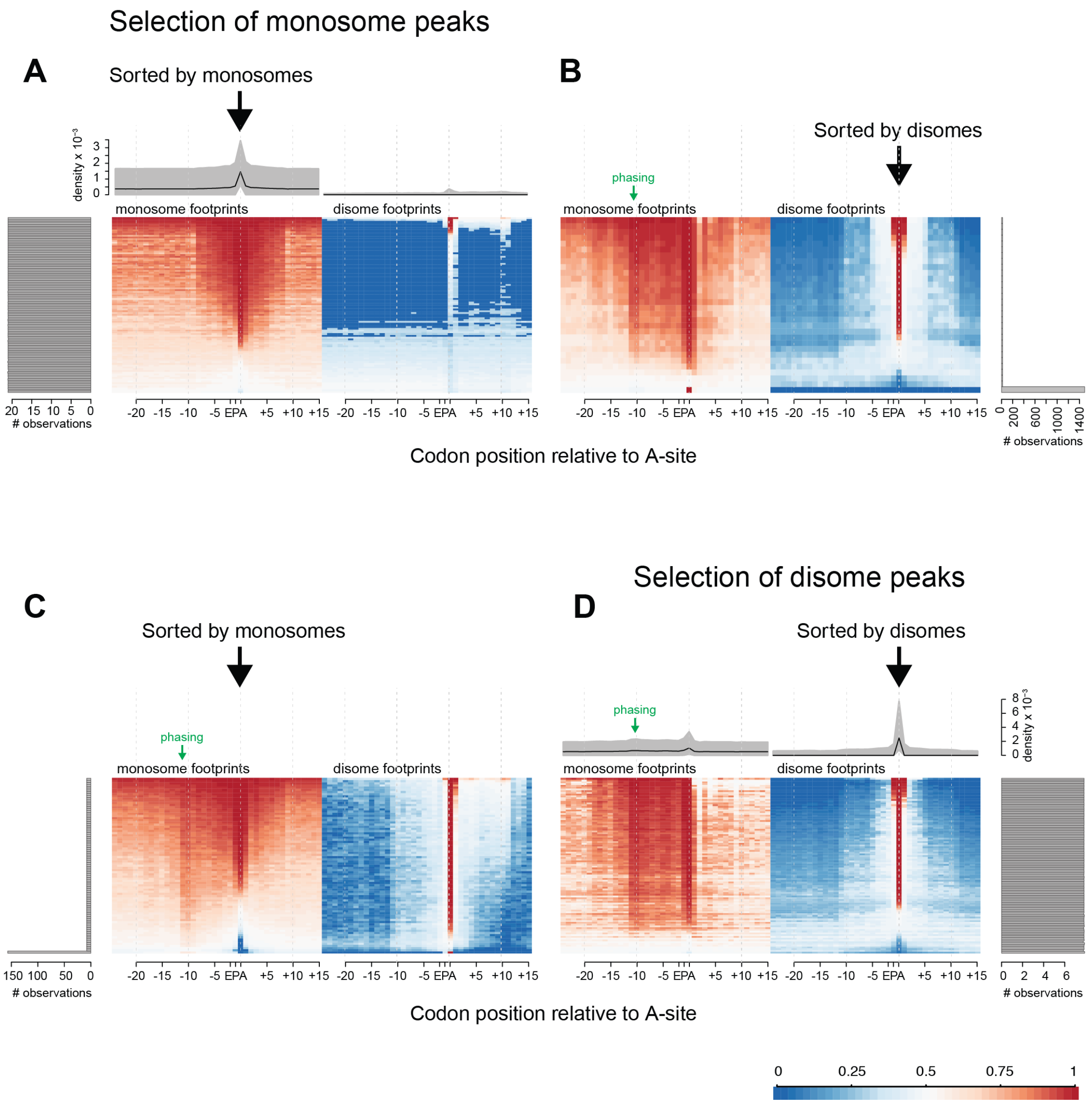

### Supplemental Tables

**Supplemental Table S1:** Sequencing and mapping information.

**Supplemental Table S2:** Amino acid enrichment at disome site (by dicodon).

**Supplemental Table S3:** Transcripts with prominent ('deterministic') disome peaks.

**Supplemental Table S4:** Enrichment analyses for top-200 genes from Supplemental Table S3.

### Supplemental Experimental Procedures

##### Experimental model and subject details

Extracts from 12-week-old male C57BL/6 mice were the same as reported previously (Janich et al., 2015), with experiments approved by the Veterinary Office of the Canon Vaud (authorization VD2376 to DG). NIH3T3 and HEK293FT cells were same cell lines as described in Janich et al. (2015). Culture conditions: DMEM; 10% FCS, 1% penicillin/streptomycin, all from Invitrogen; 37°C; 5% $CO_2$).

##### Experimental methods details

##### Northern blot

The general Northern blot protocol has been described in Gatfield et al. (2009). Briefly, nuclease-digested RNA samples (RNAse I, Ambion) were prepared as described below under "Footprint and library generation (monosome, disome, RNA)". Micrococcal nuclease (for Supplemental Figure S1C) was from New England Biolabs. Per gel, equal amounts of (digested) RNA were loaded in each lane (typically 10-25$\mu$g), electroblotted to Genescreen Plus membrane (NEN), immobilised (UV/baking), and cut into stripes that were independently hybridized with radioactively labelled oligonucleotides: Alb$_{1-22}$ ggagaaaggttac ccacttcat, Alb$_{71-101}$ cgatgggcgatctcactcttgtgtgcttctc, Alb$_{131-165}$ gagatactgggaaaaggcaa tcaggactagg, Alb$_{1099-1120}$ gatcagcaggcatggtgtcatgc, Alb$_{1805-1827}$ ttaggctaaggcgtctttgcat c, Mup7$_{1-21}$ cagcagcagcagcatcttcat, Mup7$_{58-81}$ gttccttcccgtagaactagcttc, Mup7$_{298-320}$ gtattgaatccatcatacgtcac, Mup7$_{563-584}$ tcattctcgggcctggaggcag, Mup7$_{688-708}$ tcagtgag acaggatggaatg. Please note that the lower part of the Northern blot panels shown in Supplemental Figure S1B was also used in our previous publication (Janich et al., 2016).

##### Footprint and library generation (monosome, disome, RNA)

The original mouse liver datasets for monosome footprints and RNA-seq that we used in the current study were the same as those reported in Janich et al. (2015), of which we used the three timepoints, ZT0, 2, 12. As described in the detailed, published protocol (Janich et al., 2015), for each timepoint we had two biological replicates, i.e. a total of 6 independent samples, and each sample was a pool of liver lysates from two mice. The matching disome footprint datasets from the same samples were produced within the framework of the current study. Disome footprints had already been cut simultaneously, and from the same gels, together with the monosome footprints in Janich et al. (2015), yet the disome footprint-containing gel pieces were frozen (-80°C), processed, converted to libraries and analysed only in our current study. Extracts and datasets for the "spike-in experiment" were from independent mice and produced for this study. Mouse ES-cell data were from Tuck et al. (2020).

The general protocol for extract and library preparation has been reported in Janich et al. (2015). Briefly, for extract preparation, freshly harvested mouse livers were homogenized using a motor-driven Teflon homogenizer (5-6 strokes) in 3 volumes of lysis buffer, which consists of polysome buffer (150 mM NaCl, 20 mM Tris-HCl pH7.4, 5 mM $MgCl_2$, 5 mM DTT, 100 $\mu$g·ml$^{-1}$ cycloheximide, complete EDTA-free protease inhibitors (Roche) and 40 U·ml$^{-1}$ RNasin plus (Promega)) supplemented with 1% Triton X-100, 0.5% Sodium deoxycholate. Lysates were incubated for 10 min on ice and cleared by centrifugation at 1000×g, 4°C for 10 min in a tabletop centrifuge. Supernatants were flash-frozen and stored in liquid nitrogen (storage of lysates for several months on liquid nitrogen or at -80°C possible at this step). Lysates were thawed on ice and the OD260 was determined using a Nanodrop spectrophotometer. For each replicate and timepoint, equal amounts of OD260 lysate from two mice were pooled. From the lysate pool, 15×OD260 (for liver: ca. 100 $\mu$l; different samples processed in parallel were adjusted to identical volumes with lysis buffer) were incubated with 650 U RNase I (Ambion) and 5 U Turbo DNase (Ambion) for 45 min at room temperature and gentle agitation. Nuclease digestion was stopped through addition of 8.7 $\mu$l Superasin (Ambion). Sephacryl S-400 HR spin columns (GE Healthcare Life Sciences) were 3 times pre-washed and spun for 1 min at 2000 rpm with 700 $\mu$l polysome buffer (supplemented with 20 U·ml$^{-1}$ Superasin), before applying the lysates on top of the resin and spinning 2 min at 2400 rpm, 4°C. The flow-through was immediately mixed with 1 ml Qiazol, incubated 5 min at room temperature, and the RNAs (containing the ribosome-protected mRNA fragments) were purified using miRNeasy RNA extraction kit (Qiagen) according to the manufacturer's instructions. For the disome datasets complementing the monosome data from Janich et al. (2015), 25 $\mu$g of each RNA obtained after the above purification were separated on a 15% urea-polyacrylamide gel; gel slabs of the desired footprint sizes were cut with the help of size markers (single strand RNA oligonucleotides of 26 nt and 34 nt for monosome footprints, and of 52 nt and 69 nt for disome footprints); RNA was then extracted, and rRNA depletion performed on each of the purified footprint samples, all as described in the original publication (Janich et al., 2015), before proceeding to library preparation. Since then, we have modified our default protocol, which now inverses the steps of PAGE purification and rRNA depletion, i.e. we first deplete rRNA on the full purified RNA sample, and then select footprints by size on PAGE. For the current study, the modified order applied to the spike-in experiment, and to the mESC disome profiling data published in Tuck et al. (2020). Thus, for the modified protocol, 5 $\mu$g of the RNase-digested, purified RNA were used for ribosomal RNA removal with the Ribo-Zero Gold rRNA Removal Kit (MRZG12324 Illumina) according to Illumina's protocol for TruSeq Ribo Profile (RPHMR12126 Illumina). RNA spike-in mix (2 $\mu$l), containing three 30 nt RNA oligonucleotides (sequences: AAU ACCACCCCCAUGAACGCUGCACACACG, AACUACCGACUCAUCCCAUCUUGCCAGU AC, CUAAUACUUACGAACCAGACGAAUCCCUUG) and three 60 nt oligos (AAUACC ACCCCCAUGAACGCUGCACACACGAAUACCACCCCCAUGAACGCUGCACACACG, A

ACUACCGACUCAUCCCAUCUUGCCAGUACAACUACCGACUCAUCCCAUCUUGCCAG UAC, CUAAUACUUACGAACCAGACGAAUCCCUUGCUAAUACUUACGAACCAGACGA AUCCCUUG) at 0.016 fmol$\cdot\mu$l$^{-1}$, was added at this step to the purified, rRNA-depleted RNA samples. Subsequently, monosome and disome sequencing libraries were generated according to Illumina's TruSeq Ribo-Profile protocol with minor modifications. cDNA fragments were separated on a 10% urea-polyacrylamide gel and gel slices between 70-80 nt for monosomes and 97-114 nt for disomes were excised. The PCR-amplified libraries were size selected on an 8% native polyacrylamide gel. Monosome libraries were at $\sim$150 bp and disome libraries at $\sim$180 bp. Parallel RNA-seq libraries were prepared essentially following the Illumina protocol; briefly, after total RNA extraction using miRNeasy RNA Extraction kit (Qiagen), ribosomal RNA was depleted using Ribo-Zero Gold rRNA (Illumina), and sequencing libraries generated from the heat-fragmented RNA as described (Janich et al., 2015). In the spike-in experiemnt, 3 $\mu$l of the same RNA spike mix as above were added to the total RNA after heat fragmentation (during the ice incubation step). All libraries were sequenced in-house on Illumina HiSeq 2500.

**Cloning, lentiviral production, luciferase assays**

For the generation of the Rps5 dual luciferase (Firefly/Renilla) reporter plasmid, *Rps5* CDS (without stop codon) was PCR-ampified from mouse cDNA using oligos Rps5CDS-F, aaaggatccgccaccATGACTGAGTGGGAAGCAGCCACACCAG and Rps5CDS-R, tt tggatccactagtGCGGTTAGACTTGGCCACACGCTCCAGT, digested with BamHI and cloned upstream and inframe of luciferase into BamHI-opened dual luciferase vector prLV1 (Du et al., 2014); this vector is suitable for lentiviral production), and validated by sequencing. Disome site mutants were generated by site-directed mutagenesis with the primers: Rps5mut1-up, GATGACGTGCAGATCAACgacataTCTCTGCAGGATTAC ATTG; Rps5mut1-low CAATGTAATCCTGCAGAGAtatgtcGTTGATCTGCACGTCATC; Rps5mut2-up, GATGACGTGCAGATCAACgacatcTCTCTGCAGGATTACATTG; Rps5m ut2-low CAATGTAATCCTGCAGAGAgatgtcGTTGATCTGCACGTCATC; Rps5mut3-up, GATGACGTGCAGATCAACgacattTCTCTGCAGGATTACATTG; Rps5mut3-low CAAT GTAATCCTGCAGAGAaatgtcGTTGATCTGCACGTCATC; Rps5mut4-up, GATGACGT GCAGATCAACgatataTCTCTGCAGGATTACATTG; Rps5mut4-low CAATGTAATCCT GCAGAGAtatatcGTTGATCTGCACGTCATC; Rps5mut5-up, GATGACGTGCAGATCA ACgatatcTCTCTGCAGGATTACATTG; Rps5mut5-low CAATGTAATCCTGCAGAGAg atatcGTTGATCTGCACGTCATC. All mutants were verified by sequencing.

Plasmids were used to produce lentiviral particles in HEK293FT cells with envelope pMD2.G and packaging psPAX2 plasmids, and viral transduction of NIH3T3 cells, were performed following published protocols (Salmon and Trono, 2007). 1-2 weeks after lentiviral transduction, cells were lysed in passive lysis buffer and luciferase activities were quantified using DualGlo luciferase assay system and a GloMax 96 Microplateluminometer (all from Promega). Firefly/Renilla luciferase (FL/RL) of the Rps5 wt plasmid were internally set to 100% in each experiment, and mutant FL/RL ratios expressed relative to wt.

#### Computational methods details

##### Preprocessing of Sequencing Reads

Initial quality assessment of the sequencing reads was conducted based on Illumina pipeline's (v1.82) preliminary quality values such as the percentage of clusters passed filtering (%PF clusters) and the mean quality score (PF clusters). Adapter sequences were removed with the `cutadapt` utility (Martin, 2011) and following arguments: `-a AGATCGGAAGAGCACACGTCTGAACTCCAGTC`[illegible] `--match-read-wildcards`. Trimmed read sequences were filtered by their size using an in-house Python script to conform the following inclusive ranges: [45,70] for disome footprints, [26,35] for monosome footprints, and [21,70] for total RNA reads. Finally, the reads were filtered for quality using the `fastq_quality_filter` tool from the FASTX-toolkit (http://hannonlab.cshl.edu/fastx_toolkit/) with the following arguments: `-Q33 -q 30 -p 80`.

##### Mapping of Footprints to Mouse Genome

A similar sequential mapping strategy was adapted as described in Janich et al. (2015). The preprocessed insert sequences were mapped sequentially to following databases: mouse rRNA, human rRNA, mt-tRNA, mouse tRNA, mouse cDNA from Ensembl mouse database release 91 Flicek et al. (2013) and, finally, mouse genomic sequences (Genome Reference Consortium GRCm38.p2). With the exception of the final mapping against genomic sequences, `bowtie` version 2.3.0 Langmead and Salzberg (2012) was used with the following parameters:

```
-p 2 -L 15 -k 20 --no-unal
```

After each alignment, unmapped reads were used in the succeeding mapping. For each sequence, only valid alignments with maximum alignment scores were kept. For further analysis, only alignments against mouse cDNA were used, unless specifically stated otherwise.

In parallel to the sequential mapping strategy, preprocessed total RNA sequences were also directly aligned against the mouse genome (GRCm38.p2). Alignments against genome sequence databases were performed using the `STAR` mapper version 2.5.3a (Dobin et al., 2012) with the following parameters:

```
--runThreadN 6 --genomeDir=mouse/star/Mmusculus.GRCm38.91
--readFilesCommand zcat --genomeLoad LoadAndKeep
--outSAMtype BAM SortedByCoordinate Unsorted
--alignSJDBoverhangMin 1 --alignIntronMax 1000000
--outFilterType BySJout --alignSJoverhangMin 8
--limitBAMsortRAM 15000000000
```

The output of this alignment was then processed with StringTie version 1.3.3b (Pertea et al., 2015) to estimate the number of fragments per kilobase of exon per million fragments mapped (FPKM) for each transcript (Ensembl mouse database release 91), with the following parameters:

```
-p 8 -G Mmusculus.GRCm38.91.gtf -A gene_abund.tab
-C cov_refs.gtf -B -e
```

The outputs were parsed with an in-house Python script to identify transcripts which had an FPKM $>0.2$ and an isoform abundance fraction $>0.05$ in at least 2 samples. A database of expressed transcripts ($N_{genes} = 19508$, $N_{transcripts} = 24927$) was used in further analysis. Among those, genes that were estimated to have a single expressed isoform were annotated as `single transcript genes` (N = 9711). For genes with multiple transcript isoforms, the transcipt, whose exons were inclusive of all others, was used whenever possible (N = 548).

**Quantification of mRNA and Footprint Abundances**

Abundance of total RNA reads and monosome or disome footprints was estimated per locus as described in Janich et al. (2015). Separate counts were obtained for whole gene, UTRs and CDS. Only reads that were mapped uniquely to a single gene and only to transcripts that were identified to be expressed (see Mapping of Footprints to Mouse Genome) were used. Exclusively for the analysis of ribosomal proteins (Figure S3D), this criterion was slightly relaxed to also include multireads that were mapping to a single protein coding locus. Transcripts which did not have at least 10 counts in at least one third of the samples were excluded. For all further analysis, reads that mapped to CDS regions were used, unless stated otherwise. A total of 8626 loci had above threshold read counts within the CDS for all read types: total RNA, monosome and disome.

Read counts of total RNA and footprints were normalized with upper quantile method of R package `edgeR` v3.16.5 Robinson et al. (2010). For increased comparability between datasets, RPKM values were calculated as the number of reads per 1000 bases per geometric mean of normalized read counts per million. Genes that had an average total RNA RPKM $>5$ were designated as `robustly expressed`. Combined with the single transcript genes (see Mapping of Footprints to Mouse Genome), robustly expressed single transcript genes (N = 6007) were used for analyses where inclusion of genes with multiple expressed isoforms was not possible (e.g meta-transcript analysis). Normalized footprint densities were calculated as the $log_2$-ratio of footprint-RPKM to total RNA-RPKM per gene, for disomes and monosomes. For the latter, this quantity is also called translational efficiency (TE). In mouse liver, TEs were shown to be stable over time-points around the day (Janich et al. (2015); disome densities were similarly stable between the samples (ZT0, ZT2, and ZT12) and therefore treated as replicates, unless stated otherwise.

#### Spike-in Normalization and Global Quantification of Ribosomes Retained in Disomes

Random 30 and 60 nt long RNA oligonucleotide sequences were designed following these criteria: (i) have a GC % similar to that of mouse liver translatome (mean was 52.05, 5% and 95% were 42.2 and 62.6, respectively), (ii) should be void of potential hairpin structures and self-dimerization (using `ViennaRNA` package 2.3.5, Lorenz et al. (2011)), (iii) should not be highly similar to mouse or Drosophila transcriptome and genome, (iv) should not contain certain sequences at the extremities which we were identified as highly biased in our analyses (GG, GC, CC, CG, CA, GA, TG, AC), (v) should not contain stop codons and (vi) 60mers were designed as 2 x 30mer repeats. Out of 35 possible candidates, 3 sets with different GC% were selected: 43, 50 and 56. Spike reads were mapped and processed similarly to all other reads. To avoid counting degradation products of the 60mers as 30mers, we devised a two-step counting algorithm. First, spike read distributions were inspected on total RNA reads to assess possible degradation and define proper size limits. The GC56 spike was eliminated from further analysis due to fragmentation; for others [24,31] and [45,60] inclusive size filters were used for 30- and 60mers, respectively. In addition to the size filtering, the presence/absence of the junction of the 2 x 30mer repeats were identified for all spike reads. 30mers were included if they did not have a junction, and 60mers only if they did. Spike counts were first normalized for library size with upper-quantile method and spike-in normalization factors were calculated as 60mer / 30mer ratios per sample to correct the experimental biases between the disome and monosome counts. The spike-in normalization factors were nearly identical for triplicate biological replicates (mean = 2.495, SD = 0.028). The spike-normalized counts of disomes and monosomes were then used to estimate the percentage of ribosomes that were identified within disomes to the whole, taking into account that each disome represented two ribosomes.

#### Observed-to-Expected Ratios For Proximal Sequence Features

The calculation of observed-to-expected ratios for sequence features proximal to footprint sites was performed following the principles of Ribo-seq Unit Step Transformation method (O′Connor et al., 2016). First, footprint (or total RNA read) densities were normalized to the sum of transcript densities, then a Heaviside step function was applied to individual features (codon, amino acid, 6mer, dipeptide, charge, secondary structure, PhyloP conservation categories) along each CDS, such that a feature at a position was given a score of 1 or 0 depending on whether the footprint density at that position exceeded the average of the corresponding CDS. A margin of 30 nt were excluded from each end of the CDS. Then, a typically 50-codon wide window (80-codon wide for certain analysis such as charge in Figure 4A), was moved along the CDS regions at 3 nt steps, except for analyses that required single nt resolution. Window positions were labeled relative to RUST scores, 0-position marking the score. The scores were either not offset (5') or A-site offset (see Estimation of A-site Positions). At each iteration, position specific occurrence of features was counted and associated with if there was a RUST score in that window. Present, observed and expected values of each feature at each window position were calculated as sums over all windows. When necessary, Kullback-Leibler divergence scores were calculated using the observed-to-expected ratios of all features (O′Connor et al., 2016). Enrichment was calculated as the observed-to-present ratio normalized to expected. All analyses were performed with in-house Python (creation of data matrices) and R software (visualization and statistical analysis). Features that were based on (discrete) sequence information (nucleotide or amino acid sequence) were created simply using the letters of such sequences in different word sizes (such as CCT or proline for single; CCTCCA or proline-proline for two-word). Other discrete data, such as secondary structure, were also analyzed similarly. Features that were based on continuous numeric data, were first stratified into discrete levels. For example, PhyloP conservation scores were grouped into three categories: neutral [-3, 3), conserved [3, 5) and highly conserved [5, ). Visualization of complex RUST ratios was facilitated using $log_2$ transformed position specific enrichment matrices with the `ggseqlogo` package for R (Wagih, 2017) and converting them sequence logos. For these analyses, samples were combined unless it is stated otherwise.

**Estimation of A-site Positions**

The A-sites of the monosomes (RPF) were calculated identically as described in Janich et al. (2015). For disomes, for initial analyses we used a similar approach to estimate the A-site of the upstream ribosome in the disome pair as 15 nt from the 5' end of the footprints. This approach was suitable for exploratory analyses (e.g meta-transcript analysis) for facilitating the comparability to monosome results. In other analyses, we used an empirical method to estimate the A-site of the leading (downstream) ribosome within the disome pair. In order to infer the optimum offsets for different lengths of footprints, we first split the disome footprints by their size, from 55 to 64 nt. Wihtin each size group, footprints were further split into 3 classes based on their open reading frame relative to that of the main CDS. For each group, position-specific (relative to their 5' ends at nucleotide resolution) information content matrices were calculated using the Kullback-Leibler divergence scores (O′Connor et al., 2016) of observed-to-expected ratios of codon analysis (see Calculation of Expected-to-observed Ratios For Proximal Sequence Features). For combinations of footprint size and open reading frame, where the position of PA sites could be identified as highest information positions (with 2 peaks 3 nt apart from each other) around 40 - 50 nt downstream of the 5' ends of the footprints, exact offsets were calculated as the distance of the deduced A-site from the 5' end. Following offsets for 58, 59, 60, 62 and 63 nt long disome footprints on different open reading frames were used, respectively: [45, 44, 43], [45, 44, 46], [45, 44, 46], [48, 47, 46], [48, 47, 49]. Total RNA reads were offset with different methods to be consistent with the dataset they were being compared to: by their center (general), +15 (when compared to monosomes, also selecting a similar size range of 26-35 nt) or disome offsetting (selecting a size range of 58-63).

##### Meta-transcript Analysis

Meta-transcript analyses were performed on robustly expressed single transcript genes that had a CDS region larger than 400 nt and UTRs of larger than 180 nt (N = 4994). Firstly, footprint positions were determined with appropriate A-site estimation (see Estimation of A-site Positions), then footprint counts were normalized to the total number of footprints per transcript. Mean normalized footprint densities were plotted for the first or last 400 nt of CDS plus a small region from the adjacent UTRs. For analysis of signal peptide (SP) genes, transcripts were annotated as SP or no-SP based on the Signalp protein feature from Ensembl Database v91. To calculate the probability densities of length normalized proportions of footprints within the first 75 codons and the rest of CDS, for each transcript, footprints within each portion were counted separately and normalized to library size as usual and in addition to the size of their respective counting region. Then length-normalized counts per region (first 75 codons vs rest of CDS) were expressed as a proportion to their sums, so that when footprints have similar densities between the two regions, normalized proportions would be around 0.5. The analysis was repeated for SP and no-SP genes using either disome or monosome footprints. For these analyses, samples were combined unless it is stated otherwise.

##### Visualization of Footprint Densities Across Transcripts in Relation to CDS Length

A position-specific density matrix of disome footprint densities (based on A-site estimates) was calculated from single transcript genes that contained at least 10 disome footprints (N = 9454, combined samples). Transcripts were ranked by their CDS length. To account for differences in expression levels, footprint densities were normalized to the footprint sum for each transcript. The density matrix was visualized as heatmaps by aligning transcript positions relative to the start or stop codon of each transcript. Analysis was done separately for the subset of signal peptide encoding transcripts (N=1116). Heatmap color shades were mapped to the intervals based on the 0.25, 0.5, 0.75, 0.8, 0.95 quantiles of positive disome densities.

##### Analysis of Footprint Densities in Relation to Peptide Secondary Structures

An in-house Python script was used to extract annotated secondary structures of peptides (UniProt Database (UniProt Consortium, 2018) release-2018_06) mapping them to the corresponding codon positions along CDS. This information is either used for analysis of observed-to-expected ratios (see Observed-to-Expected Ratios For Proximal Sequence Features), or studying the distribution of footprint (disome or monosome) densities across regions with pre-defined structural compositions such as structured-unstructured-structured (s-u-s). To this end, we have extracted coordinates of regions that included a stretch of structured (min. 3 aa, up to 30th position), followed by an unstructured stretch (6 to 30 aa), and finally concluded with a structured stretch (min. 3 aa, up to 30th position) or the reverse of this configuration (u-s-u) with similar size restrictions. Positions of normalized footprint peaks (normalized to transcript's mean footprint count) across regions were scaled to the length of the middle portion (unstructured portion in the case of s-u-s) and centered to its start, such that the start and the end of the middle region would correspond to 0 and 1, respectively. Distribution of footprint densities across such regions was analyzed by kernel density estimates which were weighted with normalized footprint peak densities. Significance of density probability functions were evaluated with randomized sampling. For each transcript, keeping the structures identical, peaks were randomly shuffled (N = 10000) and confidence intervals for the kernel densities were calculated. Total RNA reads were similarly analyzed as a control.

**Analysis of Evolutionary Conservation at Disome Sites**

Evolutionary conservation of sites were evaluated using the PhyloP scores (Pollard et al., 2010), that were computed from alignments of 59 vertebrate genomes to the mouse genome (phyloP60way, mm10.60way.phyloP60way.bw file) and the euarchontoglire subset (mm10.60way.phyloP60wayEuarchontoglire.bw file), which included 21 of the 60 vertebrate species within the Supraprimates (Euarchontoglires) clade. Data were retrievable from the "Conservation" tracks in the UCSC Genome Browser (Haeussler et al., 2018) or from http://hgdownload.soe.ucsc.edu/goldenPath/mm10/phyloP60way/. When required, the 60-way vertebrate PhyloP scores were stratified in 5 levels : highly accelerated [, -5), accelerated [-5, -3), neutral [-3, 3), conserved [3, 5), highly conserved [5, ), of which the presence of the first two were negligible within CDS regions. For logistic regression analysis, mean PhyloP conservation scores were calculated for dicodons using only first and second codon positions. Mean PhyloP scores were then dichotomized as low and high based on the following cutoffs: 1.3 and 5 for euarchontoglire subset and 60-way vertebrates, respectively. These thresholds discriminate between the modes of the bimodal distribution of PhyloP scores and lie approximately around 60 percentile. All possible dicodons were extracted from robustly expressed single transcripts (N = 6001, see section Quantification of mRNA and Footprint Abundances). Presence of disome peaks was defined as having an A-site density at the second codon that was larger than the mean transcript density (same Heaviside step function described in Observed-to-Expected Ratios For Proximal Sequence Features section). The binary outcome of low/high mean dicodon PhyloP score was regressed against scaled mean transcript PhyloP score, the dipeptide encoded by the dicodon (20 x 20 aminoacids excluding selenocysteine = 400 levels) and the binary disome status (present/absent) of the dicodon using a logistic model. The generalized linear regression was fit using the *glm* function of R with the 'binomial' family and the 'logit' link-function.

**Mapping of Disome Amino Acids onto Protein Three-dimensional Structures**

Structure models for target proteins were downloaded from the Protein Data Bank (PDB, http://www.rcsb.org/ ) or Protein Data Bank Europe (PDBe, www.ebi.ac.uk/pdbe/ ). Image rendering was performed with PyMol (DeLano, 2002). For proteins where no murine structure was available, data from other related mammals (mostly from *H. sapiens*) was used instead. The identification of the residues with high disome signal in the non-mouse protein was perform by manual comparison of the two protein sequences (the original for mouse and the target from the other mammal).

**Functional Enrichment Analysis of Genes with Prominent Disome Peaks**

Deterministic disome peaks were defined as prominent peaks that were not necessarily a result of high levels translational activity. To identify such peaks, library size normalized disome peaks (normalized peak count $> 5$) along each transcript were normalized to the mean monosome count of that transcripts, treating all samples as replicates and combining them. To avoid very noisy peaks, transcripts that had a mean monosome count fewer than 5 were excluded. For each transcript up to 5 peaks (defined by their codon position on the transcript) that had the highest monosome-normalized scores were collected. Finally, peaks were sorted in descending order of the normalized scores (Supplemental Table S3). To assess the reproducibility of this approach, the same analysis was also performed on individual samples. The correlation between the ranks of prominent peaks within the resulting sorted lists was then analysed using Spearman's rank correlation analysis. Correlograms were generated using the *corrgram* package in R. Top 200 genes (identified by their Ensembl IDs) from the list (combined samples, Supplemental Table S3) were subsequently submitted to functional enrichment analysis using the g:GOSt tool of web-based g:Profiler software and database platform (Raudvere et al., 2019). Statistically significantly enriched terms within three Gene Ontology (GO) groups - molecular function, cellular component and biological process - were identified (Supplemental Table S4). False discovery was controlled by the default method, g:SCS, to an experiment-wide threshold of a=0.05. As a background, list of all genes identified to have above threshold levels of total RNA, monosome and disome reads in the current study was used (N = 8626).

#### References

DeLano, W.L., 2002. Pymol: An open-source molecular graphics tool. CCP4 Newsletter on protein crystallography 40, 82–92.

Dobin, A., Davis, C.A., Schlesinger, F., Drenkow, J., Zaleski, C., Jha, S., Batut, P., Chaisson, M., Gingeras, T.R., 2012. STAR: ultrafast universal RNA-seq aligner. Bioinformatics 29, 15–21.

Du, N.H., Arpat, A.B., De Matos, M., Gatfield, D., 2014. Micrornas shape circadian hepatic gene expression on a transcriptome-wide scale. eLife 3, e02510.

Flicek, P., Ahmed, I., Amode, M.R., Barrell, D., Beal, K., Brent, S., Carvalho-Silva, D., Clapham, P., Coates, G., Fairley, S., Fitzgerald, S., Gil, L., Garcia-Giron, C., Gordon, L., Hourlier, T., Hunt, S., Juettemann, T., Kaehaeri, A.K., Keenan, S., Komorowska, M., Kulesha, E., Longden, I., Maurel, T., McLaren, W.M., Muffato, M., Nag, R., Overduin, B., Pignatelli, M., Pritchard, B., Pritchard, E., Riat, H.S., Ritchie, G.R.S., Ruffier, M., Schuster, M., Sheppard, D., Sobral, D., Taylor, K., Thormann, A., Trevanion, S., White, S., Wilder, S.P., Aken, B.L., Birney, E., Cunningham, F., Dunham, I., Harrow, J., Herrero, J., Hubbard, T.J.P., Johnson, N., Kinsella, R., Parker, A., Spudich, G., Yates, A., Zadissa, A., Searle, S.M.J., 2013. Ensembl 2013. Nucleic Acids Research 41, D48–D55.

Gatfield, D., Le Martelot, G., Vejnar, C.E., Gerlach, D., Schaad, O., Fleury-Olela, F., Ruskeepää, A.L., Oresic, M., Esau, C.C., Zdobnov, E.M., Schibler, U., 2009. Integration of microrna mir-122 in hepatic circadian gene expression. Genes & Development 23, 1313–1326.

Haeussler, M., Zweig, A.S., Tyner, C., Speir, M.L., Rosenbloom, K.R., Raney, B.J., Lee, C.M., Lee, B.T., Hinrichs, A.S., Gonzalez, J.N., Gibson, D., Diekhans, M., Clawson, H., Casper, J., Barber, G.P., Haussler, D., Kuhn, R.M., Kent, W., 2018. The UCSC Genome Browser database: 2019 update. Nucleic Acids Research 47, D853–D858.

Ingolia, N., Lareau, L., Weissman, J., 2011. Ribosome profiling of mouse embryonic stem cells reveals the complexity and dynamics of mammalian proteomes. Cell 147, 789–802.

Janich, P., Arpat, A., Castelo-Szekely, V., Lopes, M., Gatfield, D., 2015. Ribosome profiling reveals the rhythmic liver translatome and circadian clock regulation by upstream open reading frames. Genome Research 25, 1848–1859.

Janich, P., Arpat, A.B., Castelo-Szekely, V., Gatfield, D., 2016. Analyzing the temporal regulation of translation efficiency in mouse liver. Genomics Data 8, 41 – 44.

Langmead, B., Salzberg, S.L., 2012. Fast gapped-read alignment with bowtie 2. Nature Methods 9, 357–U54.

Lorenz, R., Bernhart, S.H., Höner zu Siederdissen, C., Tafer, H., Flamm, C., Stadler, P.F., Hofacker, I.L., 2011. Viennarna package 2.0. Algorithms for Molecular Biology 6, 26.

Martin, M., 2011. Cutadapt removes adapter sequences from high-throughput sequencing reads. EMBnet.journal 17, 10–12.

O'Connor, P.B.F., Andreev, D.E., Baranov, P.V., 2016. Comparative survey of the relative impact of mrna features on local ribosome profiling read density. Nature Communications 7, 12915.

Pertea, M., Pertea, G., Antonescu, C., Chang, T., Mendell, J., Salzberg, S., 2015. Stringtie enables improved reconstruction of a transcriptome from rna-seq reads. Biotechnology 33, 290–295.

Pollard, K.S., Hubisz, M.J., Rosenbloom, K.R., Siepel, A., 2010. Detection of nonneutral substitution rates on mammalian phylogenies. Genome research 20, 110–121.

Raudvere, U., Kolberg, L., Kuzmin, I., Arak, T., Adler, P., Peterson, H., Vilo, J., 2019. g:Profiler: a web server for functional enrichment analysis and conversions of gene lists (2019 update). Nucleic Acids Research 47, W191–W198.

Robinson, M.D., McCarthy, D.J., Smyth, G.K., 2010. edger: a bioconductor package for differential expression analysis of digital gene expression data. Bioinformatics 26.

Salmon, P., Trono, D., 2007. Production and titration of lentiviral vectors. Curr Protoc Hum Genet Chapter 12, Unit 12.10.

Tuck, A.C., Rankova, A., Arpat, A.B., Liechti, L.A., Hess, D., Iesmantavicius, V., Castelo-Szekely, V., Gatfield, D., Bühler, M., 2020. Mammalian rna decay pathways are highly specialized and widely linked to translation. Molecular Cell .

UniProt Consortium, T., 2018. UniProt: the universal protein knowledgebase. Nucleic Acids Research 46, 2699–2699.

Wagih, O., 2017. ggseqlogo: a versatile R package for drawing sequence logos. Bioinformatics 33, 3645–3647.
